# CD98 is critical for a conserved inflammatory response to diverse injury stimuli relevant to IPF exacerbations and COVID pneumonitis

**DOI:** 10.1101/2022.08.12.503154

**Authors:** Panayiota Stylianou, Sara Rushwan, Wei Wang, M. Azim Miah, Omeed Darweesh, Alison Mackinnon, Katy M. Roach, Charles J. Hitchman, Oksana Gonchar, Stephen Thorpe, Christopher Harris, Richard Haigh, David F Richards, Vladimir Snetkov, Jessica Beasley, Simon J Cleary, Michael Barer, Jeremy PT Ward, Claire Rooney, Frank McCaughan, Peter Bradding, Richard Beale, Martin M Knight, Tariq Sethi, Bibek Gooptu

**Author notes:** Contributed equally; each entitled to list themselves as first named author on list with appropriate recognition of others as equal contributors (Panayiota Stylianou, Sara Rushwan, Wei Wang).

## Abstract

Progressive fibrosing interstitial lung diseases (PFILDs) cause substantial morbidity and mortality. Antifibrotic agents slow progression, but most of the clinical need remains unmet. The archetypal PFILD is idiopathic pulmonary fibrosis (IPF). Chronic progression is driven by transforming growth factor (TGF-)β1 signalling. It is punctuated by inflammatory flares known as acute exacerbations (AE-IPF), which are associated with accelerated decline and high mortality. We hypothesized that acute injury responses underlying exacerbations and the mechanisms of chronic fibrosis overlap at the molecular level, via a cell surface assembly nucleated by galectin-3 that we term the ‘gal-3-fibrosome’. We focused upon a putative pro-inflammatory galectin-3 ligand, the CD98:integrin complex. Our data indicate CD98 and β1-integrin co-localise with galectin-3 within epithelial cells in IPF lung tissue, and within 40 nm in human lung tissue treated with TGF-β1 compared to controls. CD98 is required for interleukin (IL-)6 and IL-8 responses to biochemical and biophysical conditions mimicking stimuli of AE-IPF *in vivo*, *ex vivo* and in cells, and for an interstitial neutrophilic response in a mouse model. We demonstrate this pathway progresses via intracellular influx of Ca^2+^ mediated by TRPV4, and NF-κB activation, operating in positive feedback. Lastly we show the CD98- and galectin-3-dependence of IL-6 and IL-8 responses to the SARS-CoV-2 spike protein receptor binding domain and the conservation of this response pattern between lung epithelial cells and monocyte-derived macrophages. Taken together our findings identify CD98 as a key mediator of both pro-fibrotic and acute inflammatory responses in the lung with relevance to AE- and chronic progression of IPF, and the priming of fibrotic lungs for acute inflammatory responses. They similarly implicate CD98 and galectin-3 as mediators of COVID pneumonitis and worse outcomes in ILD patients with COVID.

## Introduction

Observations in idiopathic pulmonary fibrosis (IPF) demonstrate that innate injury responses can drive the progressive fibrosis phenotype associated with major unmet clinical need (1–3). A similar phenotype can ensue in other interstitial lung diseases (ILDs) with primarily auto-immune or hypersensitivity aetiology (4–9), and responds similarly to small molecule therapies that slow progression but cannot halt it (10–13). These findings support a common molecular pathway that may be accessed from different inflammatory contexts, but is most purely observed in IPF. Sustained injury responses from alveolar epithelial cells (14), and interactions with the mesenchymal population of myofibroblasts (15) appear important. Once fibrosis is established in one region of lung tissue, increased mechanical strain in neighbouring parenchyma through the ventilatory cycle may propagate it (16, 17).

Acute exacerbations of IPF (AE-IPF) are associated with rapid decline and death in IPF (18–21). Patients on the antifibrotic drug nintedanib may have reduced incidence of AE-IPF (22), supporting cross-talk between acute inflammatory responses and chronic fibrosis, but no specific therapies have yet proven effective in their treatment. Repeated sub-acute injury to alveolar epithelial cells is also likely to drive chronic decline in IPF (23, 24), where diffuse alveolar damage can be seen without clinical evidence of an acute exacerbation (25).

We hypothesized that a macromolecular checkpoint complex on the alveolar epithelial cell surface might act as a critical molecular bridge linking acute inflammatory injury and chronic fibrosis responses, allowing cross-talk and modulation. We further hypothesized that the chimaeric protein galectin-3 could nucleate such a multi-protein assembly including the TFG-β receptor, that we term the ‘gal-3-fibrosome’. Galectin-3 contains a C-terminal carbohydrate binding domain and an N-terminal domain capable of protein:protein interactions (26). Oligomerisation is stimulated by glycan binding and involves release of the N-terminal domain from the C-terminal domain (27). This can nucleate an extended lattice that also involves heterotypic interactions with binding partners. Galectin-3 is implicated as a key mediator and/or biomarker of multiple fibrotic processes, including in the lung where its protein:glycoprotein interactions appear to be critical in two different models of experimental fibrosis (28). These are blocked by an experimental anti-fibrotic therapy (29, 30). Galectin-3 potentiates TGF-β1 activity at the cell surface (28), likely through stabilisation of the TGF-β receptor (TGF-βR)II protein to which it can bind directly. We postulate that the gal-3-fibrosome may incorporate other proteins relevant to fibrosis and/or inflammation, facilitating cross-talk between TGF-β1 signalling and other acute inflammatory and chronic fibrotic injury response pathways.

The heterodimeric transmembrane protein CD98 is a promising candidate for this. Its constant 70 kDa heavy chain (CD98hc) is predominantly extracellular, with a single transmembrane sequence and a short cytoplasmic tail, and associates with a variable light chain amino-acid transport channel (31). CD98hc binds to multiple integrins, including β3- and, constitutively, β1-integrin (32–35), and is implicated as a binding partner to galectin-3 (36, 37). CD98 is involved with pro-inflammatory pathways in lymphocytes and gastrointestinal epithelia (38, 39). Integrins mediate cross-talk between the extracellular matrix and cellular responses (40, 41). β1- and β3-integrin can bind the latent form of the pro-fibrotic cytokine transforming growth factor (TGF-)β1 via a classic Arg-Gly-Asp (RGD) amino-acid sequence motif. The interaction localizes latent TGF-β1 at the cell surface prior to activation by mechanical stretch (17), other integrins (42), or protease activity (43). β1-integrin forms a transmembrane bridge between extracellular matrix proteins and the cytoskeleton via fibrillar adhesions. It mediates RhoA-, αvβ6-integrin-dependent activation of TGF-β1 in fibrosis models (44–46). Thus CD98:integrin complexes may mediate both acute cytokine responses to biochemical and mechanical stimuli and the pro-fibrotic TGF-β1 response pathway, with cross-talk between these processes. This is illustrated in our initial molecular model (Supp. Fig. 1a) for a simplified gal-3-fibrosome arrangement (galectin-3 multimer represented by dimeric component). Interestingly in the context of COVID pneumonitis, where people with ILD have a 60% increased mortality risk, a K403R mutation that has evolved in the receptor binding domain (RBD) of the SARS-CoV-2 Spike protein (47), generating an accessible RGD motif. *In vitro* data support RBD binding to β1-, β3-, and β6-integrins (48). This led us to assess whether CD98, with its capability to complex with both β1- and β3-integrins, might be a co-receptor for the SARS-CoV-2 Spike RBD, triggering pro-inflammatory responses to the initial viral:host cell encounter.

We therefore assessed interactions of putative gal-3-fibrosome components in IPF progression and acute exacerbations, with a particular focus on the role of CD98. We studied disease tissue from IPF patients, *ex vivo* human, *in vivo* mouse, and *in vitro* cell culture models. We used TGF-β1 treatment to simulate chronic progression and a range of acute stimuli, including the SARS-CoV-2 Spike protein RBD, to simulate exacerbations including COVID pneumonitis.

## Results

### Putative gal-3-fibrosome components in IPF disease tissue and disease models ex vivo and in vitro

We assessed the potential relevance of CD98:β1-integrin in IPF and profibrotic responses to TGF-β1 stimulation in human lung tissue and alveolar epithelial cells. CD98 immunoreactivity was greatly increased in IPF tissue relative to non-fibrotic control at the cellular level in *ex vivo* biopsy tissue, as was galectin-3 and β1-integrin (Fig. 1a). These three proteins co-localised in serial sections (Supp. Fig. 1b). The staining was particularly marked in epithelial cells, confirmed by E-cadherin positive staining. Next, the involvement of CD98 was assessed within a validated *ex vivo* tissue model of early pulmonary fibrosis (50). β1-integrin (ITGB1 gene) is known to demonstrate the second highest of all measured mRNA responses to stimulation with TGF-β1 in this model (50). We found that the CD98 heavy chain (SLC3A2, constant component of the CD98 heterodimer) response was also significantly increased (Fig. 1b, left panel). Interestingly galectin-3 (LGALS3) mRNA was not upregulated by this treatment in whole lung tissue (right panel), indicating that the increased galectin-3 protein signal was not due to an increased transcription response to TGF-β1 across lung tissue as a whole. We further assessed the requirement for CD98 in downstream intracellular TGF-β1 signalling in primary murine alveolar epithelial cells, using a *cre-lox* system to conditionally silence CD98. The data indicate that CD98 is critical for these downstream responses to occur (Fig. 1c, left panel). In the A549 alveolar epithelial cell model, CD98 was required for TGF-β1 to induce an epithelial-mesenchymal transitional (EMT) phenotypic profile with downregulation of E-cadherin and upregulation of α-smooth muscle actin (αSMA) (Fig. 1c, right panel).

**Fig. 1.**
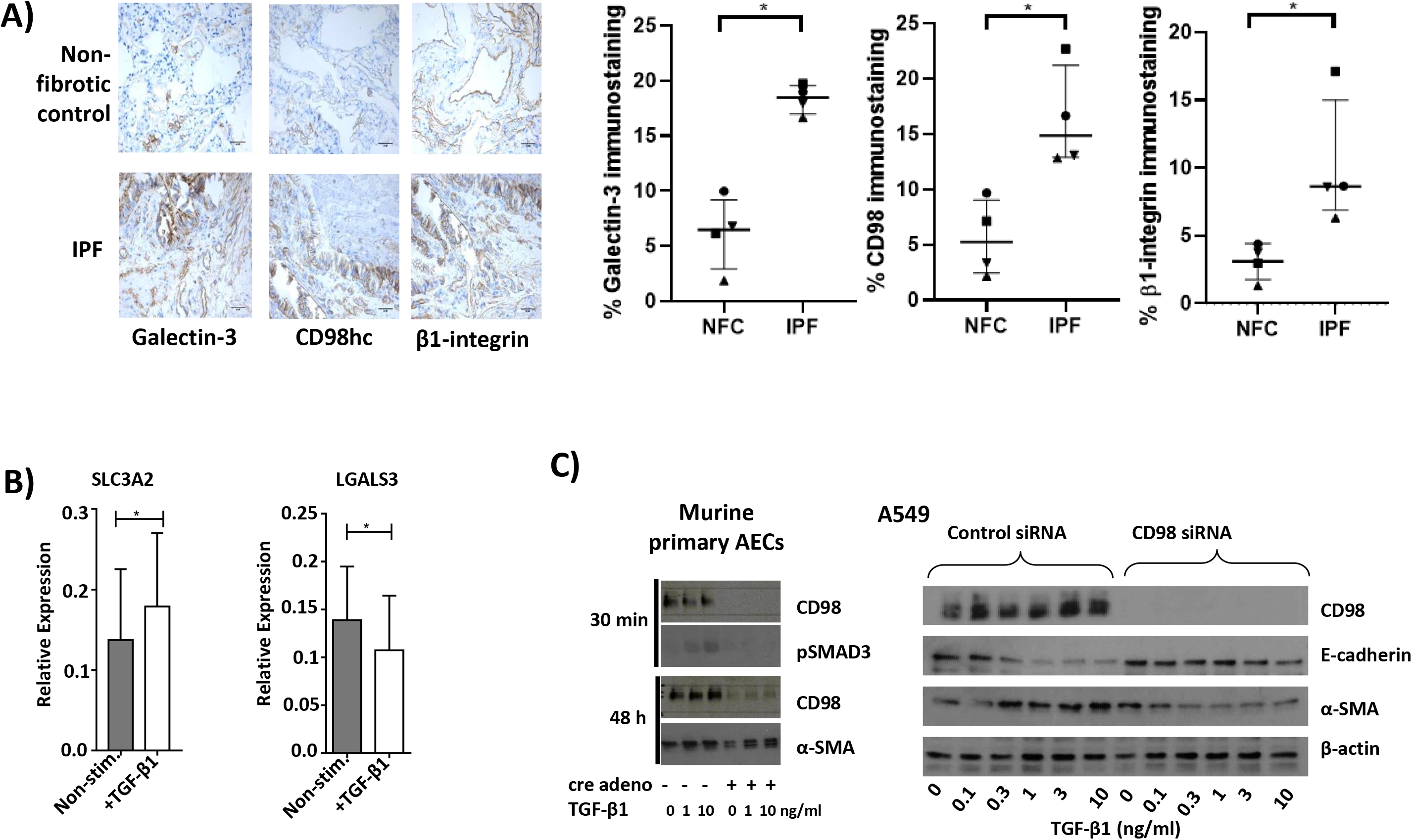
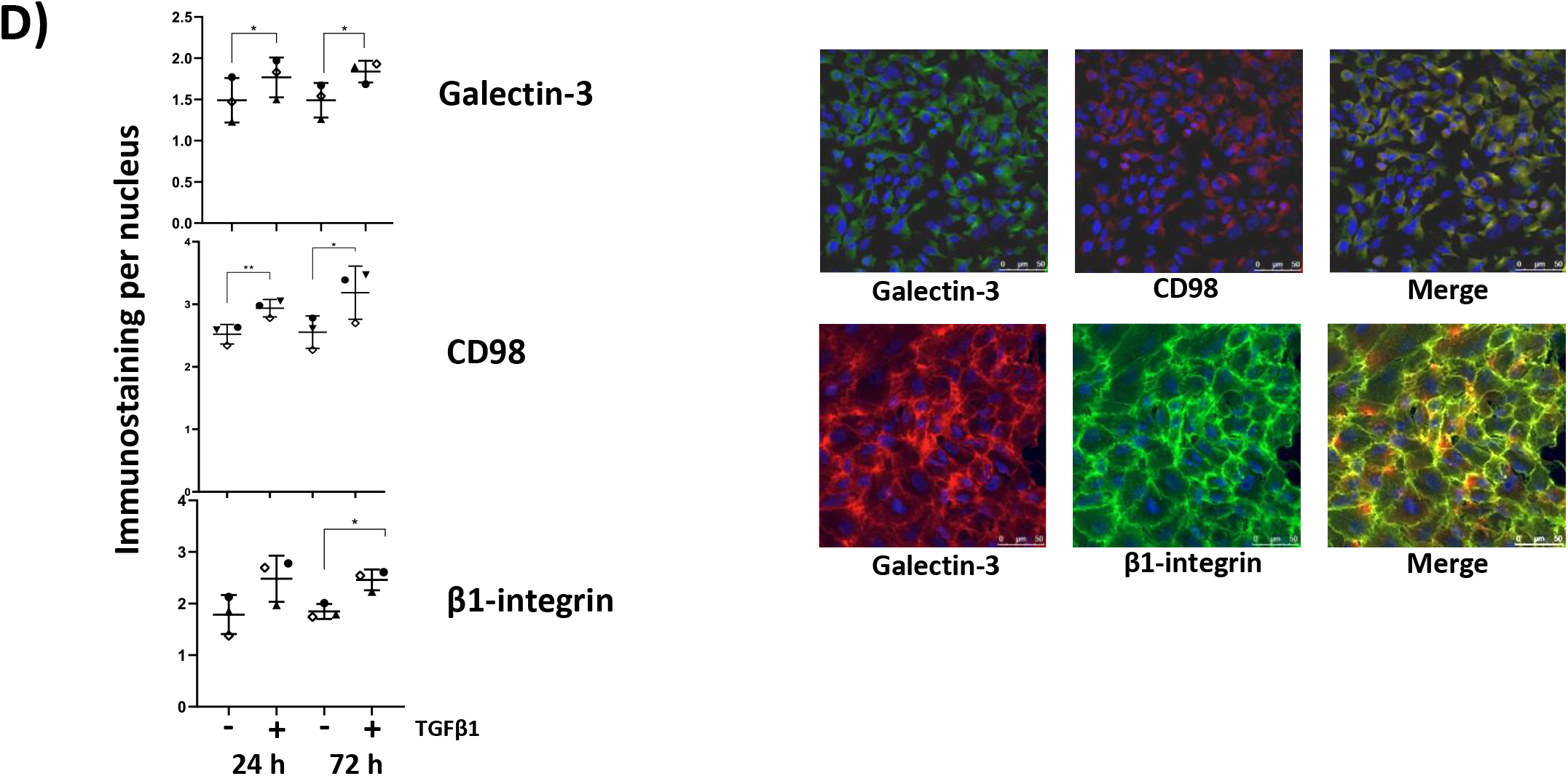
*Ex vivo* and *in vitro* studies of IPF and disease models indicate CD98 co-localises with β1-integrin and galectin-3, increases in response to TGF-β1 in lung epithelial cells, and mediates downstream signalling and function. **a) Galectin-3, CD98hc and β1-integrin protein levels are increased in IPF lung relative to non-fibrotic control lung.** Representative immunostained sections shown (left panels) and signals quantified (right panels. Median values and interquartile ranges indicated. Significance assessed by Wilcoxon signed rank test.) **b) mRNA expression responses to TGF-β1 stimulation in human lung tissue.** Quantitative (q)PCR data. Left panel: SLC3A2 response (increase, p=0.02, n=9) Right panel: LGALS3 response (decrease, p=0.02, n=7) in human lung tissue. **c) *In vitro* EMT marker responses to TGF-β1 are CD98 dependent.** SDS-PAGE probed by western blot. Left panel: Primary alveolar epithelial cells (AECs) from a murine cre-lox conditional CD98 silencing model. Right panel: A549 cells. **d) CD98, β1-integrin and galectin-3 proteins co-localise and increase expression in response to TGF-β1 stimulation in A549 cells.** Confocal immunofluorescence microscopy of galectin-3, CD98, β1-integrin protein responses to TGF-β1 stimulation in A549 cells. Left panel, individual protein levels quantified by single channel immunostaining intensity. Right panel: representative images demonstrating co-localisation. Upper row, galectin-3 labelled green (upper row) or red (lower row) by immunofluorescence, CD98 (upper row) labelled red, β1-integrin (lower row) labelled green. Nuclear staining (DAPI) blue. Merge views shown in yellow.

Treatment with TGF-β1 induced increases in CD98 and its putative binding partners within the gal-3-fibrosome, galectin-3 and β1-integrin, as quantified by western blot (Fig. 1d, left panel) and visualised by immunofluorescence (Fig. 1d, right panel) and these co-localised in a cell surface distribution. CD98 and β1-integrin separated cleanly to the membrane fraction during sample preparation (Supp. Fig. 1c). To confirm whether CD98 and hence its observed interactions localised to the cell surface or to intracellular compartments, we conducted flow cytometry in permeabilising and non-permeabilising conditions (Supp. Fig. 1d). Clear and very similar signals for CD98 were seen in both conditions, indicating all detectable CD98 was indeed distributed on the cell surface.

### Molecular organisation of putative gal-3-fibrosome components basally and in response to TGF-β1 stimulation

We further explored the hypothesis that CD98:β1-integrin complex interacts directly with galectin-3 and hence TGF-βRII within a gal-3-fibrosome using co-immunoprecipitation and proximity ligation assay (PLA). CD98 reciprocally co-immunoprecipitated with galectin-3, with β1-integrin present in both cases (Fig. 2a). Very similar levels of proteins were co-immunoprecipitated with galectin-3 across a titration of 1-10 ng/ml TGF-β1 stimulation (Supp. Fig. 2a). Co-localisation of CD98 and β1-integrin with galectin-3 was detected by PLA in the *ex vivo* human lung tissue model following TGF-β1 stimulation (Fig. 2b). Appearances were consistent with epithelial localisation insofar as cells staining positive were observed to lie between the airspace and basement membrane, but more specific characterisation was not possible with this technique in these samples.

**Fig. 2.**
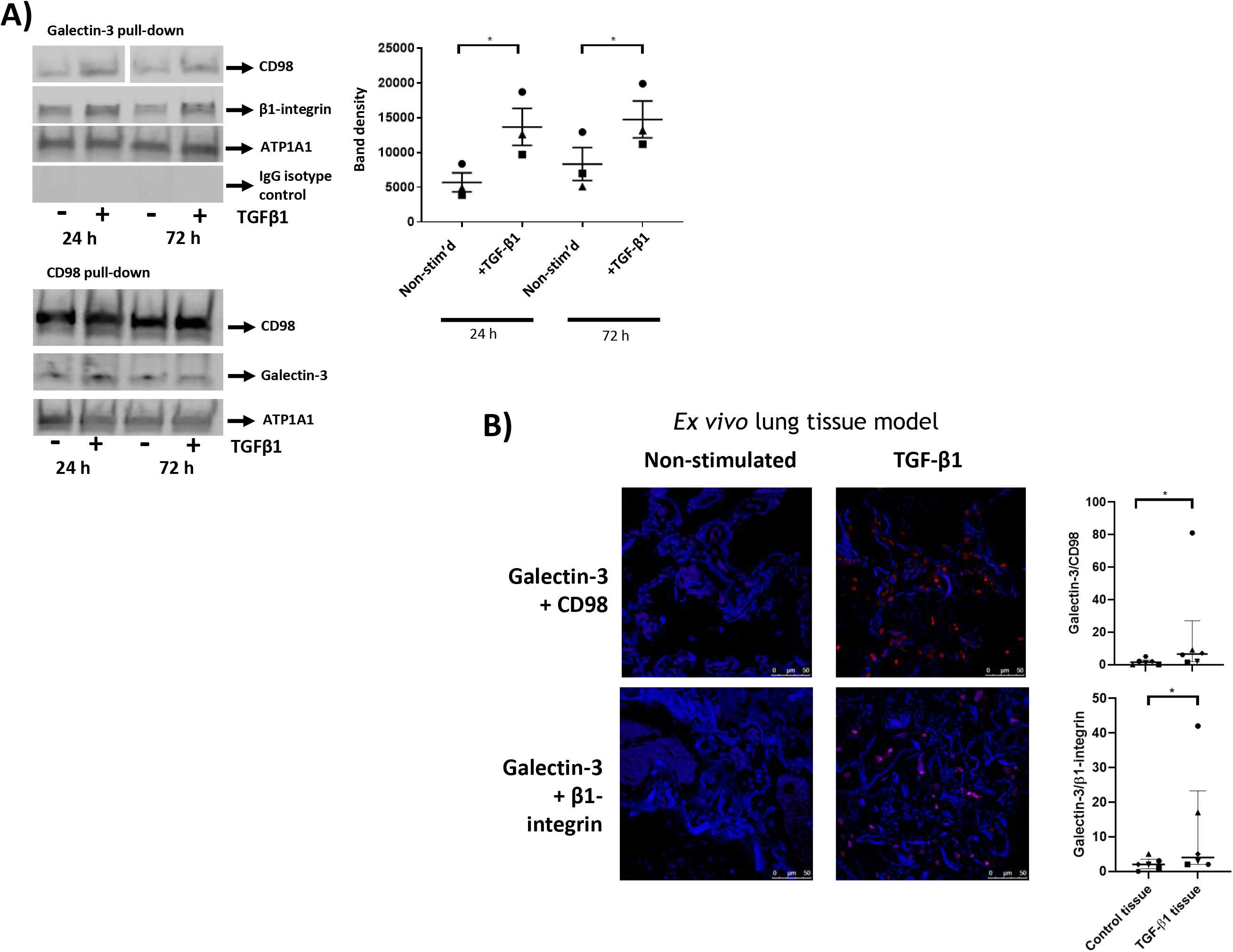
Close co-localisation of multiple putative gal-3-fibrosome components and co-stabilisation of interactions. **a) CD98, β1-integrin and galectin-3 reciprocally co-immunoprecipitate.** SDS-PAGE of A549 lysate membrane fractions probed by western blot. Upper and lower panel sets were taken from separate blots of identical aliquots from the same fractionated cell lysate samples. In the upper panel the gel that was probed with anti-CD98 antibody was run with other lanes between those displayed, these are not shown, but indicated by white space, in order to maintain alignment by condition. ATP1A1 signal serves as a marker for membrane proteins indicating effective fractionation (not seen in cytosolic fractions). IgG isotype control was consistently negative as shown in upper panel. **b) Close co-localisation of galectin-3 and CD98 increases in an *ex vivo* human lung tissue model of progressive pulmonary fibrosis.** Proximity ligation assay (PLA) of galectin-3 and CD98 or β1-integrin in human lung tissue basally and following stimulation by TGF-β1. Nuclear staining (DAPI) in blue, PLA signal detected by red fluorescence following spectral deconvolution to minimise background fluorescence, quantified in right hand plots. Median values and interquartile ranges indicated. Significance assessed by Wilcoxon signed rank test.

### CD98 mediates specific epithelial cytokine response to diverse injury stimuli modelling acute exacerbation events

These findings established that expression of CD98 is increased in pulmonary fibrosis, and that CD98 and β1-integrin closely co-localise with galectin-3 at the cell surface in response to TGF-β1 stimulation in a lung epithelial cell model and in human lung tissue. Moreover CD98 may mediate pro-fibrotic TGF-β1 signalling in lung epithelial cells. We next assessed its capacity to act as a molecular link between responses to acute injury stimuli associated with exacerbations, and chronic progression of pulmonary fibrosis. We began by evaluating its importance in mediating acute responses to experimental stimuli modelling lung injury and triggers of acute exacerbation in ILD, in human lung tissue.

### LPS stimulation in human and mouse lung tissue

Cultured *ex vivo* human lung tissue was stimulated with lipopolysaccharide (LPS, endotoxin), a PAMP that models Gram-negative bacterial infection, and the effects of a pharmacological inhibitor of CD98:integrin dependent function, cynaropicrin (65), were assessed. ELISA measurements of the acute inflammatory cytokines interleukin (IL-)6, IL-8 and tumour necrosis factor (TNF-)α were undertaken. These are associated with ALI (66) and AE-IPF (66, 67).

Cynaropicrin treatment abrogated IL-6 and IL-8 responses, but not TNF-α responses to stimulation with LPS stimulation in human lung tissue (Fig. 3a, Supp. Fig. 3a). An analogous cytokine response signature (including Macrophage Inhibitory Protein (MIP-2), as murine paralog for IL-8) was recapitulated *in vivo* in mice treated with LPS -/+ cynaropicrin (Supp. Fig. 3b). The pathophysiological significance of these effects was supported at the tissue level by the abrogation of the interstitial neutrophil infiltrate response at 24 h (Fig. 3b), for which the IL-8 response would act as a chemotactic cytokine. Over the 24 h timecourse cynaropicrin treatment was not associated with any change in BAL fluid immune cell composition (Supp. Fig. 3 c-i), indicating this was mediated by effects upon lung structural cells rather than alterations in levels of airway lymphocytes or alveolar macrophages. Despite the effects of cynaropicrin upon the parenchymal neutrophilic response, no effects were observed with airway neutrophils, indicating that the former derive from the circulation rather than the airways.

**Fig. 3.**
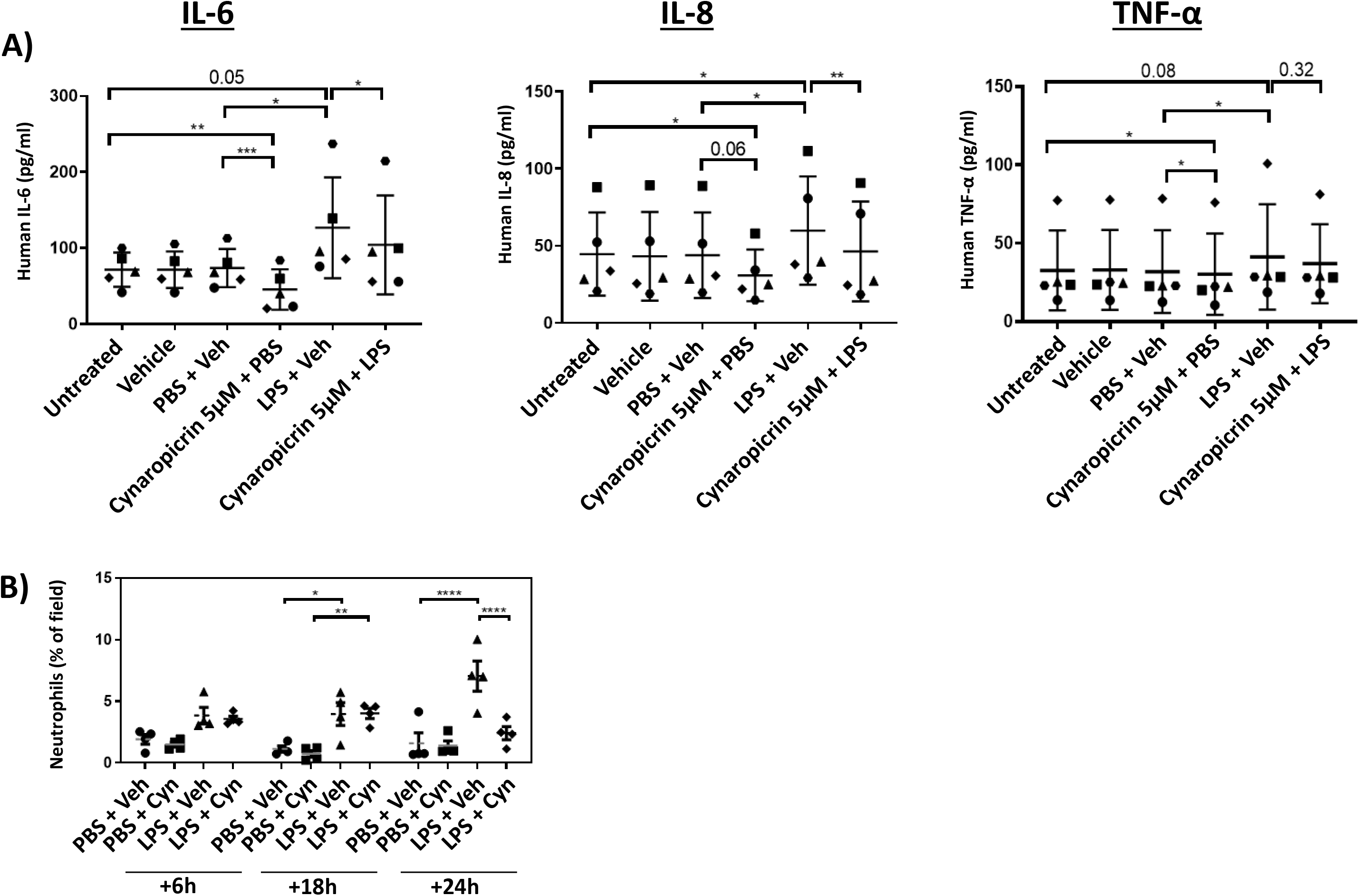
*Ex vivo* human and *in vivo* mouse studies of CD98:integrin-dependence in acute cytokine responses to LPS-induced injury. **a) BAL IL-6 and IL-8 cytokine response signature *ex vivo* (human lung tissue).** IL-6, IL-8 and TNF-α responses to LPS treatment assessed in culture supernatant by ELISA. IL-6 and IL-8, but not TNF-α, responses were abrogated by cynaropicrin treatment. Day 2 post-stimulation shown, donor response comparisons for all timepoints shown in Supp. Fig. 3. **b) Inhibition of CD98:integrin-function is associated with abrogation of tissue inflammatory response downstream of IL-8 paralogue.** Cynaropicrin treatment abrogated interstitial neutrophilia response to LPS treatment *in vivo* in mice at 24 h but not at earlier timepoints. Automated neutrophil count in fields selected, as percentage neutrophil elastase positive pixels within high power field, blinded studies.

### Responses to LPS or mechanical stimuli in alveolar epithelial cell models

We next studied the CD98-dependence and downstream mechanism of the alveolar epithelial cytokine responses to LPS in two different alveolar epithelial cell (TT1 and A549) models. This was demonstrated in both cell models (Fig. 4a, Supp. Fig. 4a) as well as equivalence of cynaropicrin treatment with CD98 silencing (Supp. Fig. 4b). These studies recapitulated the cytokine response profiles observed *ex vivo* in human lungs and *in vivo* in mouse lungs, with IL-6 and IL-8 increases, but not the increase in TNF-α, dependent upon CD98.

**Fig. 4.**
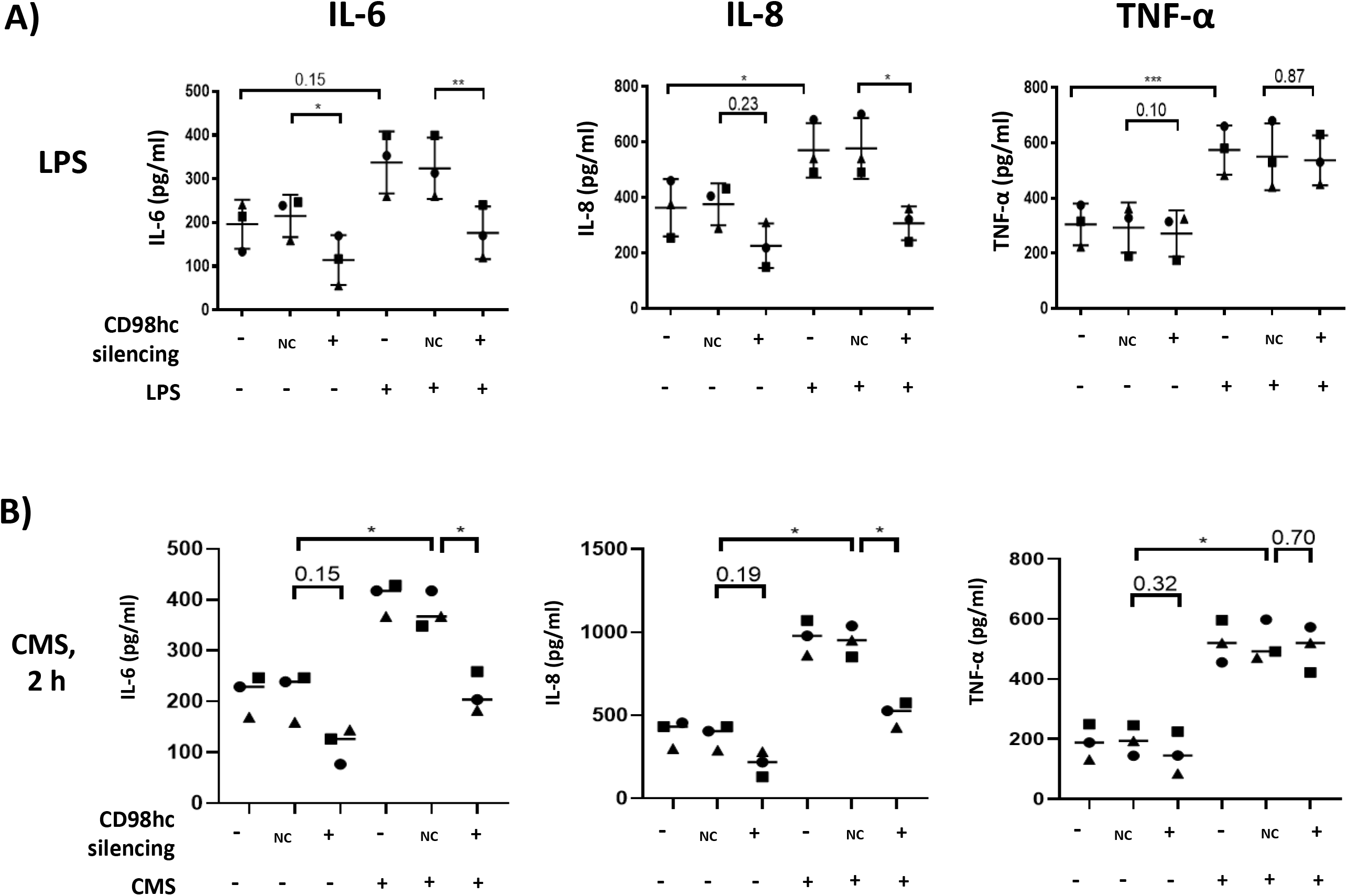
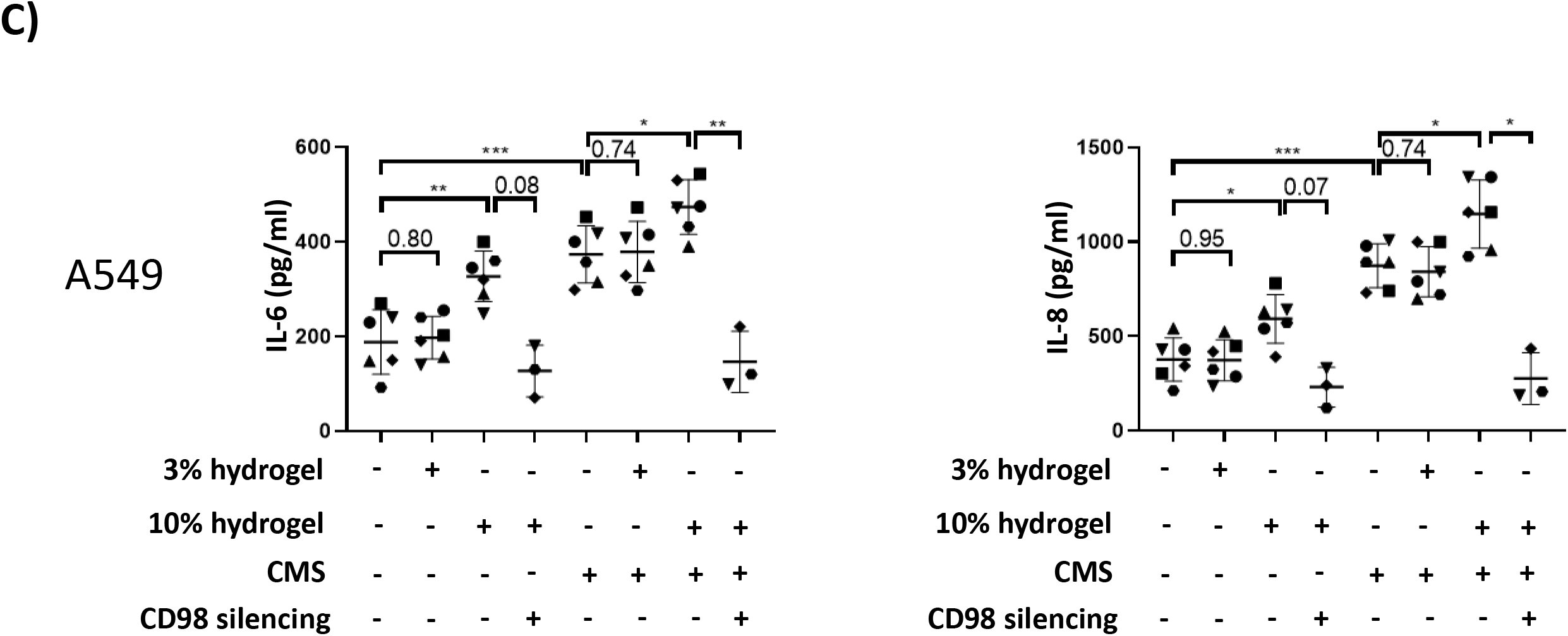
IL-6 and IL-8 responses to diverse stimuli abrogated by CD98 silencing in two different alveolar epithelial cell models. **a) CD98 mediates IL-6 and IL-8 but not TNF-α responses to LPS in alveolar epithelial cells.** ELISA data for IL-6, IL-8 and TNF-α responses to LPS stimulation, and effects of CD98 knockdown in A549 cells. NC indicates non-coding siRNA control condition. (Equivalent data for TT1 cells shown in Supp. Fig. 4a.) **b) CD98 mediates IL-6 and IL-8 but not TNF-α responses to CMS in alveolar epithelial cells.** ELISA data for IL-6, IL-8 and TNF-α responses to CMS stimulation, and effects of CD98 knockdown, in A549 cells at 2 hours. NC indicates non-coding siRNA control condition. (Equivalent data for A549 cells at 24 h and for TT1 cells at both timepoints shown in Supp. Fig. 4c.) **c) CD98 mediates IL-6 and IL-8 responses to matrix stiffness in alveolar epithelial cells.** ELISA data for IL-6 and IL-8 responses to increasing matrix stiffness, and effects of CD98 knockdown, in TT1 and A549 cells.

In parallel we assessed whether these cellular processes were similarly triggered by mechanotransductive stimuli. Mechanical stress in pulmonary fibrosis is imparted by ventilatory stretch in the context of increased matrix stiffness. We have recently demonstrated that IL-6 and IL-8 levels are increased in A549 cells by cyclic mechanical stretch (CMS) conditions that model a conventional mechanical stretch regimen during invasive lung ventilation (70). This is dependent upon amplitude of stretch change rather than peak stretch. We again used CMS was used to model mechanical lung ventilation in isolation (Fig. 4b, Supp. Figs. 4c-d) and in the context of increasing lung matrix stiffness, modelled using polyacrylamide gel matrices (hydrogels) of varying stiffness (Fig. 4c, Supp. Fig. 4e). Cytokine responses were assessed without and with CD98 knockdown.

Our data demonstrated the same CD98-dependence of IL-6 and IL-8 responses in the context of CMS stimulation (Fig. 4b, Supp. Fig. 4c-d) as observed with LPS. Increasing matrix stiffness also increased IL-6 and IL-8 secretion (Fig. 4c, Supp. Fig. 4e). CMS and matrix stiffness had additive effects, that were entirely CD98-dependent (Fig. 4c). We validated the pathophysiological relevance of the stiffness achieved by measuring modulus of elasticity values using indentation. Acrylamide(:bis-acrylamide) hydrogel conditions of 3%(:0.03%) and 10%(:0.3%) (v/v) generated values of 0.445 ± 0.080 kPa and 30.868 ± 3.198 kPa (mean ± SEM) respectively. The latter values fall within the range used to model the pathophysiological state in IPF (68). Addition of LPS to mechanically stimulating conditions resulted in consistent but small further increases in IL-6 or IL-8 secretion that did not consistently reach p<0.05 significance thresholds at n=3 (Supp. Fig. 4f). For these experiments, p-values were lower in TT1 than A549 cells, and for IL-8 responses than for IL-6 responses.

### CD98, TRPV4 and NF-κB co-operate in positive feedback to generate IL-6 and IL-8 responses to acute injury stimuli

Intracellular signalling pathways were screened to identify which mediated CD98-dependent responses (Fig. 5 and Supp. Fig. 5). These experiments identified the NF-κB signalling pathway in both responses to LPS (Fig. 5a, Supp. Fig. 5a) and CMS (Fig. 5b). They did not support involvement of phospho-p44/42 ERK, p38 (Supp. Fig. 5b-c) or JNK (Supp. Fig. 5d) signalling pathways. The critical importance of NF-κB signalling was supported by pharmacological inhibition studies with the small molecule JSH-23, which generally reduced IL-6 and IL-8 alveolar epithelial cell responses to LPS, (Fig. 5c), CMS (Fig. 5d) and matrix stiffness (Fig. 5e). At n=3, the JSH-23 conditions used were less clearly effective at reducing IL-6 and IL-8 responses to LPS than the mechanotransductive stimuli. However increasing the experimental repeats to n=8 in A549 cells treated with LPS demonstrated comparable magnitudes of abrogation and significance to those achieved for both cell models with JSH-23 treatment counteracting the effects of CMS or matrix stiffness. These findings suggest that the observed differences likely arise from variability of LPS response rather than involvement of distinct mechanistic pathways downstream of CD98.

**Fig. 5.**
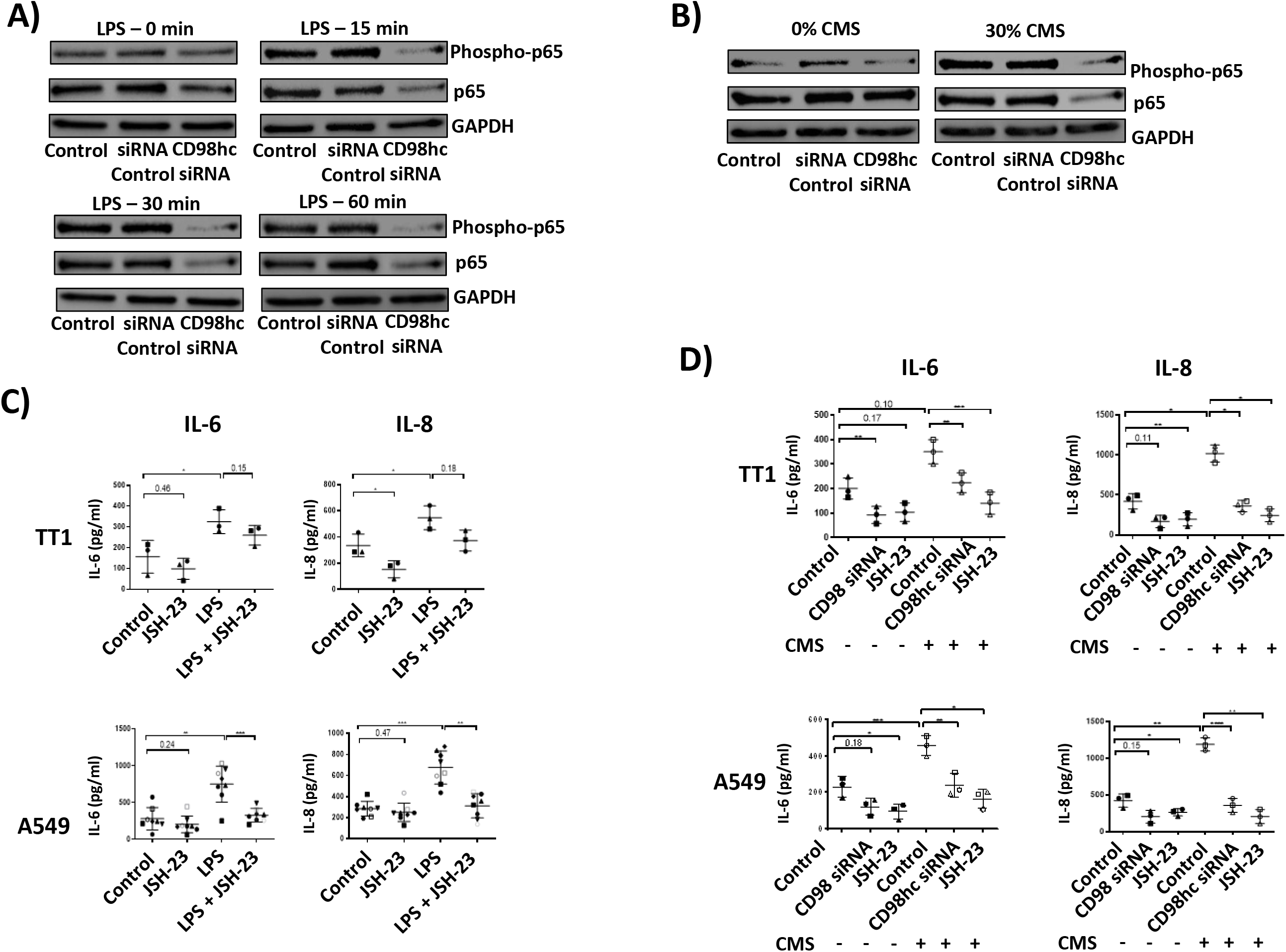
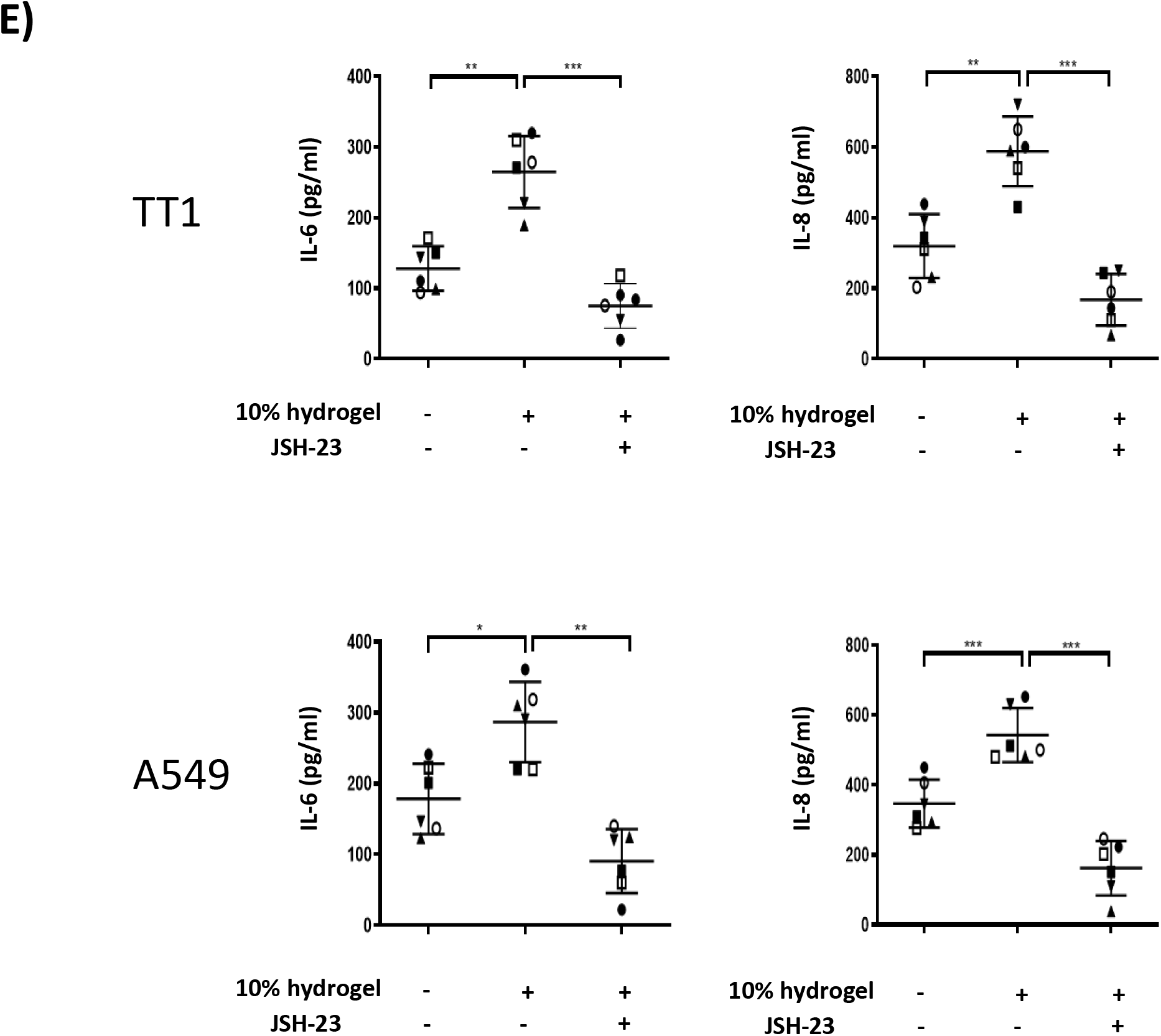
Role of NF-kB in CD98-dependent cytokine responses. **a)-b) NF-κB activation responses to injury stimuli are CD98-dependent.** CD98hc-dependence was observed for NF-κB pathway activation in the response of A549 cells to LPS (a) and CMS (b) stimulation, as assessed by SDS-PAGE probed with western blot. **c)-e) CD98-mediated cytokine responses to injury stimuli are mediated by NF-κB.** IL-6 and IL-8 responses to LPS (c), CMS (d), and matrix stiffness (e) assessed by ELISA were abrogated by NF-κB inhibition with the small molecule JSH-23 (50 μM) in TT1 and A549 cells.

We next hypothesised that CD98-dependent responses to acute injury stimuli might involve Ca^2+^ influx through the TRPV4 Ca^2+^ channel as this protein associates with β1-integrin and is implicated in mechanotransduction responses in myofibroblasts (68, 69) and a *Xenopus* oocyte expression model (70). Chelation of extracellular Ca^2+^ by EGTA or inhibition of TRPV4 reduced CD98-dependent cytokine injury responses to LPS and CMS whilst stimulation with a TRPV4 agonist potentiated the effects (Fig. 6a). These effects were consistent between TT1 and A549 cell lines and IL-6 and IL-8 cytokines. Conversely, CD98 silencing had similar effects to EGTA treatment or TRPV4 inhibition upon Ca^2+^ influx measured immediately following hypo-osmolar stretch, supporting the role of TRPV4 in mediating CD98-dependent injury responses in alveolar epithelial cells (Fig. 6b). Taken together with the CD98-dependence of the same responses to non-osmolar stretch, these findings also indicate the well-known ‘osmo-receptor’ function of TRPV4 may be mediated by CD98hc response to cellular stretch.

**Fig. 6.**
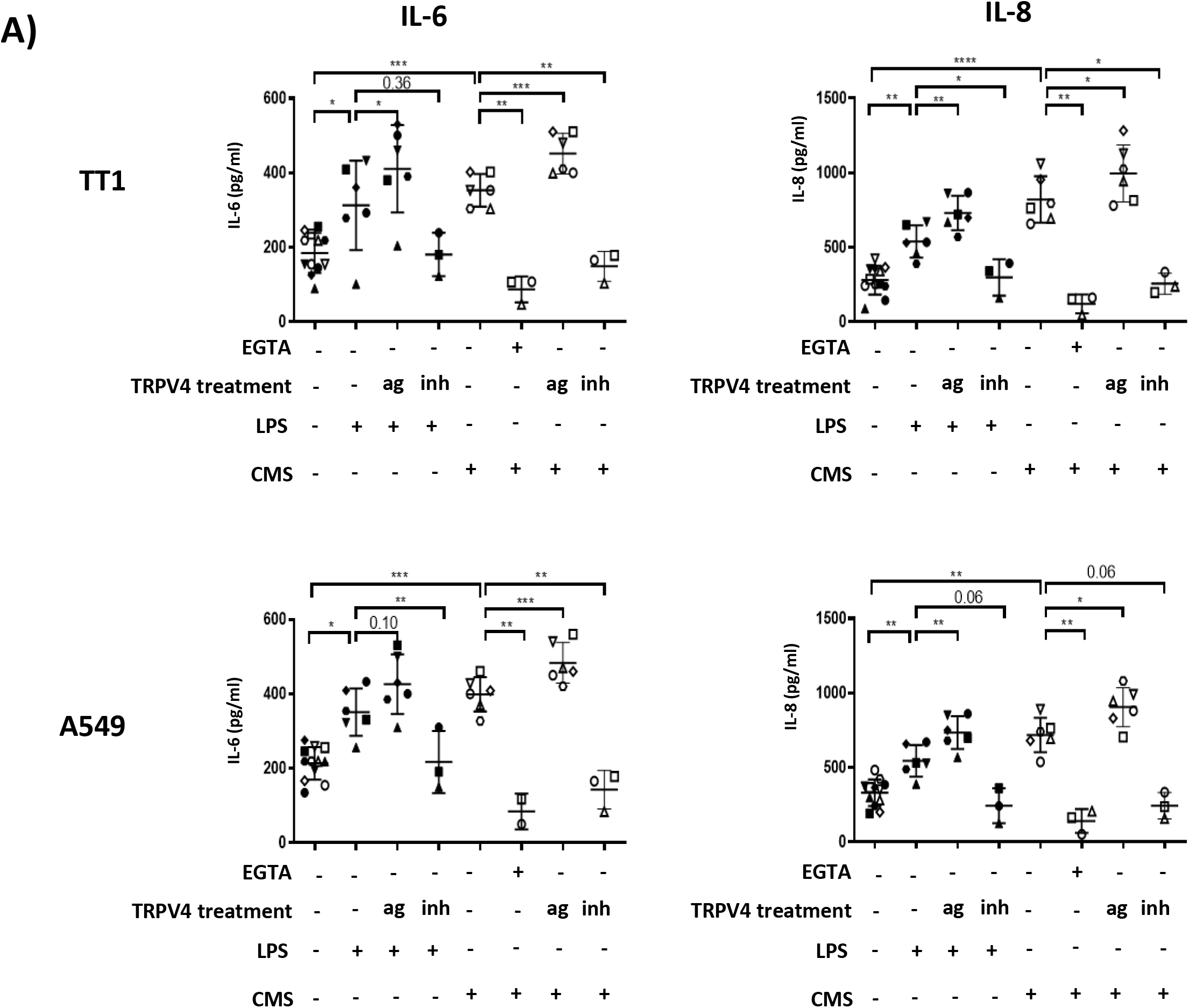
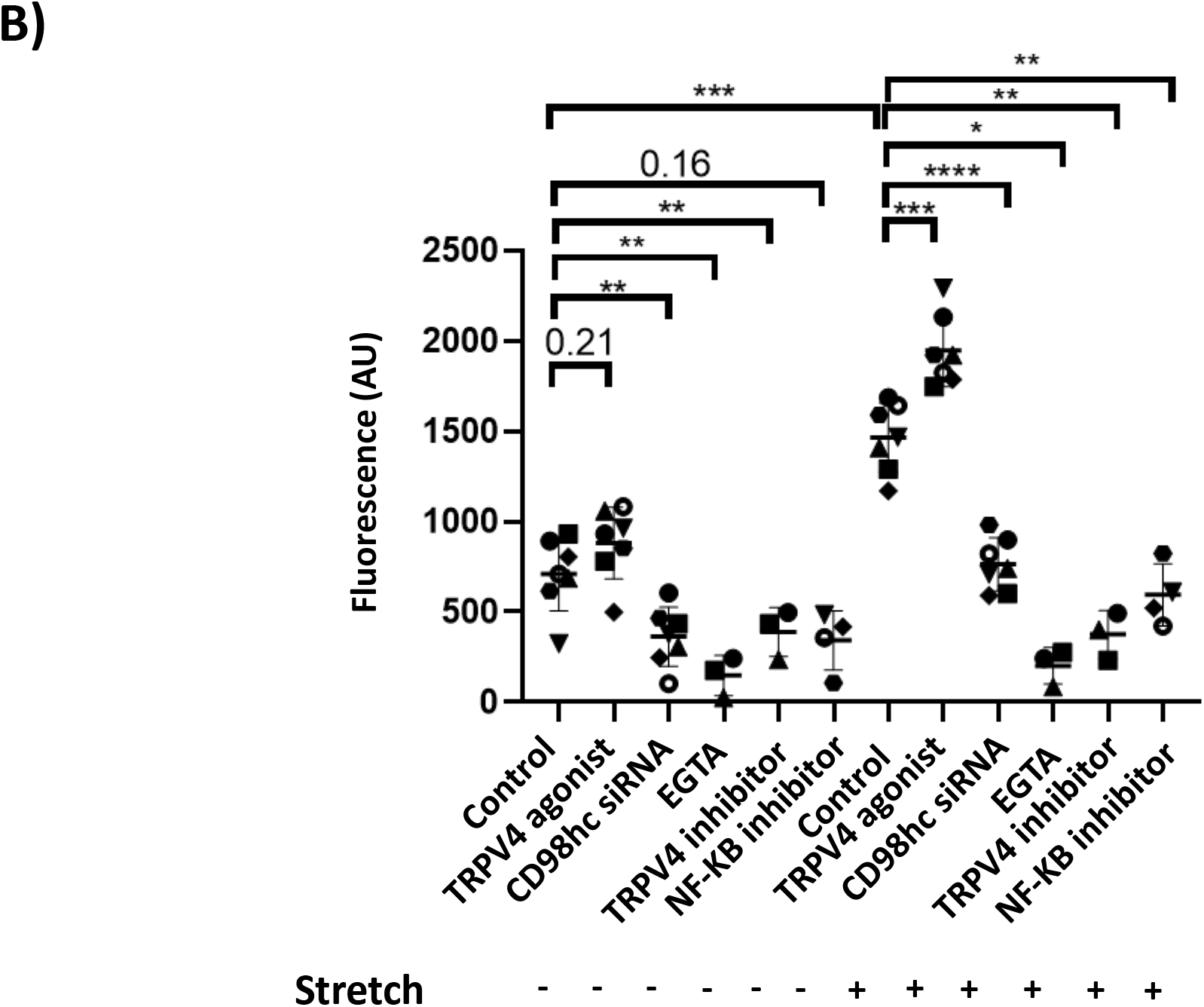
TRPV4-mediated Ca^2+^ influx and NF-κB signalling mediate the CD98-dependent IL-6 and IL-8 response in a positive feedback loop. **a) IL-6 and IL-8 responses to stretch vary with Ca^2+^ influx into cells and TRPV4 activation.** TRPV4 agonist (ag) or inhibitor (inh) treatments tended respectively to enhance or attenuate IL-6 and IL-8 responses to LPS and CMS in TT1 and A549 cells, measured by ELISA. Chelation of extracellular Ca^2+^ abrogated such responses. **b) CD98 and NF-κB mediate cellular Ca^2+^ influx responses to stretch.** Intracellular Ca^2+^ quantified by Fluo-4 assay in A549 cells. Knockdown of CD98hc affected Ca^2+^ influx similarly to EGTA treatment or TRPV4 inhibition. Remarkably, similar reductions were also observed upon inhibition of intracellular, and hence presumably ‘downstream’, NF-κB activity using JSH-23, supporting the hypothesis that CD98, TRPV4 and NF-κB cooperate in a pro-inflammatory positive feedback loop. Stimulation of NF-κB activity in Ca^2+^ chelating conditions (Supp. Fig. 6b) increased IL-6 and IL-8 secretion supporting the effector role of NF-κB.

Unexpectedly, treatment with pharmacological inhibitors of either TRPV4 or NF-κB caused a clear reduction in cellular CD98hc levels assessed by western blot, particularly in response to injury stimuli. These findings suggested a positive feedback loop whereby activity of these proteins stabilise cell surface levels of CD98 (Supp. Fig. 6a). To test this hypothesis, we measured the Ca^2+^ influx through TRPV4 in response to stretch and assessed the degree to which inhibition of intracellular NF-κB could reduce this. NF-κB inhibition entirely abrogated stretch-induced Ca^2+^ influx in a similar maner to CD98hc silencing, EGTA treatment or TRPV4 inhibition (Fig. 6b). These findings strongly support a positive feedback process in which CD98-mediated injury responses activate TRPV4 to increase intracellular Ca^2+^ and NF-κB activation, which, in turn stabilises CD98 levels. NF-κB agonism with phorbol myristate acetate (PMA) restored IL-6 and IL-8 responses when Ca^2+^ influx was prevented by chelation of extracellular Ca^2+^ using EGTA (Supp. Fig. 6b). These data support the role of NF-κB as the specific effector of cytokine release in this system.

### SARS-CoV-2 Spike Receptor Binding Domain interacts with CD98 to stimulate IL-6 and IL-8 release in lung epithelia and macrophages, independent of cellular infectivity

In light of these findings, the onset of the COVID-19 pandemic caused us to explore the possibility that the CD98:integrin complex might be involved in mediating the severe inflammatory response in lung tissue and from macrophages. The causative virus SARS-CoV-2 initially interacts with host cells via its spike protein. The receptor for the closely related spike protein of SARS-CoV-1 primarily interacts with a cell surface form of angiotensin converting enzyme (ACE)2 (71). This is abundantly expressed in nasal epithelia (72) and so its potential role in facilitating transmission of infection is clear. However, although COVID pneumonitis is a common consequence of SARS-CoV-2 and is the predominant cause of acute mortality in COVID-19, ACE2 is expressed at low levels in lung parenchyma (detectable in ∼1% of Type 2 alveolar epithelial cells) (73). These observations raised the prospect of alternative or co-receptors mediating the lung parenchymal inflammatory responses. We hypothesised that CD98:integrin complex was a candidate for this. The SARS-CoV-2 Spike protein has evolved an RGD tripeptide integrin-recognition motif absent in SARS-CoV-1, within its receptor binding domain (RBD). This can bind to β-integrins that are binding partners of CD98, including β1- and β3-integrin (74). IL-6 responses appear to be a key feature of the pathogenesis of COVID pneumonitis. This response is characterised by high levels of IL-6, and specific targeting of IL-6 signalling with IL-6 receptor blockade ameliorates disease severity and reduces mortality (75). High IL-6 and high IL-8 levels are correlated with illness duration and severity (76–78).

We explored this hypothesis using A549 cells that do not detectably express ACE2 at protein level basally (79), and subsequently in Calu-3 cells that do (80). Macrophages are also important mediators of acute inflammatory responses, implicated in the COVID pneumonitis response (81), and CD98:galectin-3 interactions are known to play a role in macrophage responses (36). We therefore also studied THP1 cells, induced into a macrophage (M0) phenotype by PMA treatment. We titrated recombinant SARS-CoV-2 Spike RBD made in HEK293 cells against these cell types. The lowest RBD concentrations generating peak cytokine responses in the titrations were chosen (for A549 cells: 2 μg/ml, for THP1 cells: 10 μg/ml, for Calu-3 cells: 35 μg/ml). Treatment with bovine serum albumin as a protein control for non-specific binding did not yield any cytokine responses at the same concentrations.

In all three cell lines SARS-CoV-2 Spike RBD stimulation induced equivalent IL-6 and IL-8 responses, but minimal TNF-α responses, consistent with mediation by a CD98-dependent mechanism (Fig. 7a). For A549 and THP1 cells the IL-6 and IL-8 responses were of similar magnitude to those detected using the LPS treatment conditions applied in A549 and TT1 cells previously. The effects of cynaropicrin treatment and of CD98 silencing on these responses were compared with the effects of dexamethasone (Supp. Fig. 7a). Both dexamethasone and cynaropicrin treatment entirely abrogated IL-6 and IL-8 responses to the SARS-CoV-2 Spike RBD in both A549 and THP1 cells (Fig. 7a). For the IL-6 response in A549 cells, effects remained statistically significant at the same thresholds if the data from the repeat generating high magnitude outlier points were removed. Silencing of CD98 or galectin-3 (Fig. 7b) were similarly effective in abrogating these responses in the A549 and Calu-3 lung epithelial cell lines. As previously described by other groups, gene silencing approaches were inefficient in the THP1 cell line in our hands, with no detectable change in protein levels observable by western blot. We demonstrated co-immunoprecipitation of CD98 with the SARS-CoV-2 Spike RBD following stimulation, using two different anti-RBD antibodies in A549 cells (Fig. 7c). Taken together these findings indicate that SARS-CoV-2 Spike RBD interacts closely with CD98 to trigger IL-6 and IL-8 release in lung epithelial cells and macrophages, via a mechanism that appears highly conserved between diverse stimuli as well as cells (Fig. 7d).

**Fig. 7.**
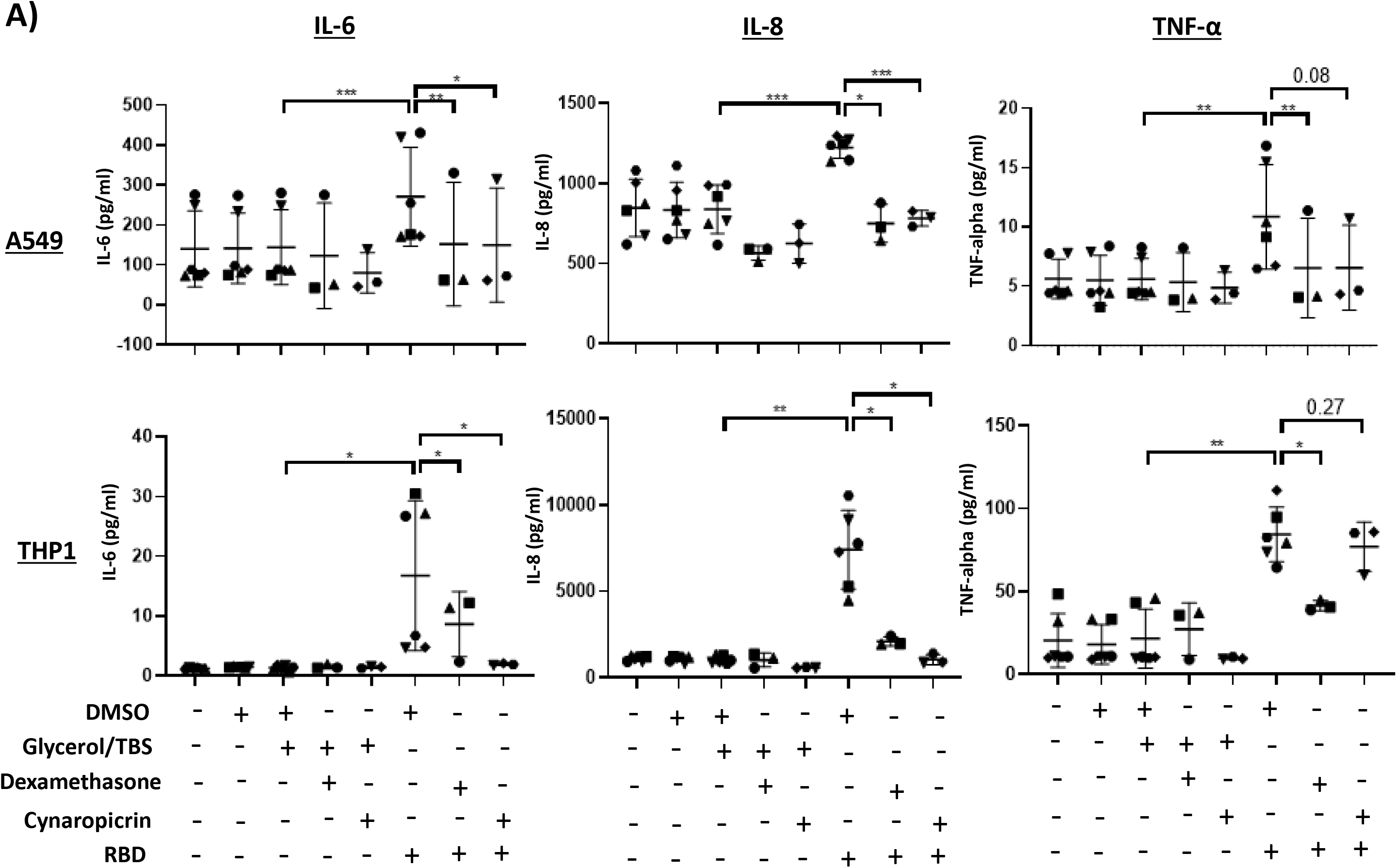
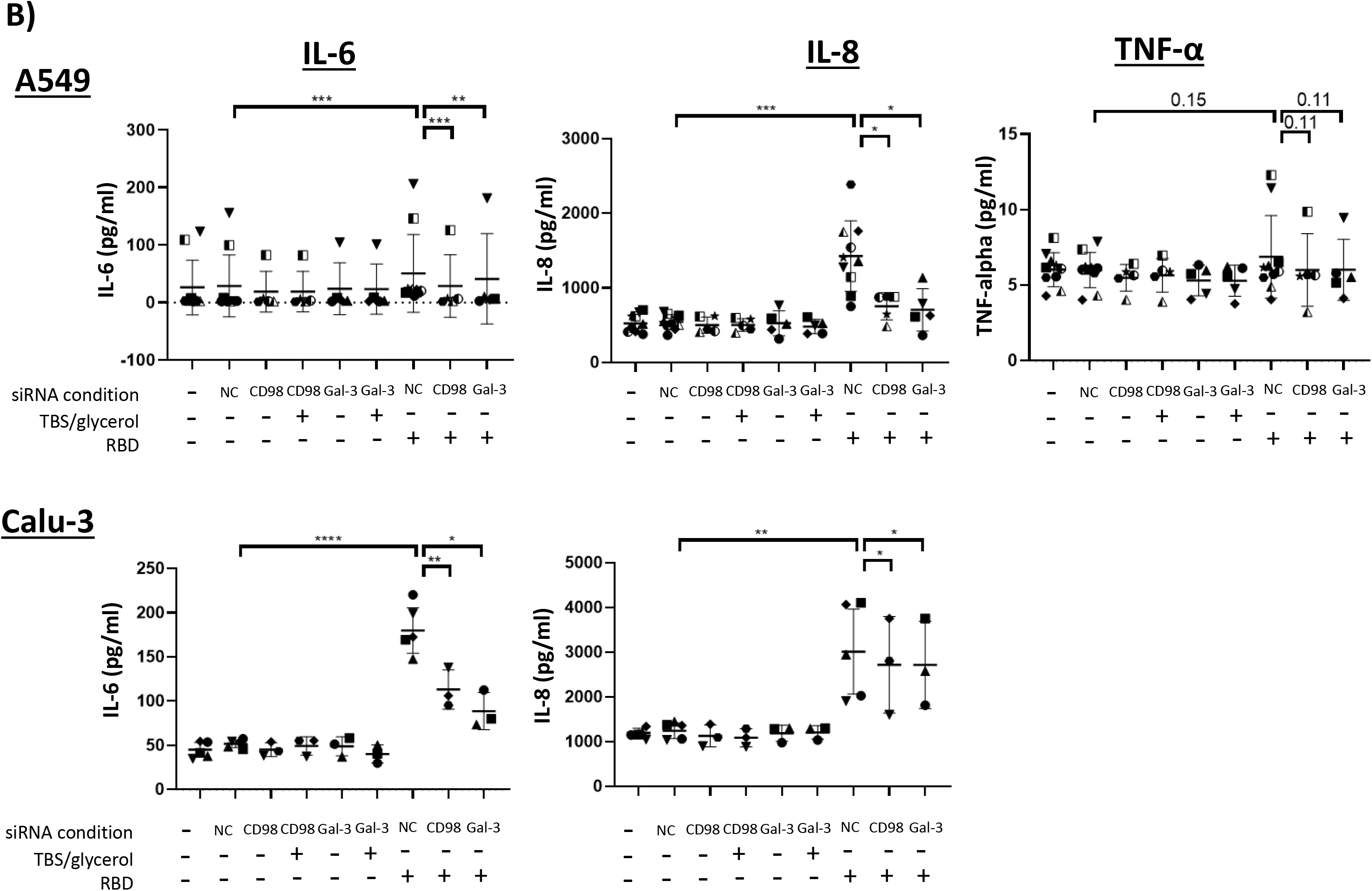
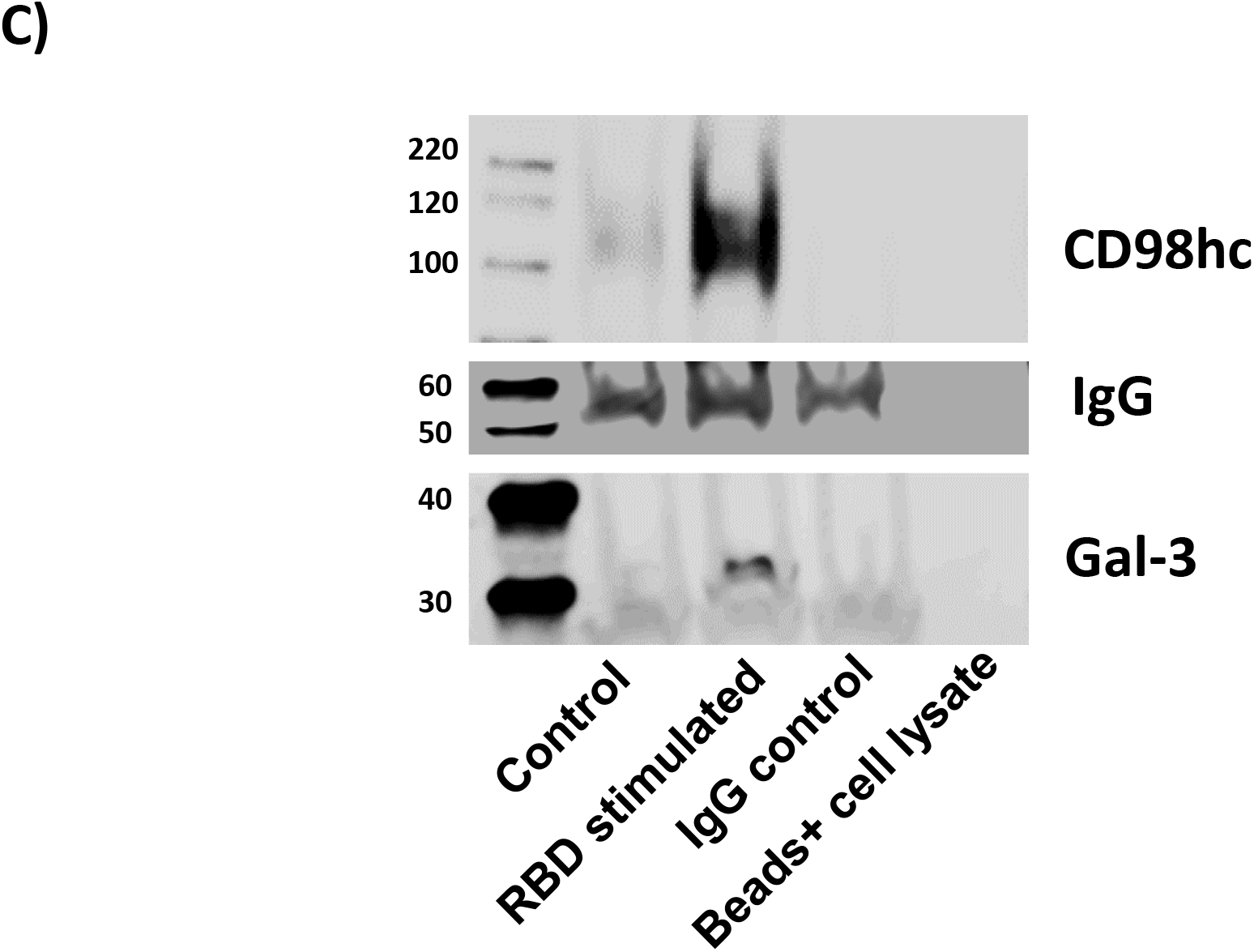
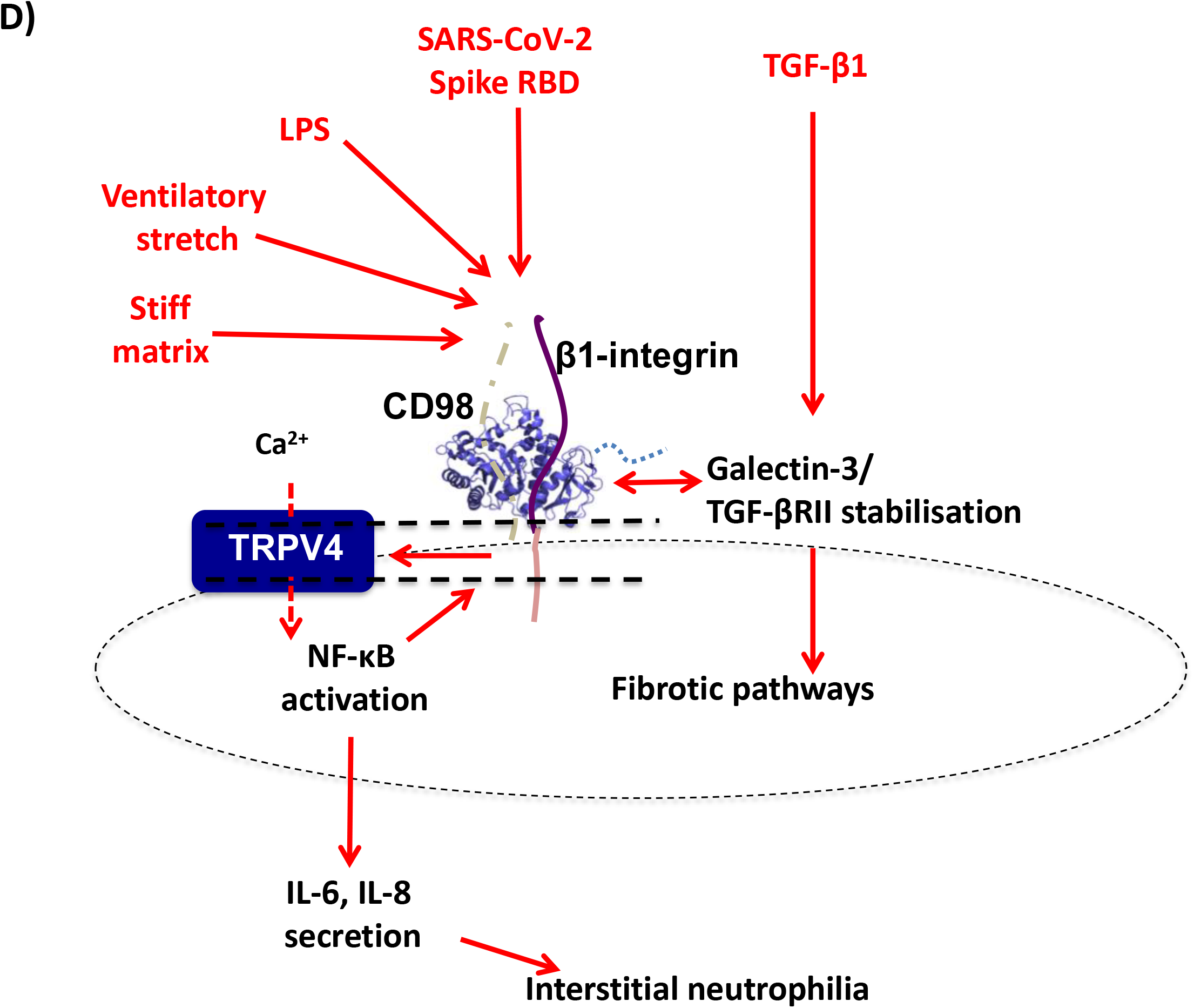
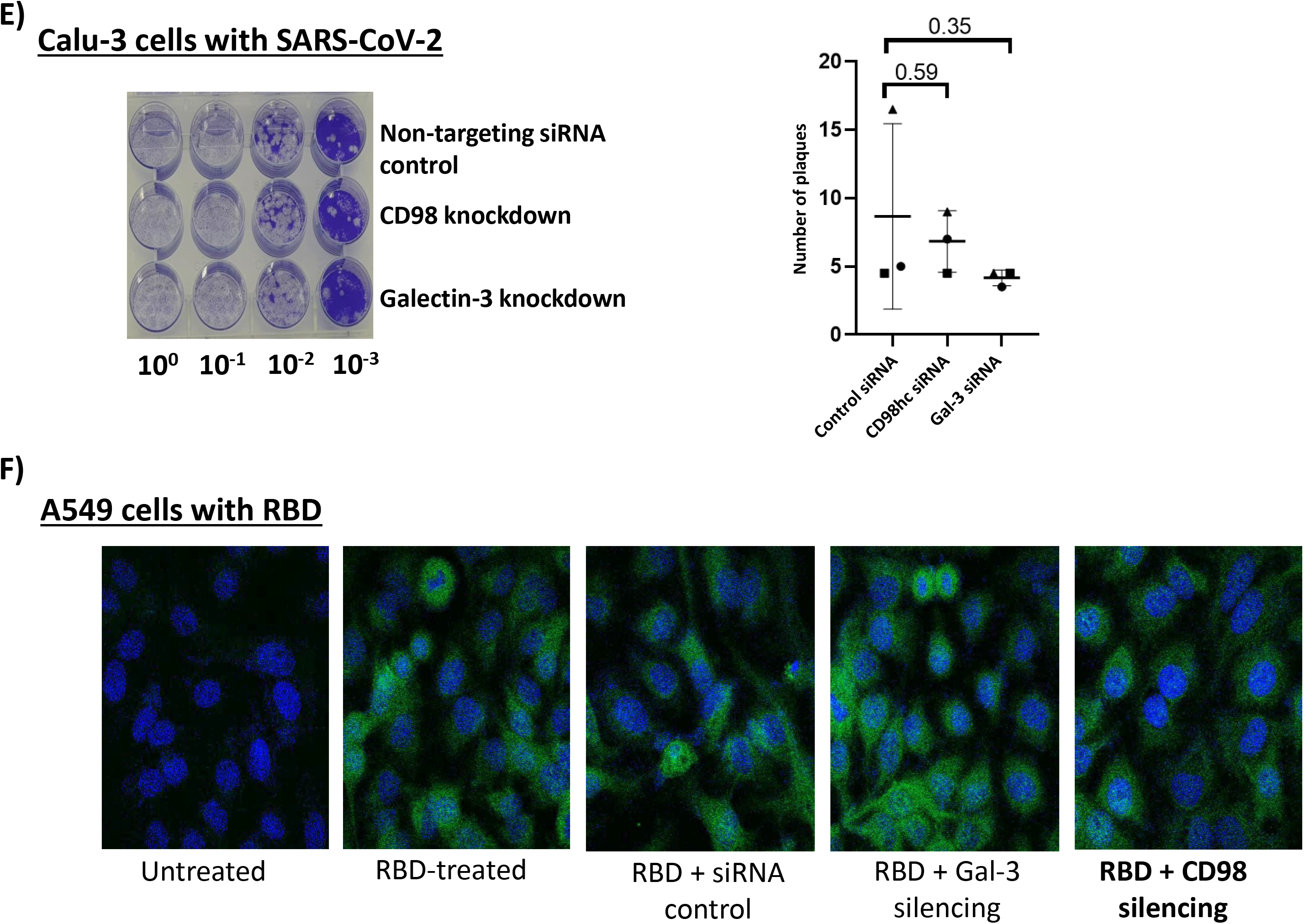
SARS-CoV-2 spike protein RBD utilises a CD98-dependent pathway to drive IL-6 and IL-8 responses in lung epithelial and monocyte-derived macrophage cell models. **a) RBD stimulates a IL-6 and IL-8 response signature to SARS-CoV-2 spike protein that is modifiable by therapeutic agents, in diverse cell lines.** ELISA data reporting IL-6, IL-8, and TNF-α release from a lung epithelial (A549) and a macrophage (THP1) cell line in response to the RBD of the SARS-CoV-2 spike protein. The experiments assessed the effects of an established treatment for COVID (dexamethasone) with cynaropicrin. Similar response profiles (strong IL-6 and IL-8 release signals, minimal TNF-α response) were seen in both cell types. IL-6 and IL-8 responses were similarly abrogated by treatment with either dexamethasone or cynaropicrin. **b) CD98 and galectin-3 mediation of IL-6 and IL-8 responses to RBD in lung epithelial cells.** The role of CD98 and galectin-3 in these responses was confirmed by knockdown in two lung epithelial cell lines. A549 cells (upper panel) do not detectably express ACE2 protein. Calu-3 bronchial epithelial cells (lower panel) express ACE2 protein and can be infected by SARS-CoV-2. siRNA silencing conditions denoted – (no siRNA construct), NC (non-coding control) or for knockdown of CD98 or gal-3 (galectin-3). Exclusion of data from repeat conditions yielding high outlier IL-6 values in A549 cells did not change the statistical significance of findings relative to conventional p-value thresholds. Calu-3 cells did not mount any detectable TNF-α response to SARS-CoV-2 Spike RBD stimulation in the conditions where IL-6 and IL-8 responses were observed in the preceding Spike RBD titration experiment. Knockdown is not effective in the THP1 macrophage model so similar CD98 silencing was not possible in that cell line. **c) SARS-CoV-2 spike protein RBD engages with host cell CD98.** Western blots of SDS-PAGE assessing co-immunoprecipitation from lysates of A549 cells stimulated with the RBD. Similar findings following pull-down with the polyclonal anti-Spike Protein S1 antibody (Invitrogen, catalogue number PA5-81795) and probing with same antibody or rabbit anti-RBD mAb (Sino Biological, catalogue number 40150-R007, shown here). CD98, and to a lesser extent galectin-3, co-immunoprecipitated with the Spike RBD. IgG levels shown as loading control. **d) Schematic of CD98-dependent mechanism of inflammatory IL-6 and IL-8 responses to diverse injury stimuli.** The data in this paper support close molecular proximity of CD98 to galectin-3 at the cell surface and its associated pro-fibrotic pathways. **e) SARS-CoV-2 infectivity in Calu-3 cells does not require CD98.** Representative plaque forming assay shown (left panel) with quantification over three repeats plotted (right panel). No infection could be demonstrated in A549 cells that lack ACE2 by this assay. **f) RBD binding to A549 cells does not require ACE2, galectin-3 or CD98.** Despite the absence of detectable ACE2 in A549 cells RBD was able to bind to the surface of A549 cells even when CD98 or galectin-3 were knocked down.

Despite this evidence of an initial pro-inflammatory interaction between the SARS-CoV-2 Spike RBD and A549 cells, as expected, the A549 cells could not be infected by SARS-CoV-2. This is consistent with the lack of intrinsic ACE2 expression, and so indicates that such Spike RBD:CD98 interactions are not sufficient for infection in our experimental conditions. Conversely in Calu-3 cells, that serve as a lung epithelial line that expresses ACE2 and can be infected by SARS-CoV-2, silencing of CD98 did not reduce infectivity (Fig. 7e). Silencing of CD98 or galectin-3 did not clearly affect SARS-CoV-2 Spike RBD binding to A549 cells (Fig. 7f). These findings suggest this protein can interact with lung epithelial cells in a number of ways, and that the pro-inflammatory interaction mediated by CD98 is likely distinct from the pro-infective interaction mediated by ACE2.

## Discussion

Previous work demonstrates that galectin-3 is increased in IPF tissue and mediates fibrotic responses to TGF-β1 or bleomycin. Functional and direct binding interactions of galectin-3 with TGF-β1 signalling and the TGF-β receptor (TGF-βR)II protein respectively indicate how this may cluster transmembrane proteins to potentiate fibrotic responses. The molecular structure of galectin-3 provides potential to bind a wide range of proteins and glycoproteins beyond this (28, 37, 82–84). This plausibly provides a molecular crosslinking platform that may recruit, augment or modify pathogenic responses in fibrosing ILDs in communication with both extracellular and intracellular triggers. We use the term ‘gal-3-fibrosome’ to describe this general concept of a pro-fibrotic macromolecular assembly nucleated at the cell surface by galectin-3. The translational potential of this approach is attested by the development of small molecule inhibitors of galectin-3:glycan interactions currently in late phase clinical studies as a treatment for IPF.

To explore the gal-3-fibrosome concept we have characterised the behaviour of CD98, which interacts with galectin-3 and has been proposed as a direct binding partner. Using *ex vivo* human lung tissue, *in vivo* mouse, and a number of cell models we specifically focused upon the potential for CD98 to mediate acute injury responses and acute exacerbations of fibrosing ILDs and associated molecular mechanisms. We have also followed up on observations from COVID pneumonitis suggesting that this pathway might play a role in associated epithelial and macrophage hyper-inflammatory responses to initial SARS-CoV-2 virus:host cell interactions.

Interruption of pro-inflammatory CD98-mediated signalling could be targeted by direct CD98 inhibition or disrupting interactions with TRPV4 to develop a novel therapeutic for exacerbations of ILD. Use of a TRPV4 inhibitor developed commercially was deemed unsuccessful in a human subject model of ALI (85). However this was not fully effective as a TRPV4 inhibitor, and further optimisation of this approach could therefore be considered. Moreover it was not tested in the context of individuals whose lung tissue was primed for activation of this pathway as our data suggest is the case in patients with IPF.

The findings in this paper establish that IPF lung tissue demonstrates increased protein levels of CD98 and a constitutive integrin binding partner compared with non-fibrotic control lung tissue. These proteins co-localise at the cellular level with galectin-3 in IPF epithelia. At the subcellular level they are found in close molecular proximity (<40 nm) in an *ex vivo* model of early TGF-β1-induced lung fibrosis.

We have identified a role for CD98 in mediating specific innate inflammatory responses to multiple diverse injury stimuli in 2 cell models, an *in vivo* mouse model and *ex vivo* human lung tissue (Fig. 7e). CD98 appears to cooperate in positive feedback with TRPV4 activation, Ca^2+^ influx and NF-κB signalling to amplify IL-6 and IL-8 secretion. Data from the mouse model support the significance of such responses at the organ level, and the therapeutic potential of inhibiting this pathway. Our additional *in vitro* studies support the potential for this pathway to mediate observed cytokine responses from lung epithelial cells and macrophages upon initial interaction with the SARS-CoV-2 Spike protein receptor binding domain. Our data indicate this interaction is distinct from interactions involving the ACE2 protein that mediate viral infectivity.

Taken together the findings provide a molecular mechanism that may explain the increased severity of COVID pneumonitis in individuals with IPF and other fibrotic ILD (86), and their increased risk of acute exacerbations from diverse aetiologies. Given the involvement of CD98 in pro-fibrotic responses to TGF-β1 stimulation in cells and mice, this mechanism could also provide a pathway for cross-talk transducing pro-inflammatory into pro-fibrotic responses.

However acute pneumonitis only results in fibrotic ILD in a small proportion of cases. When it does arise, the fibrosis is usually stable rather than progressive. These observations indicate that pro-inflammatory mechanisms occurring over a single acute presentation do not necessarily generate pro-fibrotic behaviour in the lung parenchyma. It is therefore likely that further molecular events are required for this to occur (Supp. Fig. 7). These might arise in association with chronic, low-grade activation of innate inflammatory pathways such as the epithelial CD98-TRPV4-NFkB cycle defined here. Galectin-3 appears to co-localise very closely with CD98 and β1-integrin, consistent with its putative role as a physiological binding partner. Increased recruitment of TGF-βRII and/or other galectin-3 ligands into a gal-3-fibrosome might then convert it into a checkpoint clustering both pro-inflammatory and pro-fibrotic pathways, with cross-talk allowing reinforcement of both processes.

## Materials and Methods

### *In vitro* studies

#### Cell models

Immortalised human type 1 alveolar epithelial TT1 cells were a kind gift from Prof. T. Tetley, National Heart and Lung Institute, Imperial University, London, UK. Human A549 cells used to model type 2 alveolar epithelial cells were obtained from ATCC-LCG, UK. They were cultured in conditions favouring Type 2 alveolar epithelial cell phenotypic features as described previously (49). Cell lysates were assessed using sodium dodecyl sulphate 4-12% Bis-Tris (v/v) gradient polyacrylamide gel electrophoresis (SDS-PAGE) probed by Western blot using relevant primary and secondary antibodies as indicated in Supplementary Table 1. The same antibodies were used for co-immunoprecipitation studies, with Protein G magnetic beads (DynaBeads, Invitrogen). Calu-3 cells were obtained by kind gift from the MRC Toxicology Unit. Cell viability was assessed by MTT assay and by acridine orange/DAPI (using a NucleoCounter NC-200, Chemometec, Denmark) staining to confirm that none of the treatments described had a toxic effect on the cells relative to controls.

### *Ex vivo* human tissue and culture models

All patient donors gave written informed consent and the study was approved by the National Research Ethics Service (references 04/Q2502/74, 07/MRE08/42, 08/H0406/18, and 10/H0402/12). All methods were performed in accordance with the relevant guidelines and regulations. Samples obtained were anonymised and coded before use. Human lung tissue from lung resections was harvested and cultured, and TGF-β1 stimulation administered, as described previously (50). LGALS3 and SLC3A2 mRNA responses were assessed in seven and nine samples respectively. The same *ex vivo* culture system was used to assess human lung cytokine responses to lipopolysaccharide (LPS, L5668, Sigma) stimulation by ELISA. Archival paraffin- and glycol methacrylate (GMA)-embedded human lung tissue from patients with IPF and non-fibrotic controls (healthy tissue from lung cancer resections) was obtained from blocks held at Leicester Institute of Lung Health/NIHR Leicester Biomedical Research Centre – Respiratory. Immunohistochemical studies to assess colocalisation of galectin-3, CD98hc, β1-integrin and E-cadherin were as previously described (51). Co-localisation of galectin-3, CD98hc and β1-integrin was also assessed in A549 cells by confocal microscopy and co-immunoprecipitation using previously described protocols (52) in the absence and presence of 10 ng/ml TGF-β1 stimulation or in titration conditions (0-10 ng/ml as stated).

Endogenous TGF-βRII was probed using by two monoclonal antibodies (mAbs: sc-17792, Santa Cruz, and PA5-36115, ThermoFisher). Neither detected endogenous protein in our immunofluorescence or western blot studies.

### Proximity ligation assay

Proximity ligation assay (PLA (53)) was performed using the In Situ Red Starter Kit Mouse/Rabbit (Duolink DUO92101, Sigma Aldrich). The conditions specified by the manufacturer’s instructions were optimized by incubating sections with primary antibodies overnight at 4℃ and increasing incubation time with probes and amplification reagents to 2 hours as previously described (54). Isotype controls for all primary and secondary antibodies confirmed the technical specificity of experimental findings.

### Mouse studies

CD98-*flox*ed mice were kindly provided by Prof. Mark Ginsberg (University of California, San Diego) and generated as described (55). Alveolar epithelial cells (AECs) from CD98-floxed mice were isolated as described previously (28). To induce recombination AECs were cultured with adenoviral-cre (MOI = 6) for 8h washed and cultured for a further 40 h prior to addition of TGF-β1.

The well-established murine intratracheal LPS instillation model was used to study acute PAMP-mediated lung injury (56). Seven week old wild-type male C57BL/6 mice (200-230 g, Charles River UK Ltd) were briefly anaesthetised by inhalation of 100% isoflurane liquid vapour (Centaur Services Ltd, UK) and LPS (or same volume PBS control) was administered intratracheally at 1 mg/ml (50 μl) and cynaropicrin (or same volume DMSO control) was injected intraperitoneally at 100 mg/kg (100 μl) at the same time. This was the highest dose used and showed the greatest experimental effects, without any evidence of associated toxicity at the animal or cellular level, consistent with previous reports (57). Mouse lungs were prepared as previously described (58). Four mice were studied per experimental time point, per condition. Following sacrifice, lungs were lavaged immediately. Excised lungs were then studied by immunohistochemical analysis for cytokine and cellular analyses.

Light micrographs were recorded at 10x magnification and analysed for the presence of neutrophils and macrophages using ImageJ software (National Institutes of Health, USA). Neutrophils were identified by staining with anti-neutrophil elastase antibody. Tissue was also stained for galectin-3 and CD98hc. Six representative images were analysed per treatment group and cell type. Field selections and quantitative assessments were made by an observer blinded to the experimental condition, as previously described (59).

### Enzyme-linked immunosorbent assay (ELISA)

ELISAs were set up as previously described (60). The following kits were used to detect different cytokines: Human IL-6 DuoSet, Human CXCL8/IL-8 DuoSet, Human TNF-α DuoSet, Mouse IL-6 DuoSet, Mouse CXCL2/MIP-2 DuoSet, Mouse Galectin-3 DuoSet (all from R&D Systems,UK).

### siRNA-mediated knockdown studies

TT1 and A549 cells were plated at 2 x 10^5^ cells/mL in 6-well plates, cultured for 18 h to reach 30-50% confluency, then transfected using oligofectamine transfection reagent (Thermo Fisher Scientific) with 20 μM small interfering siRNAs targeting the gene encoding human CD98hc, SLC3A2 (Thermo Fisher Scientific) or a non-targeting control construct.

### *In vitro* cyclic mechanical stretch (CMS) assay

TT1 and A549 cells were plated in 6-well collagen Bioflex culture plates (Flexcell International Corporation, Germany). CMS was applied using a Flexcell FX-4000T stretch machine (Flexcell International Corporation, Germany) programmed to stretch between specified strains (15% CMS or 21.8% CMS) at a specified frequency (0.2 Hz or 1 Hz) at either 2 or 24 h timepoints. When cells were co-treated with LPS, 4F2 mAb, GSK1016790A, HC-067047, GSK2193874 or JSH-23, these were applied 24 h prior to stretch. Ethylene glycol-bis(β-aminoethyl ether)- N,N,N’,N’-tetraacetic acid (EGTA) treatment was applied for 1 h prior to stretch when required. Cell media were refreshed before stretching. Optimisation assays demonstrated that the measured cytokine responses increased with increasing cyclic strain amplitude, and duration, but not frequency in the conditions used. Unless otherwise stated CMS conditions presented here are 0-15% strain, 2 h, 1 Hz. Early experiments screening for likely intracellular pathways mediating the CMS response used settings of 0-30% CMS. Data from these studies, illustrated in Supplementary Data, were qualitatively robust as assessed by specific knockdown and inhibitor studies. However for quantitative comparison studies we did not exceed 21.8% CMS since equibiaxial strain cannot be guaranteed beyond this.

### Matrix stiffness studies

Polyacrylamide hydrogels were prepared to generate a range of matrix stiffness conditions as previously described (61). Amino-silanated coverslips (22 mm diameter) were prepared via precipitation of NaOH prior to reaction with 3-aminopropyltriethoxysilane, rinsing with water, treatment with 0.5% glutaraldehyde in PBS for 30 min, and allowed to air dry. Chloro-silanated glass slides were prepared by coating with dichlorodimethylsilane for 5 min. 40% acrylamide and 2% bis-acrylamide stock solutions in PBS were mixed to achieve the desired concentrations, prior to addition of 10% (w/v) ammonium persulfate (1:100 (v/v)) and tetramethylethylenediamine (1:1,000 v/v) to initiate polymerisation. The solution was pipetted onto the treated side of a chloro-silanated glass slide, and the amino-silanated coverslip placed on top. The gel was allowed polymerise for up to 30 min prior to removal of the glass slide and rinsing in PBS. Gels were treated with 0.2 mg ml^-1^ sulfo-SANPAH (Pierce, Fisher Scientific) and exposed to UV light for 10 min, rinsed with 50 mM HEPES, with overnight incubation with bovine collagen type I (Thermo Fisher Scientific) in 50 mM HEPES to achieve 10 μg.cm^-2^.

Gel stiffness was assessed using indentation. Gels were submerged in PBS and tested in compression using a 3 mm diameter flat-ended cylindrical indenter using an Instron 5967 materials testing machine fitted with a ±10 N load cell (resolution 0.01 mN; Instron, High Wycombe, UK). Gel thickness was assessed by the application of a 0.5 mN pre-load. Gels were indented to 20% strain at 10% s^-1^ followed by a static hold of 120 s. Gel modulus was calculated from load-displacement data using a mathematical model derived from the solution of Sneddon for the axisymmetric Boussinesq problem (62, 63): *E = (S/2a)(1−ν^2^)*, where the indentation stiffness *S* is calculated from the tangent of the slope representing 15%-20% sample strain, *a* is the indenter radius, and Poisson’s ratio, *ν*, is assumed to be 0.5 (64).

### Intracellular Ca^2+^ assays

A549 cells were plated into 96-well blackbottom assay plates (Corning Incorporated, UK) and treated with combinations of GSK1016790A, HC-067047, JSH-23, the 4F2 mAb and CD98hc siRNA for 24 h, and EGTA for 2 h. Hypotonicity-induced stretch was induced by replacing the media with a mixture of 85% (v/v) DMEM. Measurement of intracellular Ca^2+^ influx was conducted using the Fluo-4 DirectTM Ca^2+^ Assay Kit (Thermo Fisher Scientific). Fluorescence was measured using a SpectraMax Plus 384 microplate reader (excitation 495 nm, emission 516 nm) with SoftMax Pro Software version 6 (both from Molecular Devices).

### Statistical analyses

For individual comparisons of clear pre-test biological interest where experimental differences involved a single variable, parametric data were assessed using paired t-tests and non-parametric data using Wilcoxon signed-rank test. For comparisons across multiple groups and timepoints, two-way ANOVA was followed by Sidak’s test for individual comparisons. Data analysis was performed using GraphPad Prism software version 6 (GraphPad Software, Inc., USA). Data are presented as the mean ± standard error of the mean (SEM) unless otherwise stated.

### Preparation and immunoprecipitation of recombinant SARS-CoV-2 spike protein receptor binding domain

Codon-optimized cDNA encoding the receptor binding domain of the SARS-CoV-2 spike protein (RBD, residues 319-541), together with an N-terminal CD5 secretory leader signal and C-terminal hexahistidine-tag was a kind gift from Prof Nick Brindle (Department of Molecular and Cellular Biology, University of Leicester). It was cloned into a pcDNA3.1 plasmid, transfected into suspension HEK293 cells and the secreted protein purified by affinity chromatography using a nickel sepharose column. This was stored in Tris-buffered saline solution. Following stimulation of cells with Spike RBD, its co-immunoprecipitation with CD98 and galectin-3 was assessed. To these ends, anti-Spike RBD antibody (Sino Biological, Catalog Number: 40150-R007) and Protein A/G magnetic beads (Thermo Scientific, MA, USA) were used according to the manufacturers’ protocols.

Stated p-values for all comparisons of interest are obtained from paired (same starting sample) or unpaired (different starting samples) t-testing unless otherwise stated. Mean +/- standard deviation indicated unless otherwise stated. * p<0.05, ** p<0.01, *** p<0.001, ns or quoted p-value: non-significant relative to p=0.05 boundary. All p-values assessed by t-tests given together in Supplementary Table 2.

## Acknowledgments

The authors wish to thank Prof Gisli Jenkins, Dr Alison John (Imperial College London) and Prof Catherine Hawrylowicz (King’s College London) for helpful discussions of the research as it progressed.

We thank the Advanced Imaging Facility (RRID:SCR_020967) at the University of Leicester for support.

Research in this work was supported by funding from the British Lung Foundation and Asthma+Lung UK, MRC (UK) (including Confidence in Concept award administered by Leicester Drug and Diagnostics Discovery (LD3) scheme), BBSRC, the Wellcome Trust, Guy’s & St Thomas’ Charity. Funding for the mechanobiology facilities used in this study came from EPSRC and Queen Mary strategic equipment funding.

The work was supported in part by the National Institute for Health Research (NIHR) Leicester Biomedical Research Centre - Respiratory. The views expressed are those of the authors and not necessarily those of the funders or affiliated bodies.

PB has received research funding from Genentech, Inc. via the University Hospitals of Leicester NHS Trust and from DEShawResearch via the University of Leicester, consultancies for Boehringer-Ingelheim (BI) and Genentech via the University of Leicester. Support to attend scientific meetings from Chiesi, Teva and Sanofi Genzyme. AM is a shareholder and employee of Galecto, Inc. BG has received consultancy fees from Vertex, and GSK, and research support in kind from Galecto, Inc. MMK currently has industry funding from Emulate for the Queen Mary+Emulate Organs-on-chips Centre and related research activity (https://www.cpm.qmul.ac.uk/emulate/).

## References

1. O’Dwyer DN, Ashley SL, Moore BB. Influences of innate immunity, autophagy, and fibroblast activation in the pathogenesis of lung fibrosis. Am J Physiol Lung Cell Mol Physiol. 2016;311(3):L590–601.

2. Xu Y, Mizuno T, Sridharan A, Du Y, Guo M, Tang J, Wikenheiser-Brokamp KA, Pearl AT, Funari VA, Gokey JJ, Stripp BR, Whitsett JA. Single-cell RNA sequencing identifies diverse roles of epithelial cells in idiopathic pulmonary fibrosis. JCI Insight. 2016;1(20):e90558.

3. Idiopathic Pulmonary Fibrosis Clinical Research N, Raghu G, Anstrom KJ, King TE, Jr., Lasky JA, Martinez FJ. Prednisone, azathioprine, and N-acetylcysteine for pulmonary fibrosis. N Engl J Med. 2012;366(21):1968–77.

4. Flaherty KR, Travis WD, Colby TV, Toews GB, Kazerooni EA, Gross BH, Jain A, Strawderman RL, Flint A, Lynch JP, Martinez FJ. Histopathologic variability in usual and nonspecific interstitial pneumonias. Am J Respir Crit Care Med. 2001;164(9):1722–7.

5. Monaghan H, Wells AU, Colby TV, du Bois RM, Hansell DM, Nicholson AG. Prognostic implications of histologic patterns in multiple surgical lung biopsies from patients with idiopathic interstitial pneumonias. Chest. 2004;125(2):522–6.

6. Myers JL. Hypersensitivity pneumonia: the role of lung biopsy in diagnosis and management. Mod Pathol. 2012;25 Suppl 1:S58–67.

7. Wang P, Jones KD, Urisman A, Elicker BM, Urbania T, Johannson KA, Assayyag D, Lee J, Wolters PJ, Collard HR, Koth LL. Pathologic Findings and Prognosis in a Large Prospective Cohort of Chronic Hypersensitivity Pneumonitis. Chest. 2017;152(3):502–9.

8. Singh N, Varghese J, England BR, Solomon JJ, Michaud K, Mikuls TR, Healy HS, Kimpston EM, Schweizer ML. Impact of the pattern of interstitial lung disease on mortality in rheumatoid arthritis: A systematic literature review and meta-analysis. Semin Arthritis Rheum. 2019.

9. Kelly BT, Moua T. Overlap of interstitial pneumonia with autoimmune features with undifferentiated connective tissue disease and contribution of UIP to mortality. Respirology. 2018;23(6):600–5.

10. King TE, Jr., Bradford WZ, Castro-Bernardini S, Fagan EA, Glaspole I, Glassberg MK, Gorina E, Hopkins PM, Kardatzke D, Lancaster L, Lederer DJ, Nathan SD, Pereira CA, Sahn SA, Sussman R, Swigris JJ, Noble PW, ASCEND Study Group. A phase 3 trial of pirfenidone in patients with idiopathic pulmonary fibrosis. N Engl J Med. 2014;370(22):2083–92.

11. Richeldi L, du Bois RM, Raghu G, Azuma A, Brown KK, Costabel U, Cottin V, Flaherty KR, Hansell DM, Inoue Y, Kim DS, Kolb M, Nicholson AG, Noble PW, Selman M, Taniguchi H, Brun M, Maulf FL, Girard M, Stowasser S, Schlenker-Herceg R, Disse B, Collard HR, INPULSIS Trial Investigators. Efficacy and safety of nintedanib in idiopathic pulmonary fibrosis. N Engl J Med. 2014;370(22):2071–82.

12. Wells AU, Flaherty KR, Brown KK, Inoue Y, Devaraj A, Richeldi L, Moua T, Crestani B, Wuyts WA, Stowasser S, Quaresma M, Goeldner RG, Schlenker-Herceg R, Kolb M, INBUILD Trial Investigators. Nintedanib in patients with progressive fibrosing interstitial lung diseases-subgroup analyses by interstitial lung disease diagnosis in the INBUILD trial: a randomised, double-blind, placebo-controlled, parallel-group trial. Lancet Respir Med. 2020;8(5):453–60.

13. Kuwana M, Ogura T, Makino S, Homma S, Kondoh Y, Saito A, Ugai H, Gahlemann M, Takehara K, Azuma A. Nintedanib in patients with systemic sclerosis-associated interstitial lung disease: A Japanese population analysis of the SENSCIS trial. Mod Rheumatol. 2020:1–10.

14. Chambers RC, Mercer PF. Mechanisms of alveolar epithelial injury, repair, and fibrosis. Annals of the American Thoracic Society. 2015;12 Suppl 1:S16–20.

15. Bagnato G, Harari S. Cellular interactions in the pathogenesis of interstitial lung diseases. Eur Respir Rev. 2015;24(135):102–14.

16. Haak AJ, Tan Q, Tschumperlin DJ. Matrix biomechanics and dynamics in pulmonary fibrosis. Matrix Biol. 2018;73:64–76.

17. Froese AR, Shimbori C, Bellaye PS, Inman M, Obex S, Fatima S, Jenkins G, Gauldie J, Ask K, Kolb M. Stretch-induced Activation of Transforming Growth Factor-beta1 in Pulmonary Fibrosis. Am J Respir Crit Care Med. 2016;194(1):84–96.

18. Mallick S. Outcome of patients with idiopathic pulmonary fibrosis (IPF) ventilated in intensive care unit. Respir Med. 2008;102(10):1355–9.

19. Mura M, Porretta MA, Bargagli E, Sergiacomi G, Zompatori M, Sverzellati N, Taglieri A, Mezzasalma F, Rottoli P, Saltini C, Rogliani P. Predicting survival in newly diagnosed idiopathic pulmonary fibrosis: a 3-year prospective study. Eur Respir J. 2012;40(1):101–9.

20. Simon-Blancal V, Freynet O, Nunes H, Bouvry D, Naggara N, Brillet PY, Denis D, Cohen Y, Vincent F, Valeyre D, Naccache JM. Acute exacerbation of idiopathic pulmonary fibrosis: outcome and prognostic factors. Respiration; international review of thoracic diseases. 2012;83(1):28–35.

21. Natsuizaka M, Chiba H, Kuronuma K, Otsuka M, Kudo K, Mori M, Bando M, Sugiyama Y, Takahashi H. Epidemiologic survey of Japanese patients with idiopathic pulmonary fibrosis and investigation of ethnic differences. Am J Respir Crit Care Med. 2014;190(7):773–9.

22. Watanabe T, Minezawa T, Hasegawa M, Goto Y, Okamura T, Sakakibara Y, Niwa Y, Kato A, Hayashi M, Isogai S, Kondo M, Yamamoto N, Hashimoto N, Imaizumi K. Prognosis of pulmonary fibrosis presenting with a usual interstitial pneumonia pattern on computed tomography in patients with myeloperoxidase anti-neutrophil cytoplasmic antibody-related nephritis: a retrospective single-center study. BMC Pulm Med. 2019;19(1):194.

23. Albert RK, Smith B, Perlman CE, Schwartz DA. Is Progression of Pulmonary Fibrosis due to Ventilation-induced Lung Injury? Am J Respir Crit Care Med. 2019;200(2):140–51.

24. Molyneaux PL, Maher TM. The role of infection in the pathogenesis of idiopathic pulmonary fibrosis. Eur Respir Rev. 2013;22(129):376–81.

25. Tiitto L, Bloigu R, Heiskanen U, Paakko P, Kinnula VL, Kaarteenaho-Wiik R. Relationship between histopathological features and the course of idiopathic pulmonary fibrosis/usual interstitial pneumonia. Thorax. 2006;61(12):1091–5.

26. Dumic J, Dabelic S, Flogel M. Galectin-3: an open-ended story. Biochim Biophys Acta. 2006;1760(4):616–35.

27. Lin YH, Qiu DC, Chang WH, Yeh YQ, Jeng US, Liu FT, Huang JR. The intrinsically disordered N-terminal domain of galectin-3 dynamically mediates multisite self-association of the protein through fuzzy interactions. J Biol Chem. 2017;292(43):17845–56.

28. Mackinnon AC, Gibbons MA, Farnworth SL, Leffler H, Nilsson UJ, Delaine T, Simpson AJ, Forbes SJ, Hirani N, Gauldie J, Sethi T. Regulation of transforming growth factor-β1-driven lung fibrosis by galectin-3. Am J Resp Crit Care Med. 2012;185:537–46.

29. Slack RJ, Hirani N, Gibbons MA, Simpson AJ, Ford P, Leffler H, Nilsson UJ, Sethi T, Pedersen A, Schambye H, Maher TM, MacKinnon AC. Translational pharmacology of TD139, an inhaled small molecule galectin-3 (Gal-3) inhibitor for the treatment of idiopathic pulmonary fibrosis (IPF). FASEB journal : official publication of the Federation of American Societies for Experimental Biology. 2020;34(S1).

30. Chan YC, Lin HY, Tu Z, Kuo YH, Hsu SD, Lin CH. Dissecting the Structure-Activity Relationship of Galectin-Ligand Interactions. Int J Mol Sci. 2018;19(2).

31. Fotiadis D, Kanai Y, Palacin M. The SLC3 and SLC7 families of amino acid transporters. Molecular aspects of medicine. 2013;34(2-3):139–58.

32. Prager GW, Feral CC, Kim C, Han J, Ginsberg MH. CD98hc (SLC3A2) interaction with the integrin beta subunit cytoplasmic domain mediates adhesive signaling. J Biol Chem. 2007;282(33):24477–84.

33. Veettil MV, Sadagopan S, Sharma-Walia N, Wang FZ, Raghu H, Varga L, Chandran B. Kaposi’s sarcoma-associated herpesvirus forms a multimolecular complex of integrins (alphaVbeta5, alphaVbeta3, and alpha3beta1) and CD98-xCT during infection of human dermal microvascular endothelial cells, and CD98-xCT is essential for the postentry stage of infection. J Virol. 2008;82(24):12126–44.

34. Fenczik CA, Sethi T, Ramos JW, Hughes PE, Ginsberg MH. Complementation of dominant suppression implicates CD98 in integrin activation. Nature. 1997;390(6655):81–5.

35. Rintoul RC, Buttery RC, Mackinnon AC, Wong WS, Mosher D, Haslett C, Sethi T. Cross-linking CD98 promotes integrin-like signaling and anchorage-independent growth. Mol Biol Cell. 2002;13(8):2841–52.

36. MacKinnon AC, Farnworth SL, Hodkinson PS, Henderson NC, Atkinson KM, Leffler H, Nilsson UJ, Haslett C, Forbes SJ, Sethi T. Regulation of alternative macrophage activation by galectin-3. J Immunol. 2008;180(4):2650–8.

37. Dalton P, Christian HC, Redman CW, Sargent IL, Boyd CA. Membrane trafficking of CD98 and its ligand galectin 3 in BeWo cells--implication for placental cell fusion. FEBS J. 2007;274(11):2715–27.

38. Xue FM, Zhang HP, Hao HJ, Shi ZY, Zhou C, Feng B, Yang PC. CD98 positive eosinophils contribute to T helper 1 pattern inflammation. PLoS One. 2012;7(12):e51830.

39. Nguyen HT, Dalmasso G, Torkvist L, Halfvarson J, Yan Y, Laroui H, Shmerling D, Tallone T, D’Amato M, Sitaraman SV, Merlin D. CD98 expression modulates intestinal homeostasis, inflammation, and colitis-associated cancer in mice. J Clin Invest. 2011;121(5):1733–47.

40. Case LB, Waterman CM. Integration of actin dynamics and cell adhesion by a three-dimensional, mechanosensitive molecular clutch. Nat Cell Biol. 2015;17(8):955–63.

41. Bachmann M, Kukkurainen S, Hytonen VP, Wehrle-Haller B. Cell Adhesion by Integrins. Physiol Rev. 2019;99(4):1655–99.

42. Shi M, Zhu J, Wang R, Chen X, Mi L, Walz T, Springer TA. Latent TGF-beta structure and activation. Nature. 2011;474(7351):343–9.

43. Jenkins G. The role of proteases in transforming growth factor-beta activation. Int J Biochem Cell Biol. 2008;40(6-7):1068–78.

44. Danen EH, Sonneveld P, Brakebusch C, Fassler R, Sonnenberg A. The fibronectin-binding integrins alpha5beta1 and alphavbeta3 differentially modulate RhoA-GTP loading, organization of cell matrix adhesions, and fibronectin fibrillogenesis. The Journal of cell biology. 2002;159(6):1071–86.

45. Munger JS, Huang X, Kawakatsu H, Griffiths MJ, Dalton SL, Wu J, Pittet JF, Kaminski N, Garat C, Matthay MA, Rifkin DB, Sheppard D. The integrin alpha v beta 6 binds and activates latent TGF beta 1: a mechanism for regulating pulmonary inflammation and fibrosis. Cell. 1999;96(3):319–28.

46. Jenkins RG, Su X, Su G, Scotton CJ, Camerer E, Laurent GJ, Davis GE, Chambers RC, Matthay MA, Sheppard D. Ligation of protease-activated receptor 1 enhances alpha(v)beta6 integrin-dependent TGF-beta activation and promotes acute lung injury. J Clin Invest. 2006;116(6):1606–14.

47. Sigrist CJ, Bridge A, Le Mercier P. A potential role for integrins in host cell entry by SARS-CoV-2. Antiviral Res. 2020;177:104759.

48. Calver J, Joseph C, John AE, Organ L, Fainberg H, Porte J, Mukhopadhyay S, Barton L, Stroberg E, Duval E, Copin M, Poissy J, Streinestrel K, Tatler AL, Jenkins G. S31 The novel coronavirus SARS-CoV-2 binds RGD integrins and upregulates avb3 integrins in Covid-19 infected lungs Thorax. 2021;76:A22–A3.

49. Cooper JR, Abdullatif MB, Burnett EC, Kempsell KE, Conforti F, Tolley H, Collins JE, Davies DE. Long Term Culture of the A549 Cancer Cell Line Promotes Multilamellar Body Formation and Differentiation towards an Alveolar Type II Pneumocyte Phenotype. PLoS One. 2016;11(10):e0164438.

50. Roach KM, Sutcliffe A, Matthews L, Elliott G, Newby C, Amrani Y, Bradding P. A model of human lung fibrogenesis for the assessment of anti-fibrotic strategies in idiopathic pulmonary fibrosis. Sci Rep. 2018;8(1):342.

51. Bradding P, Feather IH, Howarth PH, Mueller R, Roberts JA, Britten K, Bews JP, Hunt TC, Okayama Y, Heusser CH, Bullock GR, Church MK, Holgate ST. Interleukin 4 is localized to and released by human mast cells. J Exp Med. 1992;176(5):1381–6.

52. Stylianou P, Clark K, Gooptu B, Smallwood D, Brightling CE, Amrani Y, Roach KM, Bradding P. Tensin1 expression and function in chronic obstructive pulmonary disease. Sci Rep. 2019;9(1):18942.

53. Soderberg O, Gullberg M, Jarvius M, Ridderstrale K, Leuchowius KJ, Jarvius J, Wester K, Hydbring P, Bahram F, Larsson LG, Landegren U. Direct observation of individual endogenous protein complexes in situ by proximity ligation. Nat Methods. 2006;3(12):995–1000.

54. Trifilieff P, Rives ML, Urizar E, Piskorowski RA, Vishwasrao HD, Castrillon J, Schmauss C, Slättman M, Gullberg M, Javitch JA. Detection of antigen interactions ex vivo by proximity ligation assay: endogenous dopamine D2-adenosine A2A receptor complexes in the striatum. Biotechniques. 2011;51(2):111–8.

55. Cantor J, Slepak M, Ege N, Chang JT, Ginsberg MH. Loss of T cell CD98 H chain specifically ablates T cell clonal expansion and protects from autoimmunity. J Immunol. 2011;187:851–60.

56. Matute-Bello G, Downey G, Moore BB, Groshong SD, Matthay MA, Slutsky AS, Kuebler WM, Acute Lung Injury in Animals Study Group. An official American Thoracic Society workshop report: features and measurements of experimental acute lung injury in animals. Am J Respir Cell Mol Biol. 2011;44(5):725–38.

57. da Silva CF, Batista Dda G, De Araujo JS, Batista MM, Lionel J, de Souza EM, Hammer ER, da Silva PB, Mieri MD, Adams M, Zimmermann S, Hamburger M, Brun R, Schühly W, Soreiro MNC. Activities of psilostachyin A and cynaropicrin against Trypanosoma cruzi in vitro and in vivo. Antimicrobial agents and chemotherapy. 2013;57(11):5307–14.

58. Cuello AC, editor. Immunohistochemistry II. New York: John Wiley & Sons; 1993.

59. Cleary SJ, Hobbs C, Amison RT, Arnold S, O’Shaughnessy BG, Lefrançais E, Mallavia B, Looney MR, Page CP, Pitchford SC. LPS-induced lung platelet recruitment occurs independently from neutrophils, PSGL-1, and P-selectin. Am J Resp Cell Mol Biol. 2019;61:232–43.

60. Schmidt SD, Mazzella MJ, Nixon RA, Mathews PM. Abeta measurement by enzyme-linked immunosorbent assay. Methods Mol Biol. 2012;849:507–27.

61. Tse JR, Engler AJ. Preparation of hydrogel substrates with tunable mechanical properties. Curr Protoc Cell Biol. 2010;Chapter 10:Unit 10 6.

62. Sneddon IN. The relation between load and penetration in the axisymmetric boussinesq problem for a punch of arbitrary profile. International Journal of Engineering Science. 1965;3(1):47–57.

63. Delaine-Smith RM, Burney S, Balkwill FR, Knight MM. Experimental validation of a flat punch indentation methodology calibrated against unconfined compression tests for determination of soft tissue biomechanics. J Mech Behav Biomed Mater. 2016;60:401–15.

64. Takigawa T, Morino Y, Urayama K, Masuda T. Poisson’s ratio of polyacrylamide (PAAm) gels. Polymer Gels and Networks. 1996;4(1):1–5.

65. Cho JY, Kim AR, Joo HG, Kim BH, Rhee MH, Yoo ES, Katz DR, Chain BM, Jung, JH. Cynaropicrin, a sesquiterpene lactone, as a new strong regulator of CD29 and CD98 functions. Biochemical and biophysical research communications. 2004;313(4):954–61.

66. Papiris SA, Tomos IP, Karakatsani A, Spathis A, Korbila I, Analitis A, Kolilekas L, Kagouridis K, Loukides S, Karakitsos P, Manali ED. High levels of IL-6 and IL-8 characterize early-on idiopathic pulmonary fibrosis acute exacerbations. Cytokine. 2018;102:168–72.

67. Schupp JC, Binder H, Jager B, Cillis G, Zissel G, Muller-Quernheim J, Prasse A. Macrophage activation in acute exacerbation of idiopathic pulmonary fibrosis. PLoS One. 2015;10(1):e0116775.

68. Rahaman SO, Grove LM, Paruchuri S, Southern BD, Abraham S, Niese KA, Scheraga RG, Ghosh S, Thodeti CK, Zhang DX, Moran MM, Schilling WP, Tschumperlin DJ, Olman MA. TRPV4 mediates myofibroblast differentiation and pulmonary fibrosis in mice. J Clin Invest. 2014;124(12):5225–38.

69. Matthews BD, Thodeti CK, Tytell JD, Mammoto A, Overby DR, Ingber DE. Ultra-rapid activation of TRPV4 ion channels by mechanical forces applied to cell surface beta1 integrins. Integr Biol (Camb). 2010;2(9):435–42.

70. Loukin S, Zhou X, Su Z, Saimi Y, Kung C. Wild-type and brachyolmia-causing mutant TRPV4 channels respond directly to stretch force. J Biol Chem. 2010;285:27176–81.

71. Li W, Moore MJ, Vasilieva N, Sui J, Wong SK, Berne MA, Somasundaran M, Sullivan JL, Luzuriaga K, Greenough TC, Choe H, Farzan M. Angiotensin-converting enzyme 2 is a functional receptor for the SARS coronavirus. Nature. 2003;426:450–4.

72. Sungnak W, Huang N, Becavin C, Berg M, Queen R, Litvinukova M, Talavera-Lόpez C, Maatz H, Reichart D, Sampaziotis F, Worlock KB, Yoshida M, Barnes JL, HCA Lung Biological Network. SARS-CoV-2 entry factors are highly expressed in nasal epithelial cells together with innate immune genes. Nature Medicine. 2020;26:681–7.

73. Bezara MO, Thurman A, Pezzulo A, Leidinger MR, Klesney-Tait JA, Karp PH, Tan P, Wohlford-Lenane C, Jr PBM, Meyerholz DK. Heterogeneous expression of the SARS-Coronavirus-2 receptor ACE2 in the human respiratory tract. EBioMedicine. 2020;60(102976).

74. Munger JS, Harpel JG, Giancotti FG, Rifkin DB. Interactions between growth factors and integrins: latent forms of transforming growth factor-β are ligands for the integrin αvβ1. Mol Biol Cell. 1998;9:2627–38.

75. Group RC. Tocilizumab in patients admitted to hospital with COVID-19 (RECOVERY): a randomised, controlled, open-label, platform trial. Lancet. 2021;397:1637–45.

76. Del Valle DM, Kim-Schulze S, Huang HH, Beckmann ND, Nirenberg S, Wang B, Lavin Y, Swartz TH, Madduri D, Stock A, Marron, TU, Xie H, Patel M, Tuballes K, Oekelen OV, Rahman A, Kovatch P, Aberg JA, Schadt E, Jagannath S, Mazumdar M, Charney AW, Firpo-Betancourt A, Mendu DR, Jhang J, Reich D, Sigel K, Cordon-Cardo C, Feldmann M, Parekh S, Merad M, Gnjatic S. An inflammatory cytokine signature predicts COVID-19 severity and survival. Nature Medicine. 2020;26:1636–43.

77. Li L, Li J, Gao M, Fan H, Wang Y, Xu X, Chen C, Liu J, Kim J, Aliyari R, Zhang J, Jin Y, Li X, Ma F, Shi M, Cheng G, Yang H. Interleukin-8 as a biomarker for disease prognosis of coronavirus disease-2019 patients. Frontiers in immunology. 2020;11:602395.

78. Laing AG, Lorenc A, del Molino del Barrio I, Das A, Fish M, Monin L, Muńoz-Ruiz M, McKenzie DR, Hayday TS, Francos-Quijorna I, Kamdar S, Joseph M, Davies D, Davis R, Jennings A, Zlatareva I, Vantourout P, Wu Y, Sofra V, Cano F, Greco M, Theodoridis E, Freedman JD, Gee S, Chan JNE, Ryan S, Bugallo-Blanco E, Peterson P, Kisand K, Haljasmägi L, Chadli L, Moingeon P, Martinez L, Merrick B, Bisnauthsing K, Brooks K, Ibrahim MAA, Mason J, Gomez FL, Babalola K, Abdul-Jawad S, Cason J, Mant C, Seow J, Graham C, Doores KJ, Rosa FD, Edgeworth J, Shankar-Hari M, Hayday AC. A dynamic COVID-19 immune signature includes associations with poor prognosis. Nature Medicine. 2020;26:1623–35.

79. Patra T, Meyer K, Geerling L, Isabell TS, Hoft DF, Brien J, Pinto AK, Ray RB, Ray R. SARS-CoV-2 spike protein promotes IL-6 trans-signaling by activation of angiotensin II receptor signaling in epithelial cells. PLoS Pathogens. 2020;16:e1009128.

80. Ren X, Glende J, Al-Falah M, De Vries V, Schwegmann-Wessels C, Qu X, Tan L, Tscherning T, Deng H, Naim HY, Herrler G. Analysis of ACE2 in polarized epithelial cells: surface expression and function as receptor for severe acute respiratory syndrome-associated coronavirus. Journal of General Virology. 2006;87:1691–5.

81. Merad M, Martin JC. Pathological inflammation in patients with COVID-19: a key role for monocytes and macrophages. Nature Reviews Immunology. 2020;20:355–62.

82. Nabi IR, Shankar J, Dennis JW. The galectin lattice at a glance. J Cell Sci. 2015;128(13):2213–9.

83. Argueso P, Guzman-Aranguez A, Mantelli F, Cao Z, Ricciuto J, Panjwani N. Association of cell surface mucins with galectin-3 contributes to the ocular surface epithelial barrier. J Biol Chem. 2009;284(34):23037–45.

84. Muramatsu T. Basigin (CD147), a multifunctional transmembrane glycoprotein with various binding partners. J Biochem. 2016;159(5):481–90.

85. Mole S, Harry A, Fowler A, Hotee S, Warburton J, Waite S, Beerahee M, Behm D, Badorrek P, Muller M, Faulenbach C, Lazaar A, Hohfield JM. A segmental LPS challenge study to investigate the pharmacodynamics of a TRPV4 antagonist (GSK2798745) in healthy participants. Thorax. 2019;74:A153.

86. Drake TM, Docherty AB, Harrison EM, Quint JK, Adamali H, Agnew S, Babu S, Barber CM, Barratt, S, Bendstrup E, Bianchi S, Villegas DC, Chaudhuri N, Chua F, Coker R, Chang W, Crawshaw A, Crowley LE, Dosanjh D, Fiddler CA, Forrest IA, George PM, Gibhbons MA, Groom K, Haney S, Hart SP, Heiden E, Henry M, Ho LP, Hoyles RK, Hutchinson J, Hurley K, Jones M, Jones S, Kokosi M, Kreuter M, MacKay LS, Mahendran S, Margaritopoulo G, Molina-Molina M, Molyneaux PL, O’Brien A, O’Reilly K, Packham A, Parfrey H, Poletti V, Porter JC, Renzoni E, Rivera-Ortega P, Russell AM, Saini G, Spencer LG, Stella GM, Stone H, Sturney S, Thickett D, Thillai M, Wallis T, Ward K, Wells AU, West A, Wickremasinghe M, Woodhead F, Hearson G, Howard L, Baillie JK, Openshaw PJM, Semple MG, Stewart I, Jenkins RG, ISARIC4C Investigators. Outcome of Hospitalization for COVID-19 in Patients with Interstitial Lung Disease. An International Multicenter Study. Am J Resp Crit Care Med. 2020;202:1656–65.

