## Supplemental material for "CD98 is critical for a conserved inflammatory response to diverse injury stimuli relevant to IPF exacerbations and COVID pneumonitis"

**Supplementary Figure Legends**

**Supp. Fig. 1**

**a) Model of a modular gal-3-fibrosome.** Illustrated by CD98:integrin complex potentially mediating acute inflammatory responses and TGF-βRII mediating chronic pro-fibrotic signalling by TGF-β1. There is potential for direct molecular cross-talk between these pathways through stabilised interactions within a physical complex.

**1b) Galectin-3, CD98, β1-integrin co-localise at the cellular level in IPF tissue**. Serial sections from same lung tissue blocks, stained for two or three of these proteins are displayed alongside each other horizontally for non-fibrotic control tissue (left) and IPF tissue (right). Epithelial staining with E-cadherin is shown in experiment displayed on the lowest row, in which a lower concentration of an anti-β1-integrin antibody was used based upon strength of staining in earlier experiments. **c) Membrane fractionation and Co-IP of galectin-3, CD98 and β1-integrin.** Membrane fractionation and Co-IP of galectin-3, CD98 and β1-integrin in A549 cells basally and upon stimulation with TGF-β1. SDS-PAGE probed with western blot analysis. The F1ATPase subunit ATP1A1 is a membrane-specific marker, whilst β-actin is a general cellular housekeeping protein. **d) Flow cytometry supports CD98 localisation to the A549 cell surface.** CD98hc protein distribution to the cell surface was assessed by comparing flow cytometry data in non-permeabilised and permeabilised cells. A representative trace is shown (left) and quantification of three biological repeats (centre). The similarity between these findings is strongly supportive of predominantly cell surface distribution in these conditions, whilst the specificity of the signal for CD98 is supported by findings when expression of this protein was silenced by siRNA knockdown (right).

**Supp. Fig. 2 Concentration titration of co-immunopreciptiation studies.**

SDS-PAGE probed by western blot, molecular weights (kDa) indicated for marker ladder. Similar co-immunoprecipitation of CD98 (upper panel) and $\beta$1-integrin (lower panel) was observed with stimulation across a 1-10 ng/ml TGF-$\beta$1 titration at 24 (left) and 72 (right) hours, suggesting that the phenomena studied were not qualitatively dependent upon specific TGF-$\beta$1 concentrations within this range.

**Supp. Fig. 3a) Day 1, 2, and 7 data for IL-6, IL-8 and TNF-α secretion responses to LPS -/+ cynaropicrin from *ex vivo* human lung tissue model.** Individual data from ELISA of culture supernatants, across the timecourse studied.

**b) *In vivo* mouse model cytokine responses to LPS stimulation -/+ cynaropicrin treatment.** IL-6, MIP-1α (murine paralog of IL-8) secretion responses to LPS stimulation were abrogated by cynaropicrin treatment (n=4) similarly to the responses in *ex vivo* human lung tissue.

**c)-i) Flow cytometry analysis of immune cell phenotypes within mouse BAL samples indicates no change in airway populations with cynaropicrin treatment at 24 hours.** Cell surface markers and general immune cell type correlates as indicated. PBS: phosphate buffered saline, Veh: DMSO vehicle control for cynaropicrin treatment, Cyn: Cynaropicrin. Full data distributions shown (left) with n=4 quantification plotted (right).

**Supp. Fig. 4a) CD98 mediates IL-6 and IL-8 but not TNF-α responses to LPS in TT1 cells.** ELISA data for IL-6, IL-8 and TNF-α responses to LPS stimulation, and effects of CD98 knockdown in TT1 cells. NC indicates non-coding siRNA control condition. (Equivalent data for A549 cells shown in Fig. 4a.)

**b) CD98 silencing and cynaropicrin have similar effects upon LPS responses *in vitro*.** Cynaropicrin treatment in cells recapitulates cynaropicrin behaviour observed *in vivo* and *ex vivo*, and is equivalent to CD98 knockdown.

**c) CD98 mediates IL-6 and IL-8 but not TNF-α responses to CMS in alveolar epithelial cells**. ELISA data for IL-6, IL-8 and TNF-α responses to CMS stimulation, and effects of CD98 knockdown, in A549 cells at 24 h and for TT1 cells at 2 and 24 h. NC indicates non-coding siRNA control condition. (Equivalent data for A549 cells at 2 h shown in Fig. 4b.)

**d) CD98 silencing and cynaropicrin have similar effects upon cytokine responses to CMS.** ELISA data for IL-6, IL-8 and TNF-α responses to CMS stimulation, with comparison of the effects of CD98 knockdown and pharmacological inhibition with cynaropicrin in TT1 and A549 cells.

**e) Dose:response relationships for IL-6 and IL-8 secretion with increasing matrix stiffness.** ELISA data for IL-6, IL-8 and TNF-α responses to CMS stimulation, with comparison of the effects of CD98 knockdown and pharmacological inhibition with cynaropicrin in TT1 and A549 cells.

**f) Additive effects of matrix stiffness, CMS and LPS on TT1 and A549 cells.** Addition of LPS to CMS and/or increased matrix stiffness conditions indicated small further incremental responses that did not consistently reach significance at a p-value $<0.05$, n=3, LPS and CMS conditions as used previously.

**Supp. Fig. 5 Lack of CD98-dependent responses to LPS or CMS stimulus in probed intracellular pathways other than those related to NF-κB signalling**

MAPK pathways did not show any CD98-dependent response to LPS (a) or CMS (b) stimulation. A p38 phosphorylation response was observed but this was not CD98-dependent. JNK phosphorylation was also assessed. However JNK levels did not appreciably change with stimulation or CD98 siRNA treatment, and no phospho-JNK signal was observed in any condition.

**Supp. Fig. 6a) TRPV4 inhibition and NF-kB inhibition reduce cellular CD98hc levels.** A549 cells treated with TRPV4 inhibitors (data shown for HC-067047, HC, with CMS, upper) or the NF-κB inhibitor JSH-23 (with LPS, lower) demonstrated reduced levels of CD98hc protein relative to control when assessed by SDS-PAGE/Western blot. The deficit was more marked in the absence of stimulus, and treatment with the cross-linking 4F2 mAb did not affect CD98 levels (shown here basally).

**b) NF-κB agonist stimulates IL-6 and IL-8 release from A549s in the absence of intracellular $Ca^{2+}$ influx.** Evidence of a positive feedback loop between CD98, TRPV4, $Ca^{2+}$ influx and NF-κB activation raised the possibility that NF-κB activity might not be the most direct element in mediating the associated release of IL-6 and IL-8. To test this, the preceding $Ca^{2+}$ influx step was blocked by EGTA treatment, preventing the CMS response involving these cytokines. Treatment with the NF-κB agonist PMA was then able to stimulate IL-6 and IL-8 release despite $Ca^{2+}$ chelation, supporting its role as the effector molecule within the feedback loop.

**Supp. Fig. 7 Proposed relationships between acute lung injury, COVID pneumonitis, and exacerbations and progression of fibrotic ILD e.g. IPF**

CD98:galectin-3 interactions provide a potential molecular basis for the transduction of acute injury responses to fibrotic responses. In IPF these appear enriched, predisposing to both development of acute inflammatory responses and potentially progressive sequelae. Where the primary insult is the acute injury, development of fibrosis and subsequent progressive behaviour may relate to the degree to which CD98:galectin-3 co-stabilisation occurs at the cell surface.

### Supp. Fig. 1A)

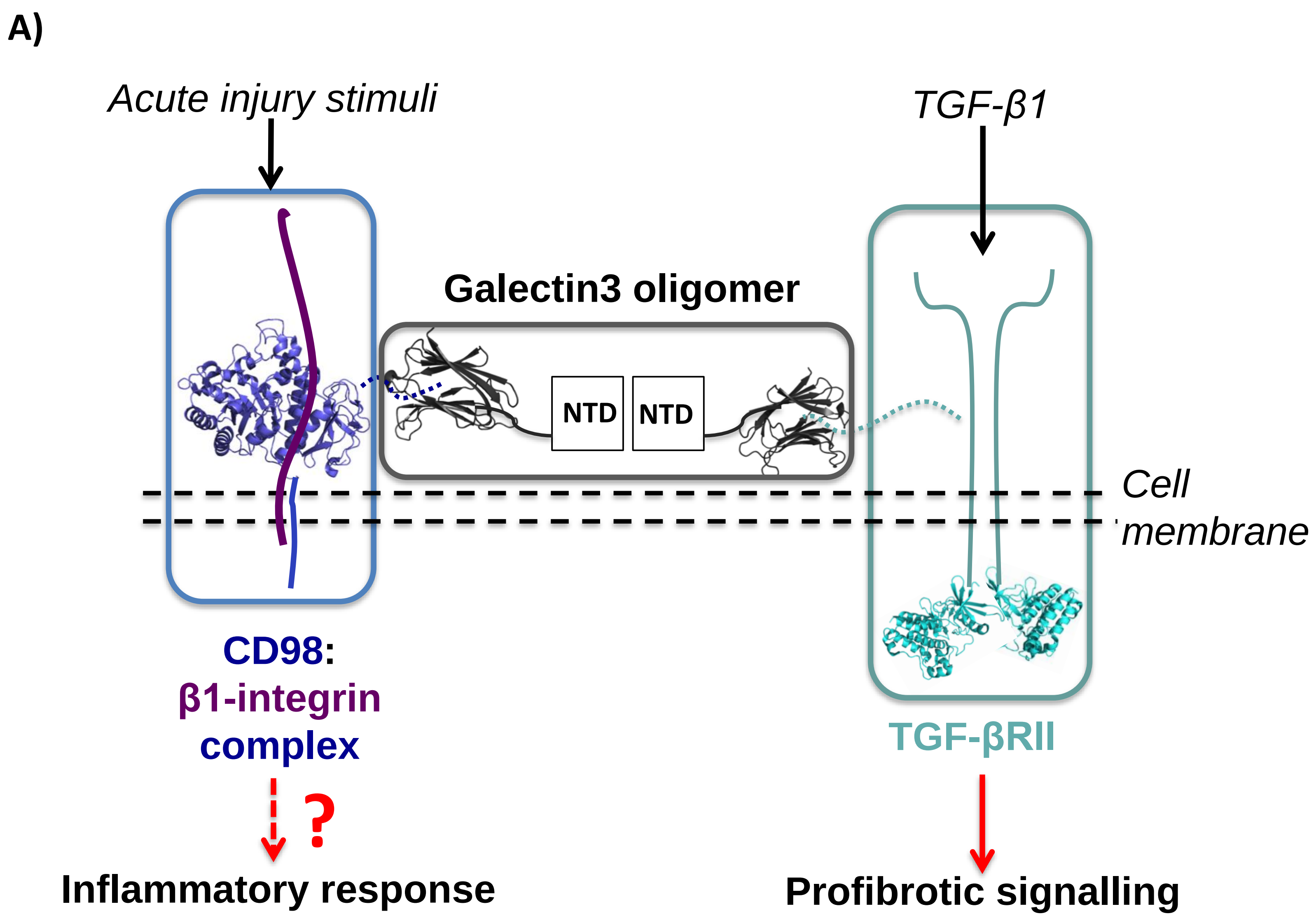
A)
Acute injury stimuli
TGF-β1
Galectin3 oligomer
NTD
NTD
Cell membrane
CD98:
β1-integrin
complex
TGF-βRII
?
Inflammatory response
Profibrotic signalling

### Supp. Fig. 1B)

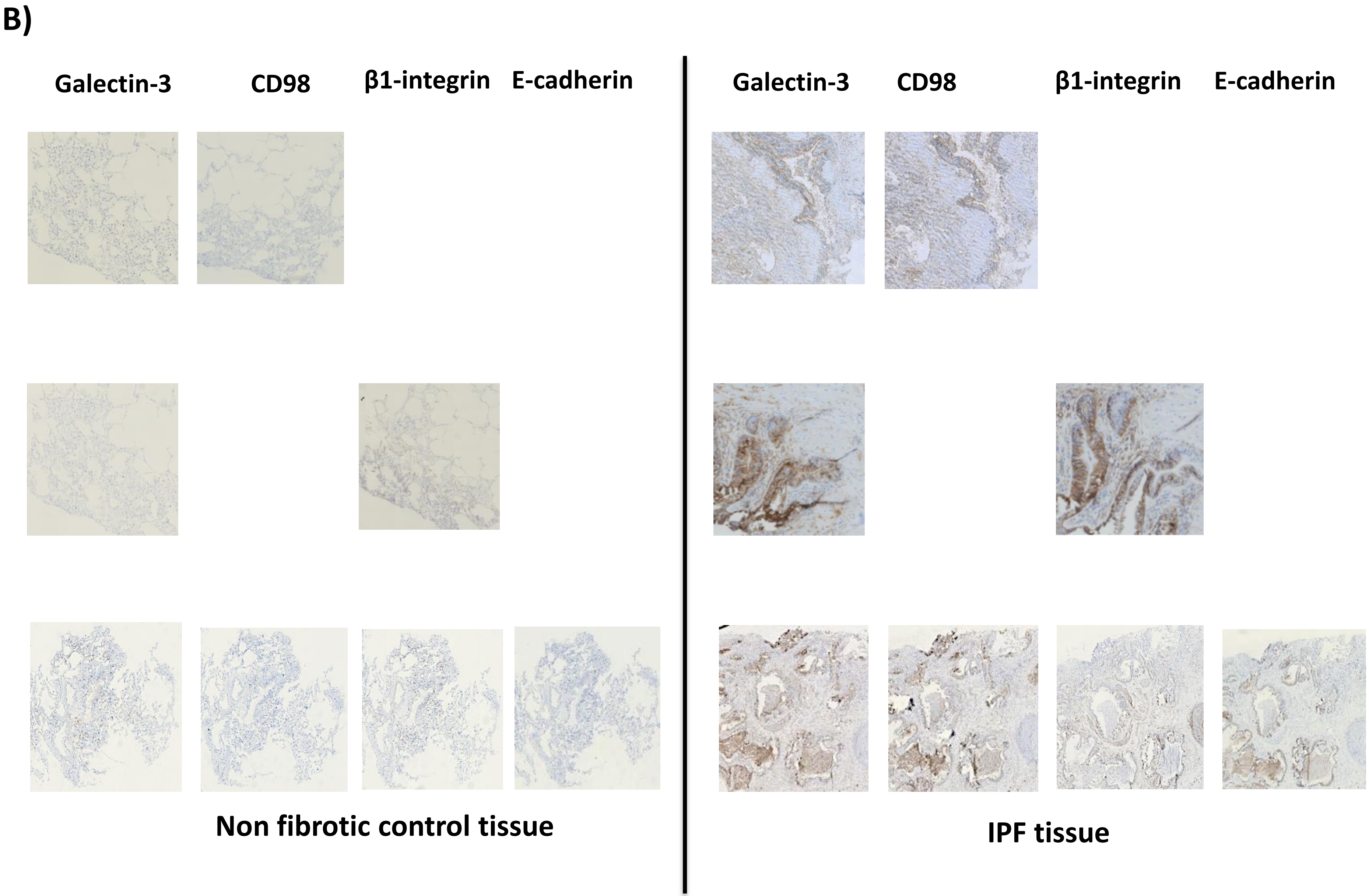
B)
Galectin-3
CD98
β1-integrin
E-cadherin
Galectin-3
CD98
β1-integrin
E-cadherin
Non fibrotic control tissue
IPF tissue

### Supp. Fig. 1C)

**C)**

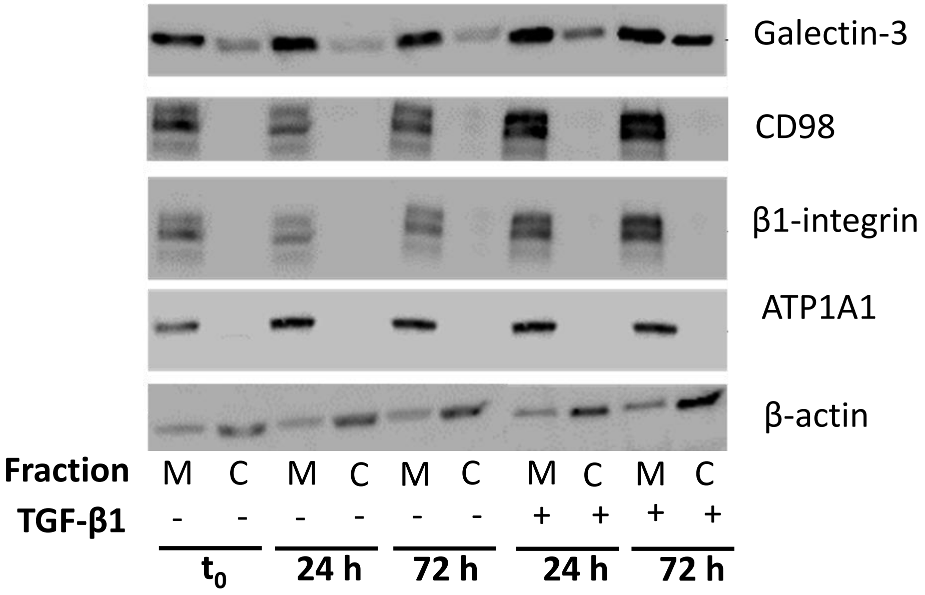
Galectin-3
CD98
β1-integrin
ATP1A1
β-actin
Fraction M C M C M C M C M C
TGF-β1 - - - - - - + + + +
t0 24 h 72 h 24 h 72 h

### Supp. Fig. 1D)

**D)**

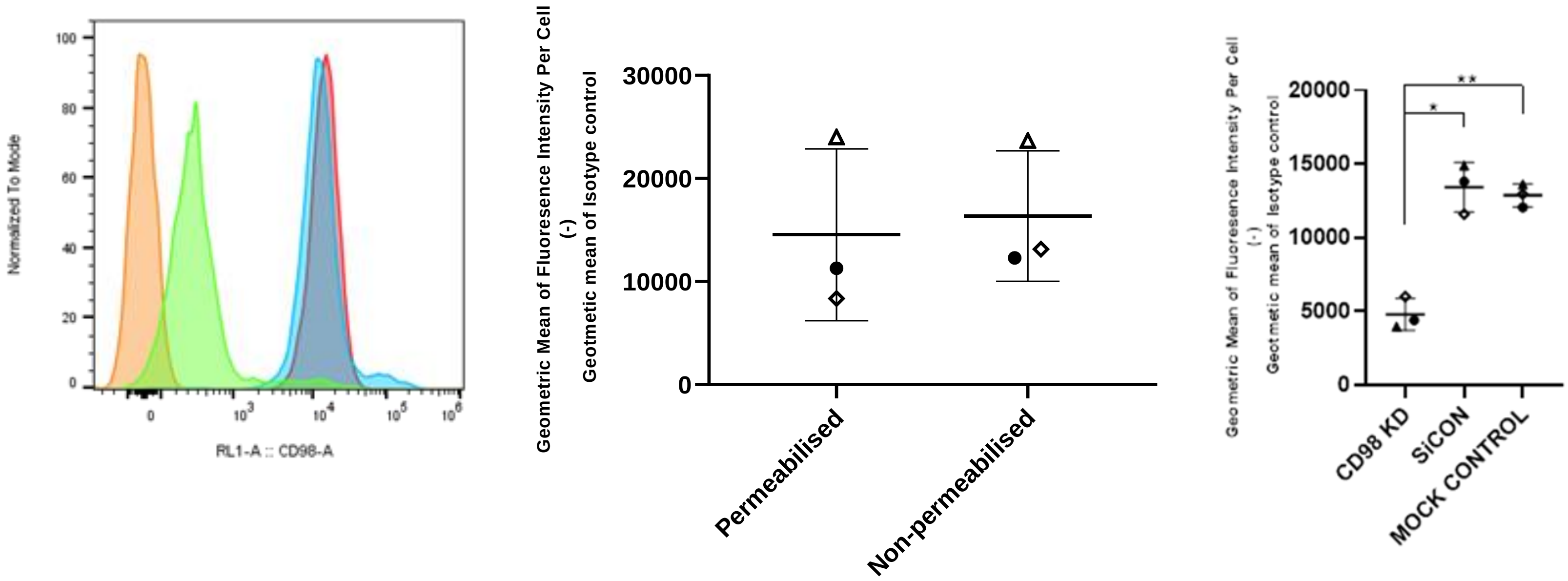

Normalized To Mode
RL1-A :: CD98-A
Geometric Mean of Fluoresence Intensity Per Cell
(-)
Geotmetic mean of Isotype control
30000
20000
10000
0
Permeabilised
Non-permeabilised
Geometric Mean of Fluoresence Intensity Per Cell
(-)
Geotmetic mean of Isotype control
20000
15000
10000
5000
0
CD98 KD
SiCON
MOCK CONTROL

### Supp. Fig. 2

Galectin-3 pulldown

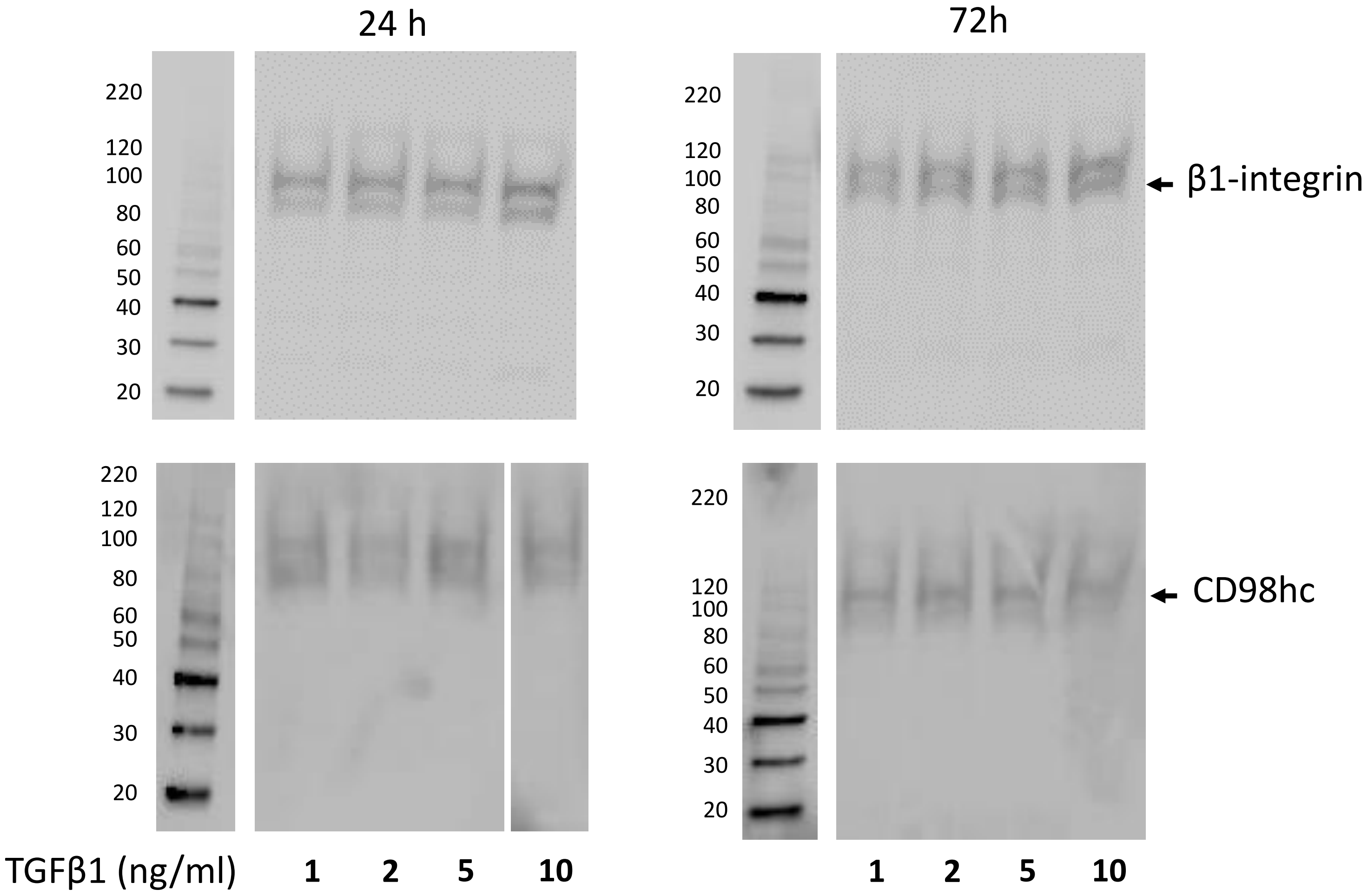

### Supp. Fig. 3A)

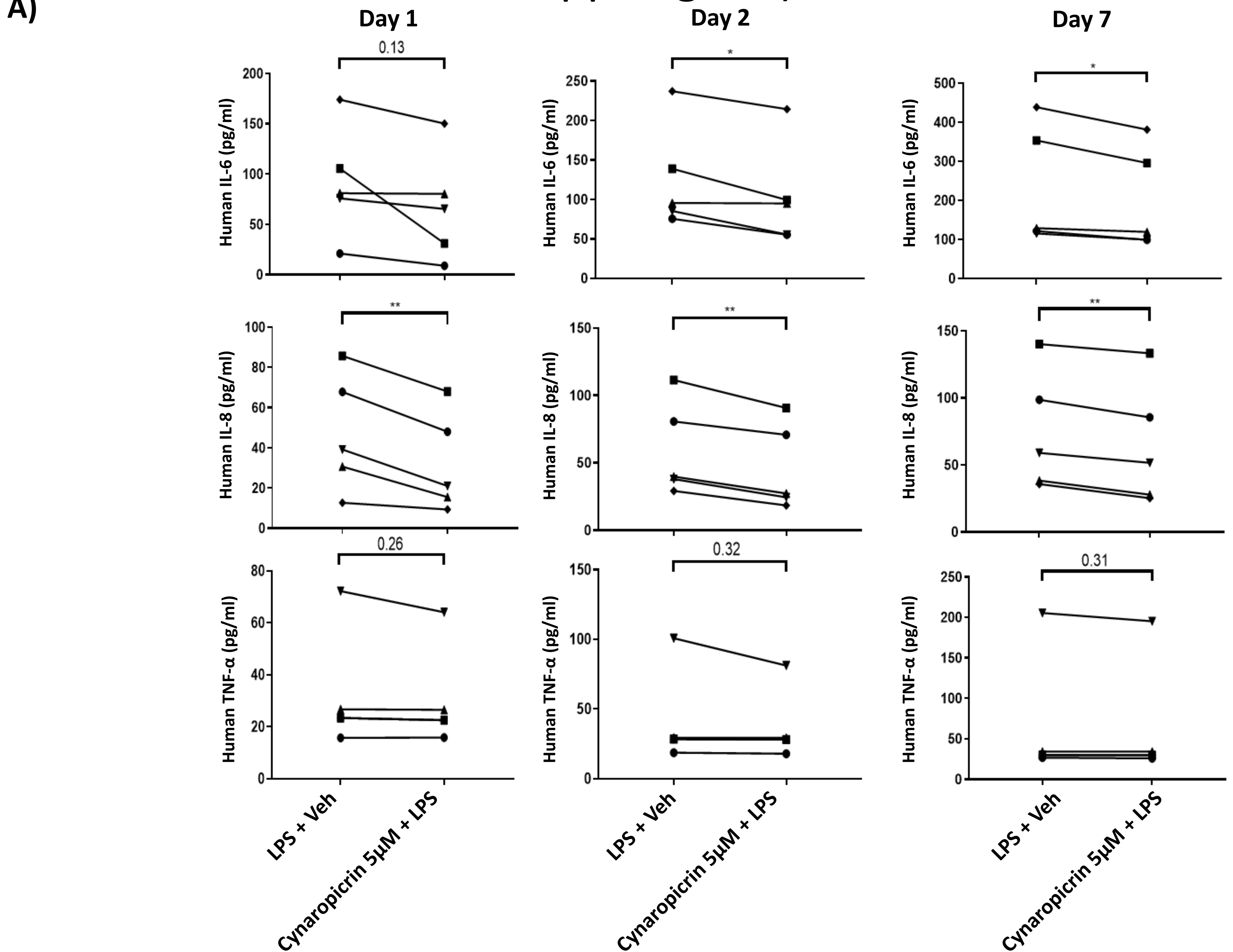

A)
Day 1
Day 2
Day 7
0.13
*
*
Human IL-6 (pg/ml)
**
**
**
Human IL-8 (pg/ml)
0.26
0.32
0.31
Human TNF-α (pg/ml)
LPS + Veh
Cynaropicrin 5µM + LPS

### Supp. Fig. 3B)

**B)**

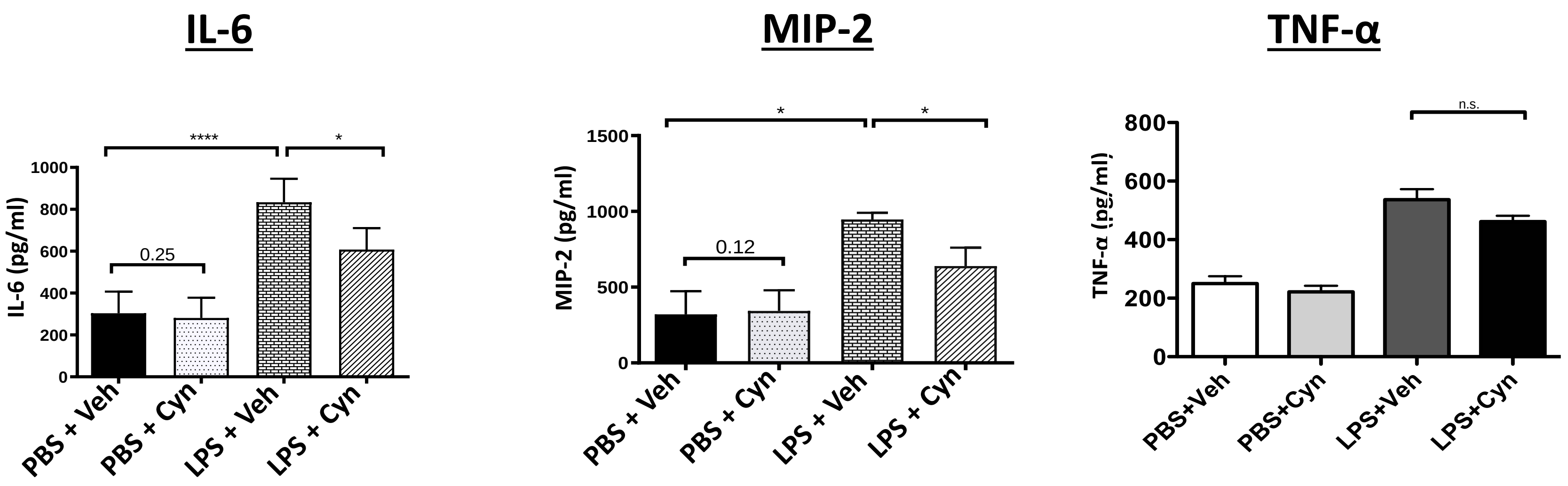
IL-6
MIP-2
TNF-α
IL-6 (pg/ml)
MIP-2 (pg/ml)
TNF-α (pg/ml)
****
*
0.25
0.12
n.s.
1000
800
600
400
200
0
1500
500
PBS + Veh
PBS + Cyn
LPS + Veh
LPS + Cyn
PBS+Veh
PBS+Cyn
LPS+Veh
LPS+Cyn

### Supp. Fig. 3C)

**C)**

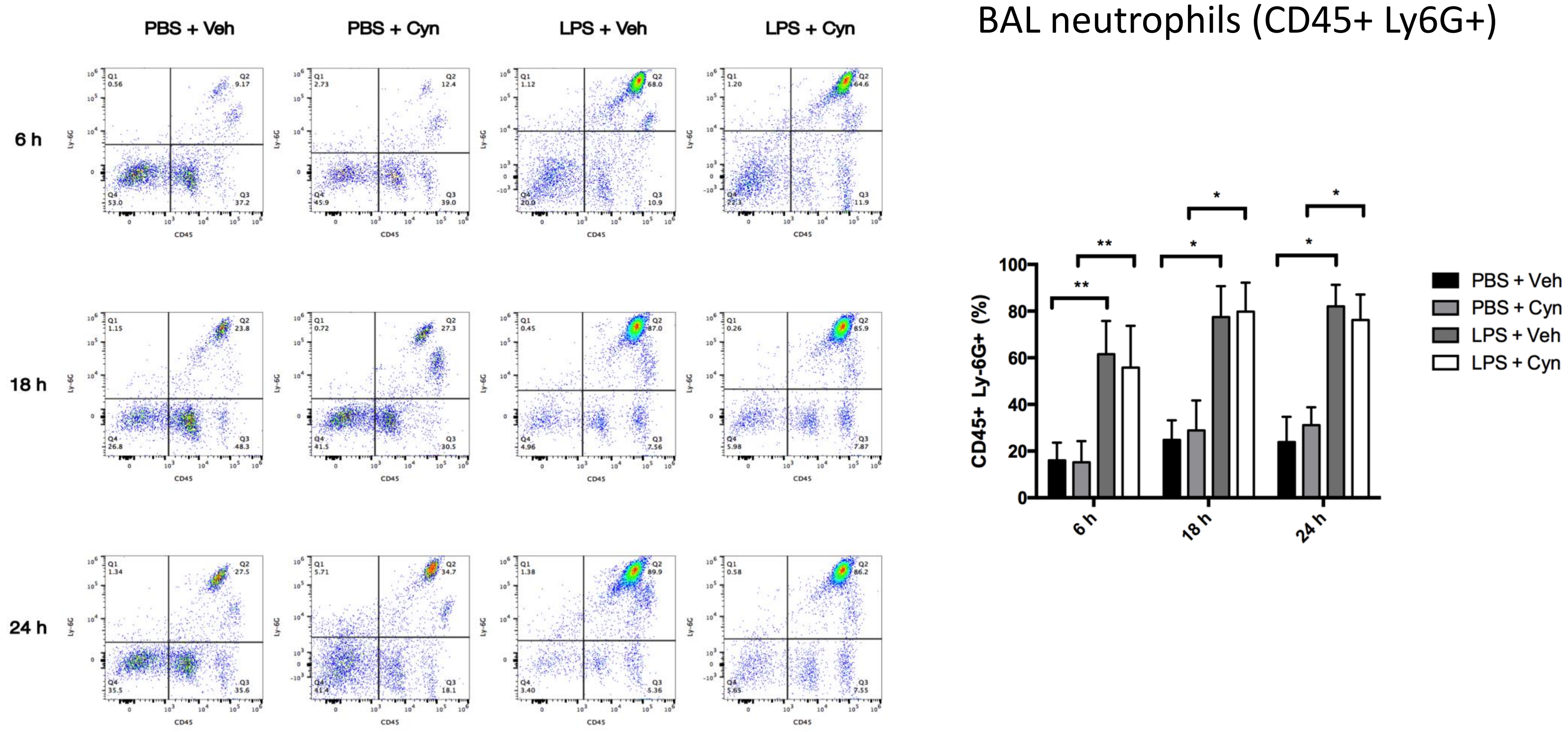

PBS + Veh
PBS + Cyn
LPS + Veh
LPS + Cyn
6 h
18 h
24 h
Ly-6G
CD45
BAL neutrophils (CD45+ Ly6G+)
CD45+ Ly-6G+ (%)
PBS + Veh
PBS + Cyn
LPS + Veh
LPS + Cyn
**
*

### Supp. Fig. 3D)

**D)**

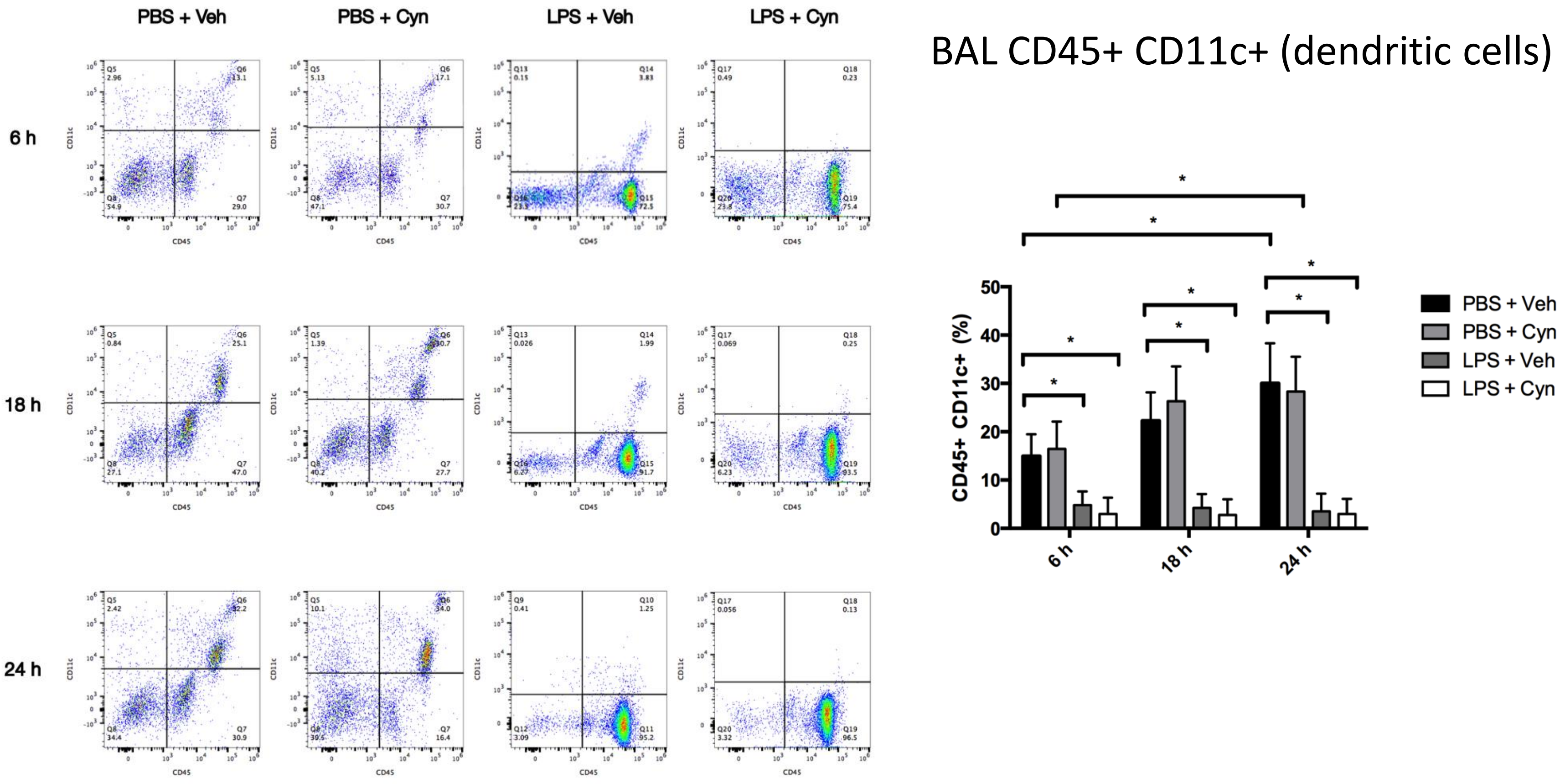

PBS + Veh
PBS + Cyn
LPS + Veh
LPS + Cyn
6 h
18 h
24 h
BAL CD45+ CD11c+ (dendritic cells)
CD45+ CD11c+ (%)
PBS + Veh
PBS + Cyn
LPS + Veh
LPS + Cyn

### Supp. Fig. 3E)

**E)**

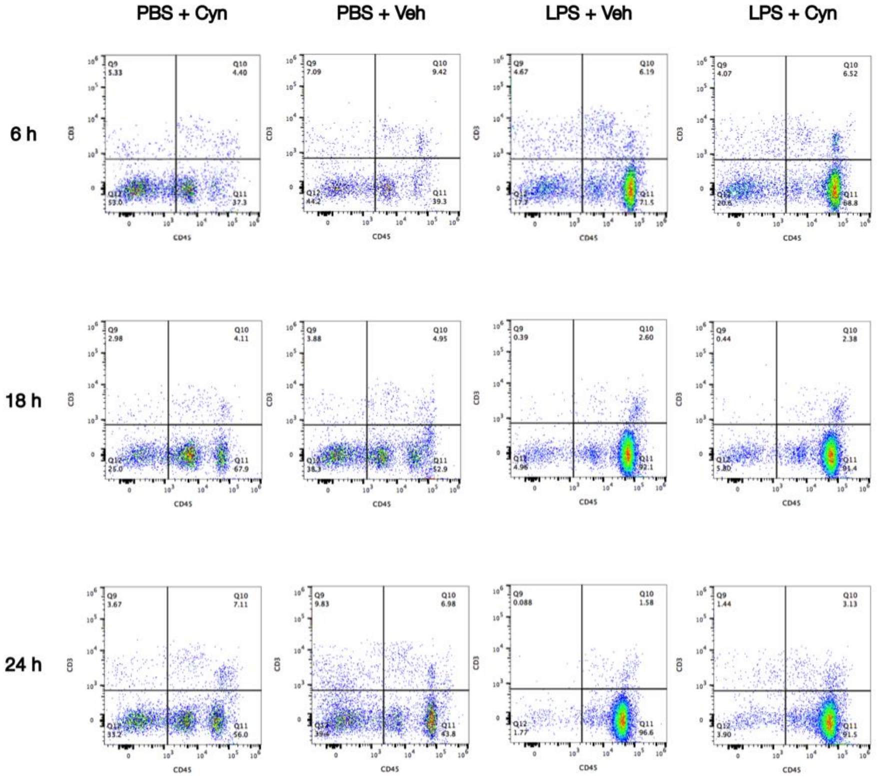


BAL CD45+ CD3+ (T cells)

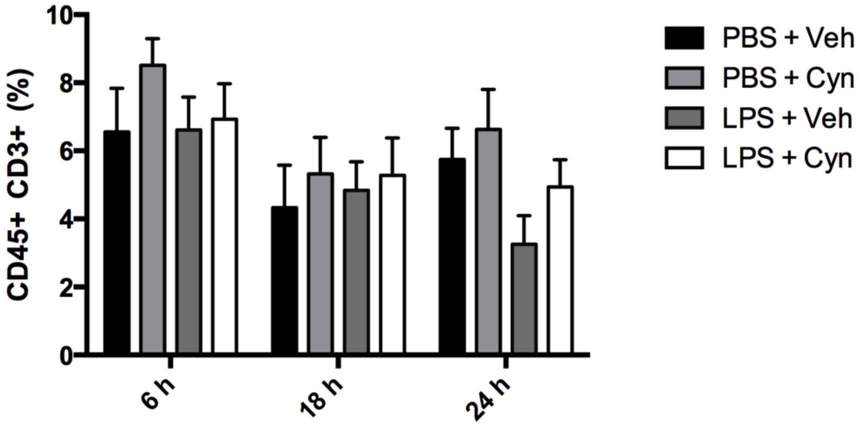

### Supp. Fig. 3F)

**F)**

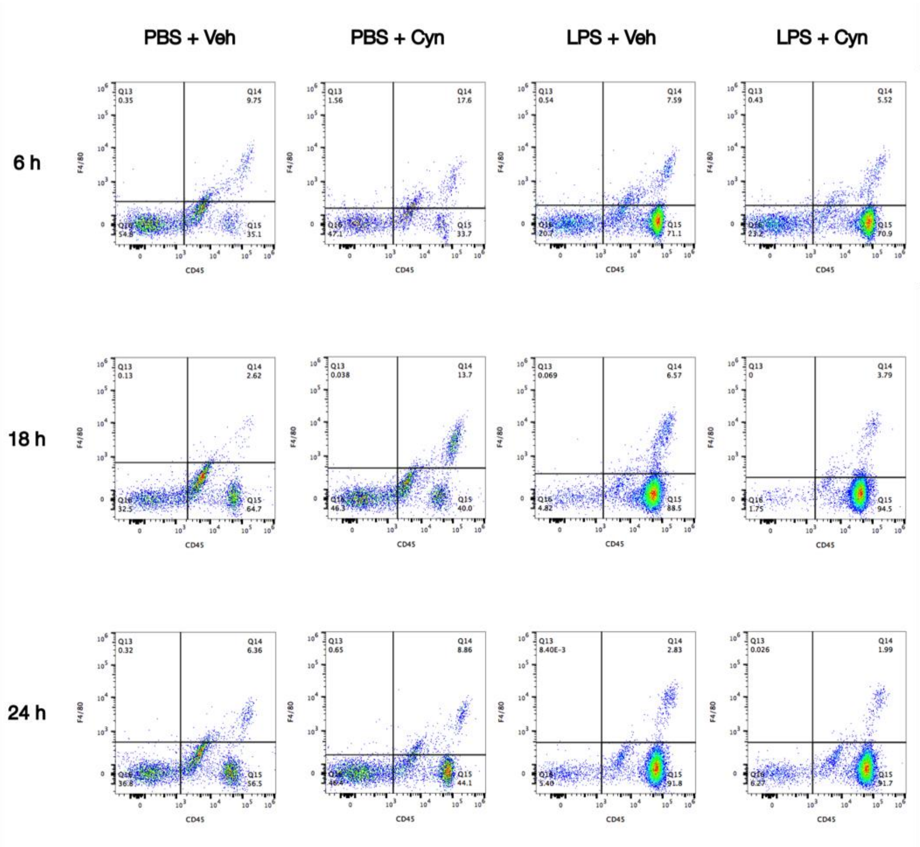


BAL CD45+ F4/80+ (macrophages)

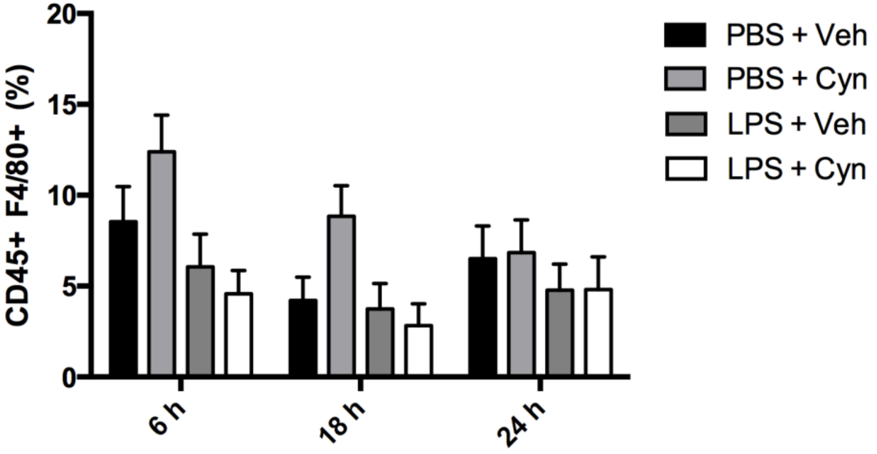

### Supp. Fig. 3G)

**G)**

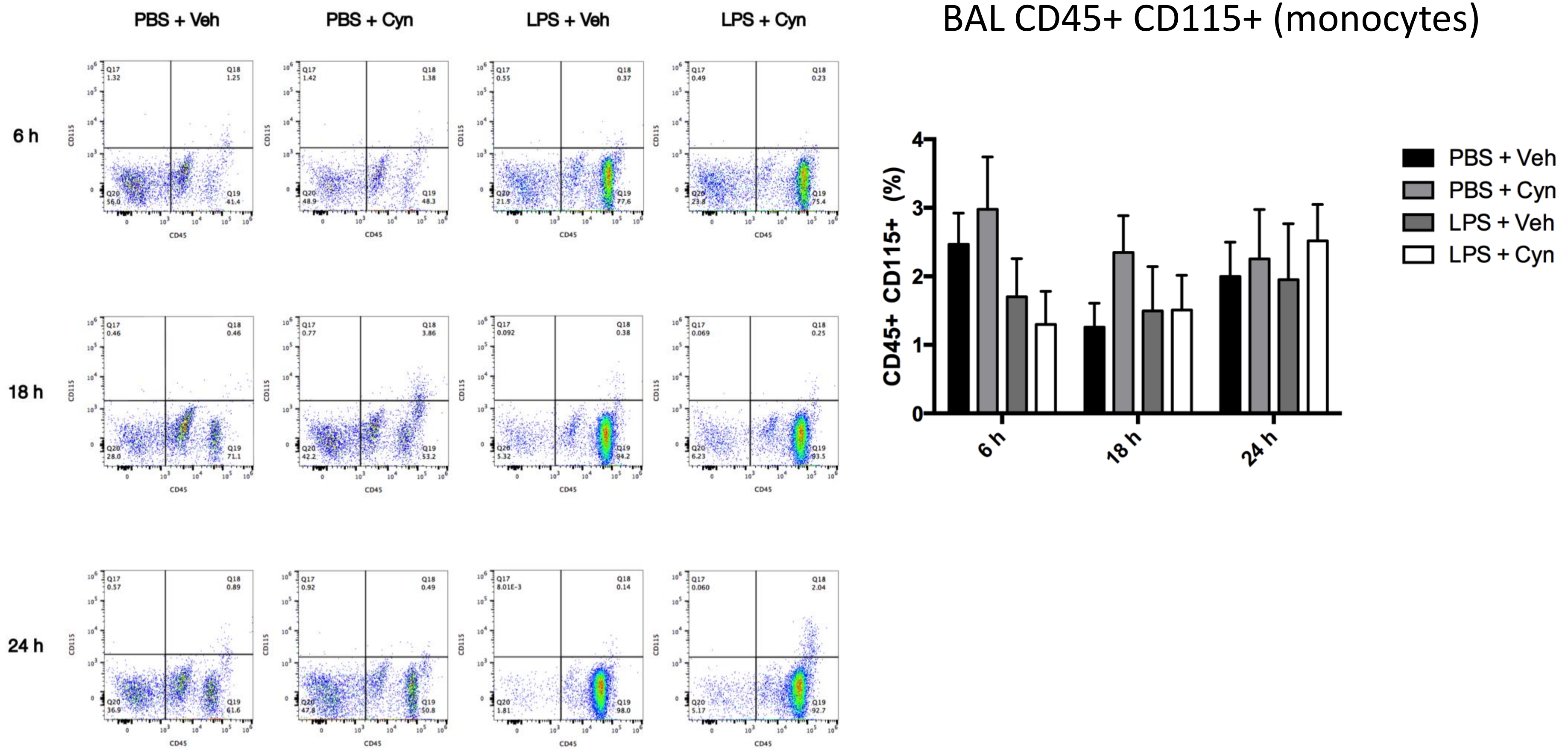

PBS + Veh
PBS + Cyn
LPS + Veh
LPS + Cyn
6 h
18 h
24 h
CD45
CD115
BAL CD45+ CD115+ (monocytes)
CD45+ CD115+ (%)
PBS + Veh
PBS + Cyn
LPS + Veh
LPS + Cyn

### Supp. Fig. 3H)

**H)**

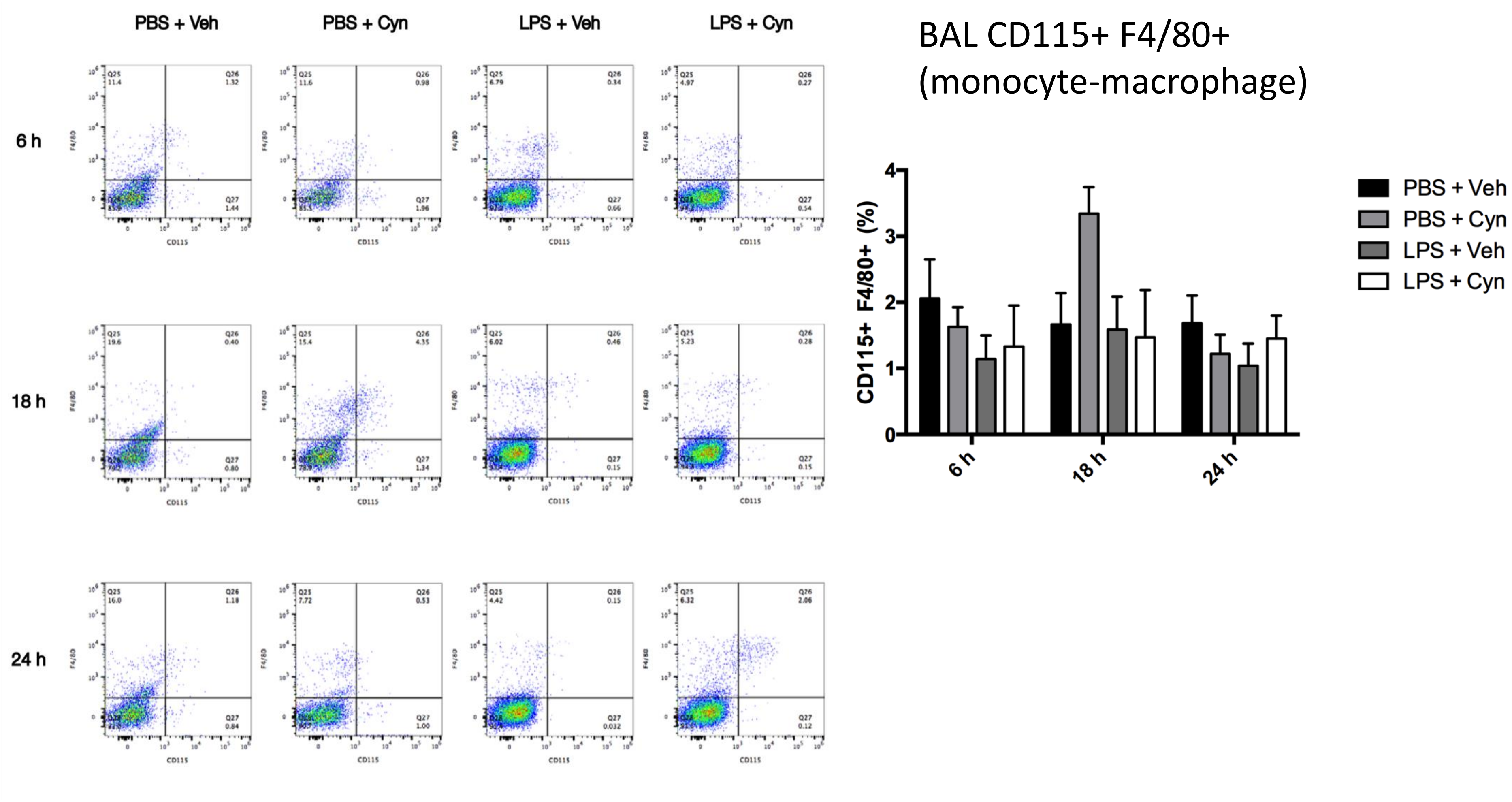
PBS + Veh
PBS + Cyn
LPS + Veh
LPS + Cyn
6 h
18 h
24 h
CD115
F4/80
BAL CD115+ F4/80+
(monocyte-macrophage)
CD115+ F4/80+ (%)
0
1
2
3
4
6 h
18 h
24 h
PBS + Veh
PBS + Cyn
LPS + Veh
LPS + Cyn

### Supp. Fig. 3I)

**I)**

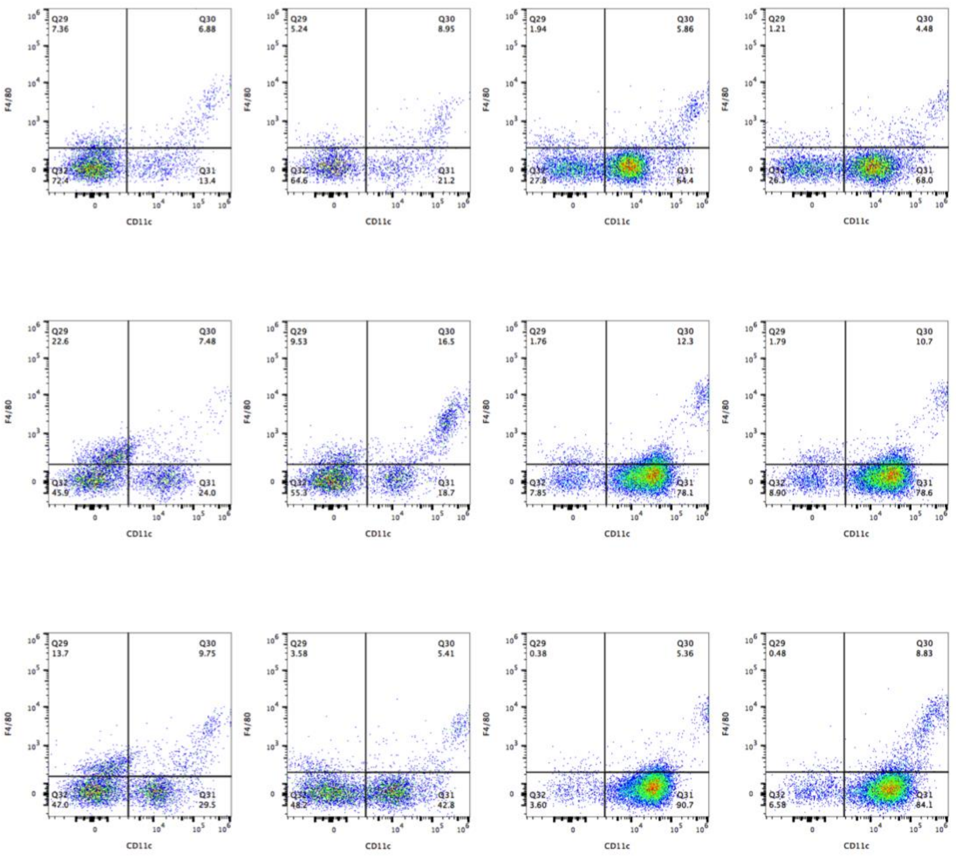

BAL CD11c+ F4/80+
(dendritic cell-macrophage)

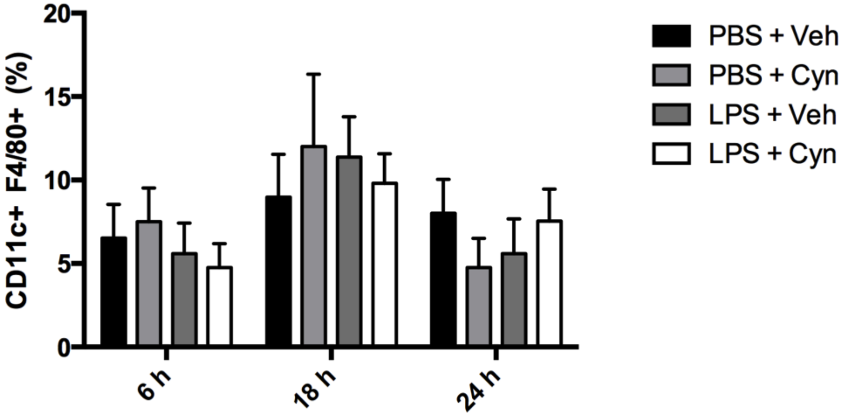

### Supp. Fig. 4A)

**A)**

**TT1, LPS**

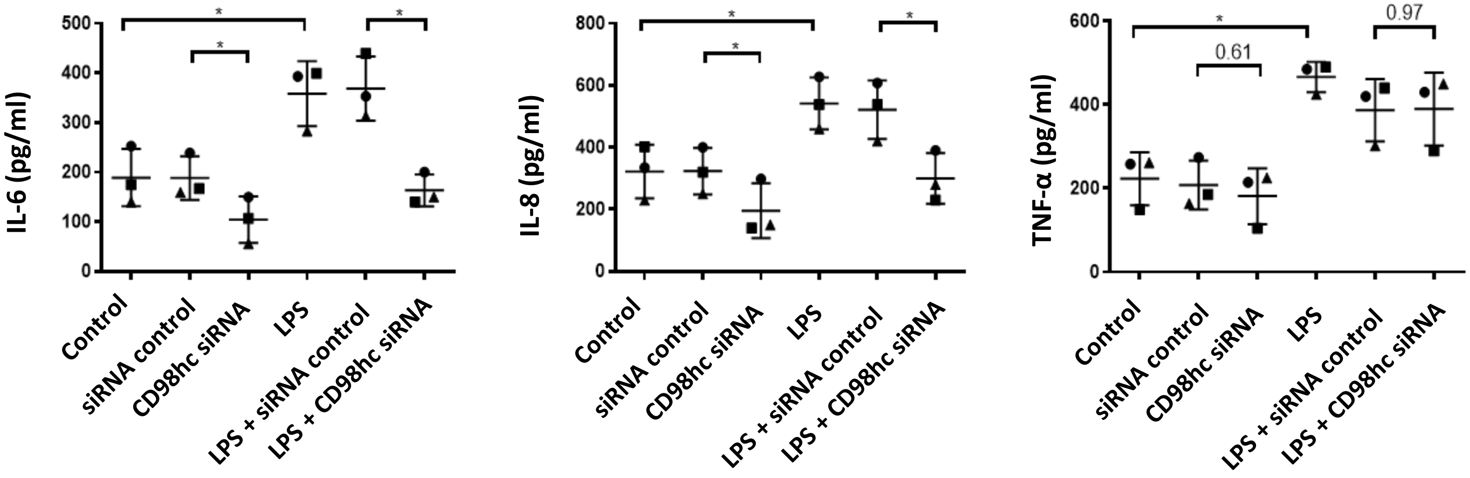

### Supp. Fig. 4B)

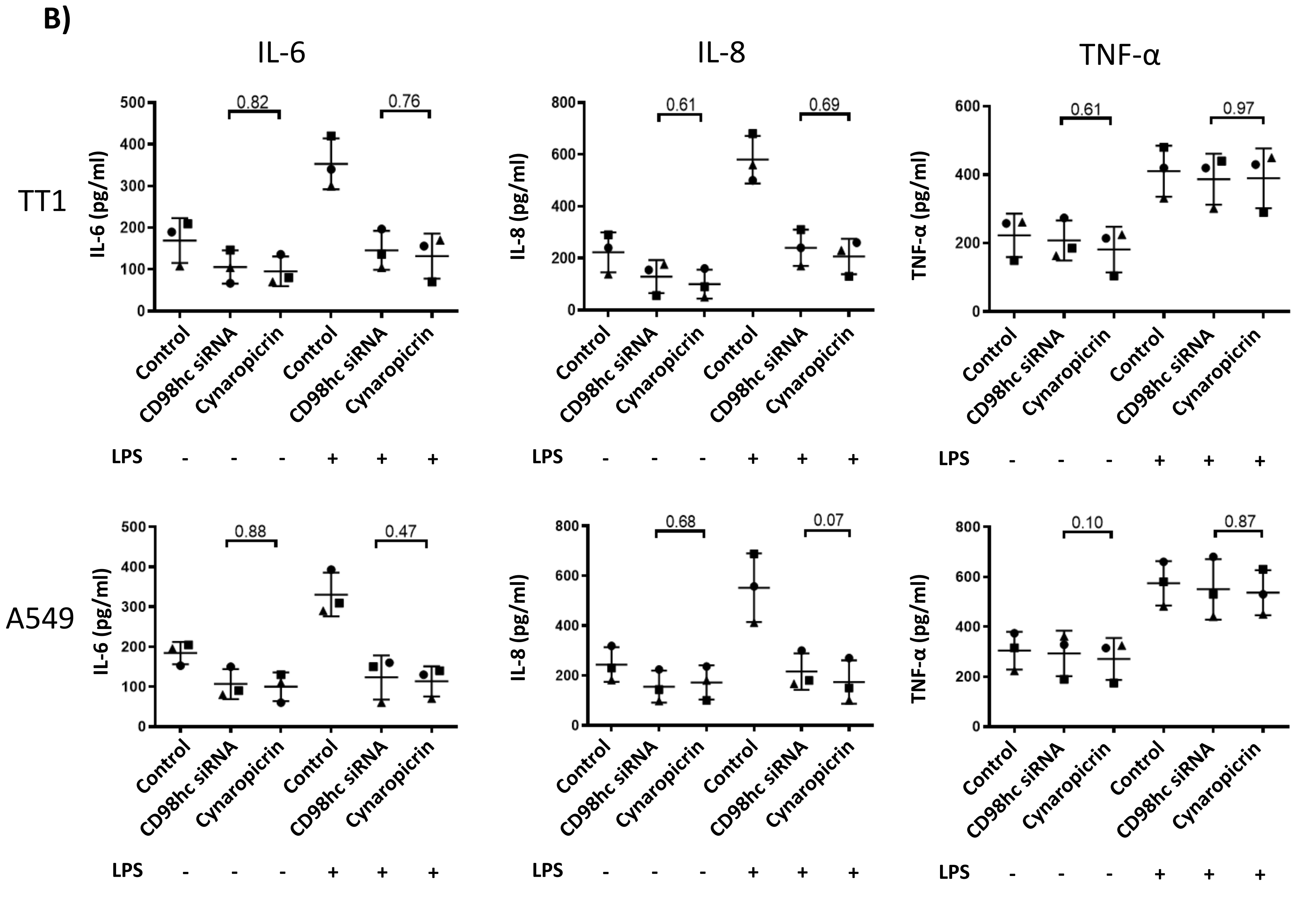
B)
IL-6
IL-8
TNF-α
TT1
A549
IL-6 (pg/ml)
IL-8 (pg/ml)
TNF-α (pg/ml)
0.82
0.76
0.61
0.69
0.61
0.97
0.88
0.47
0.68
0.07
0.10
0.87
Control
CD98hc siRNA
Cynaropicrin
LPS - - - + + +

### Supp. Fig. 4C)

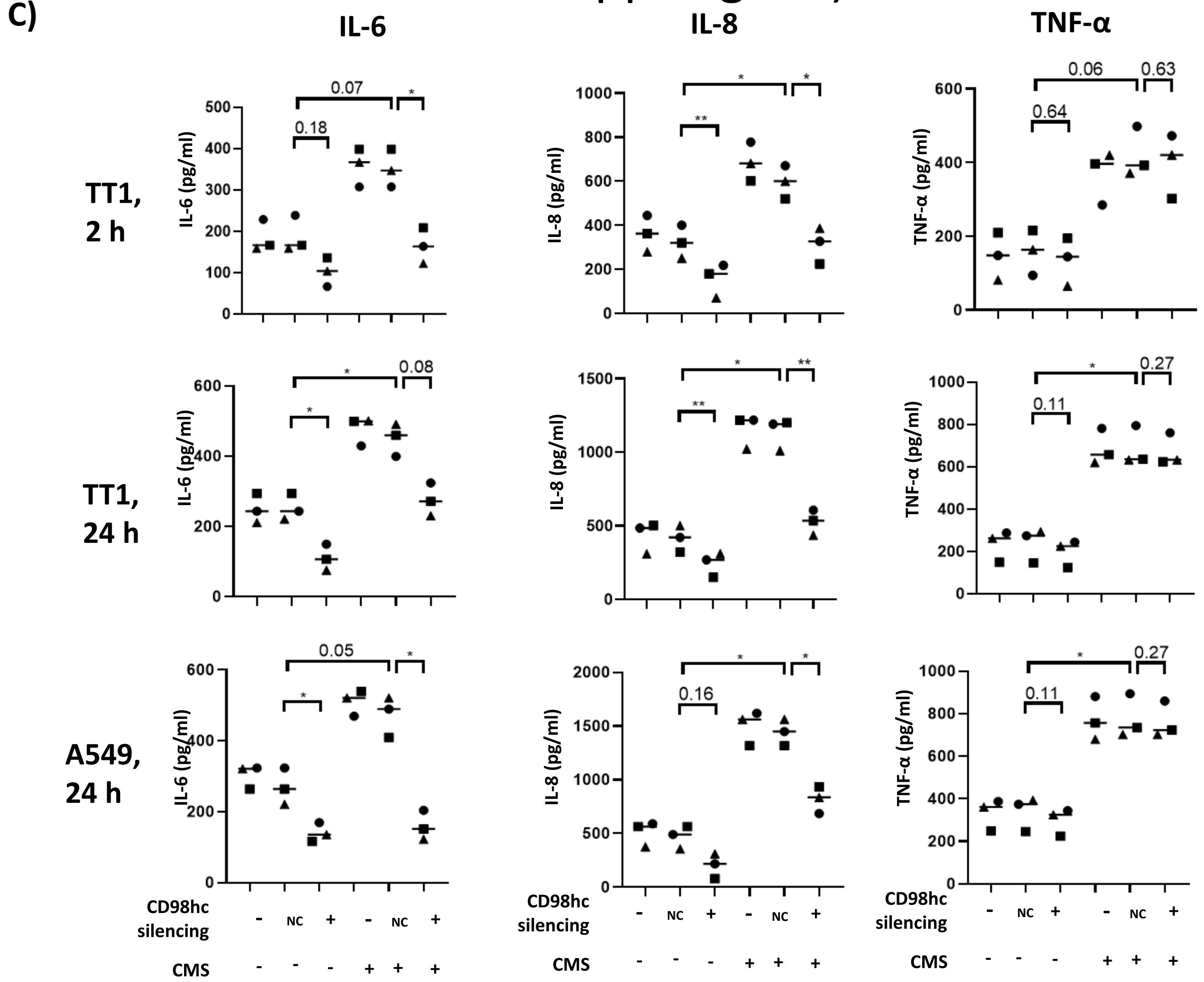
C)
IL-6
IL-8
TNF-α
TT1, 2 h
TT1, 24 h
A549, 24 h
IL-6 (pg/ml)
IL-8 (pg/ml)
TNF-α (pg/ml)
0.07
0.18
0.06
0.63
0.64
0.08
0.27
0.11
0.05
0.16
CD98hc silencing
- NC + - NC +
CMS
- - - + + +

### Supp. Fig. 4D)

**D)**

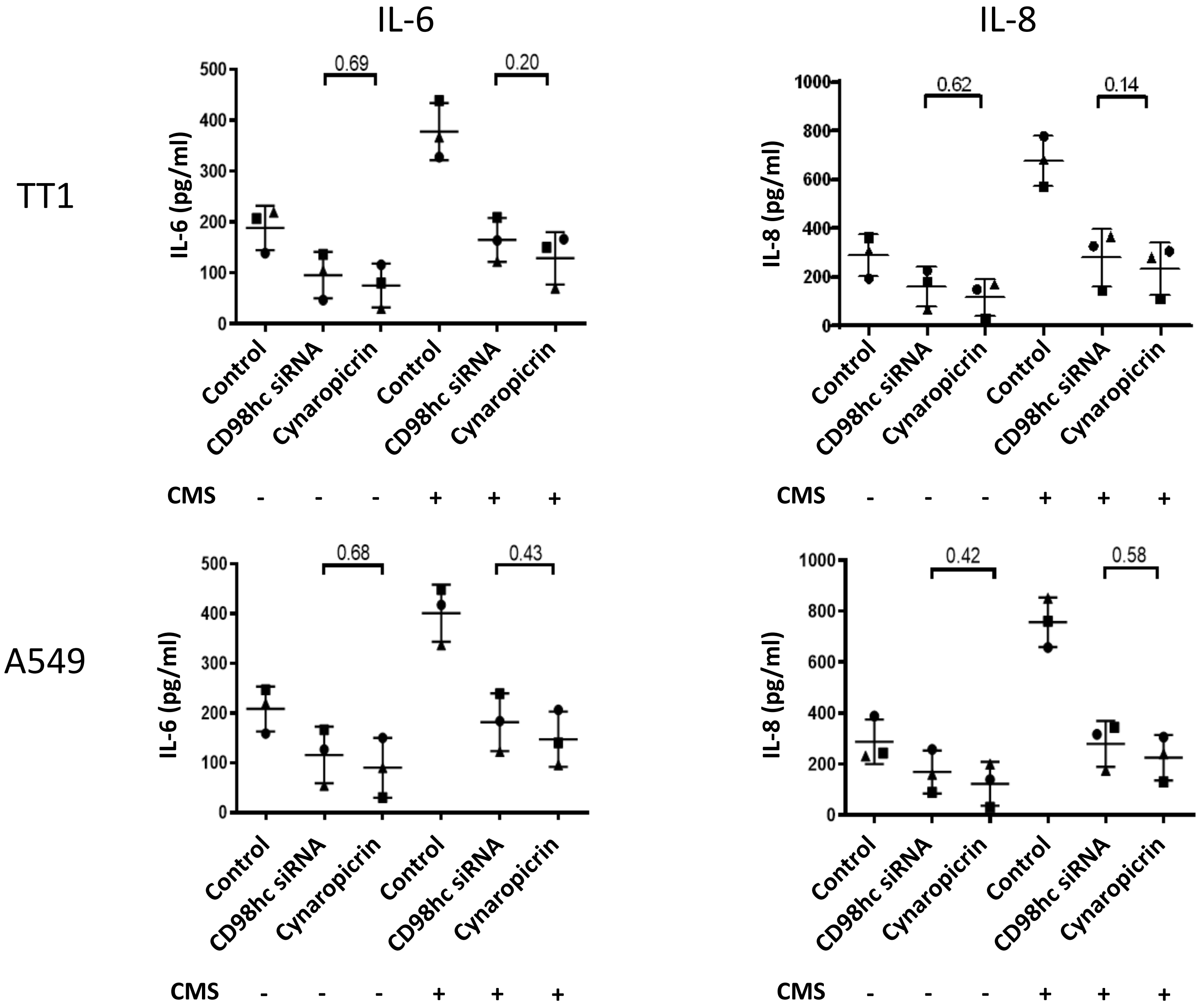
IL-6
IL-8
TT1
A549
IL-6 (pg/ml)
IL-8 (pg/ml)
0.69
0.20
0.62
0.14
0.68
0.43
0.42
0.58
0
100
200
300
400
500
0
200
400
600
800
1000
Control
CD98hc siRNA
Cynaropicrin
CMS - - - + + +

### Supp. Fig. 4E)

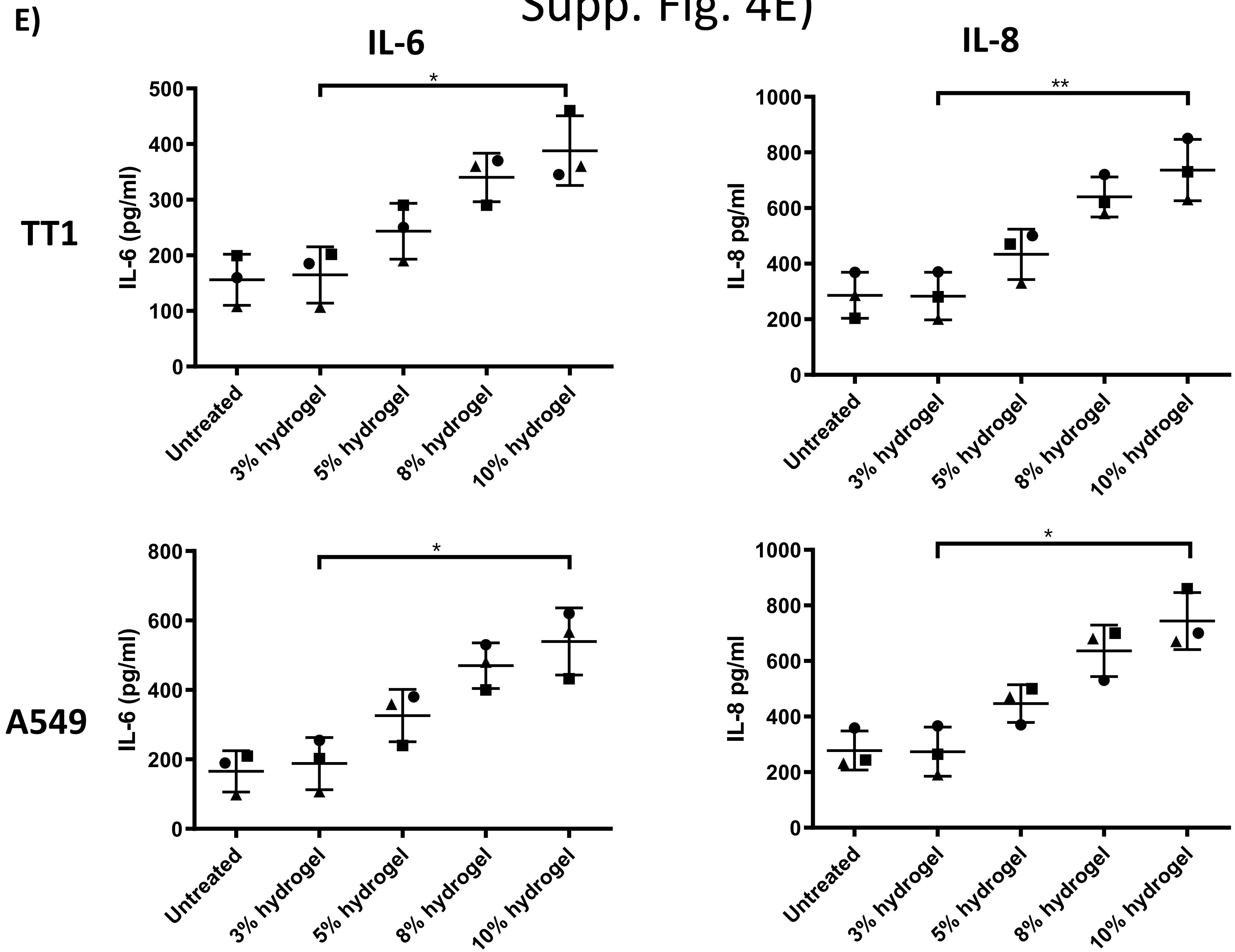
E)
IL-6
IL-8
TT1
A549
IL-6 (pg/ml)
IL-8 pg/ml
0
100
200
300
400
500
600
800
1000
*
**
Untreated
3% hydrogel
5% hydrogel
8% hydrogel
10% hydrogel

### Supp. Fig. 4F)

**F)**

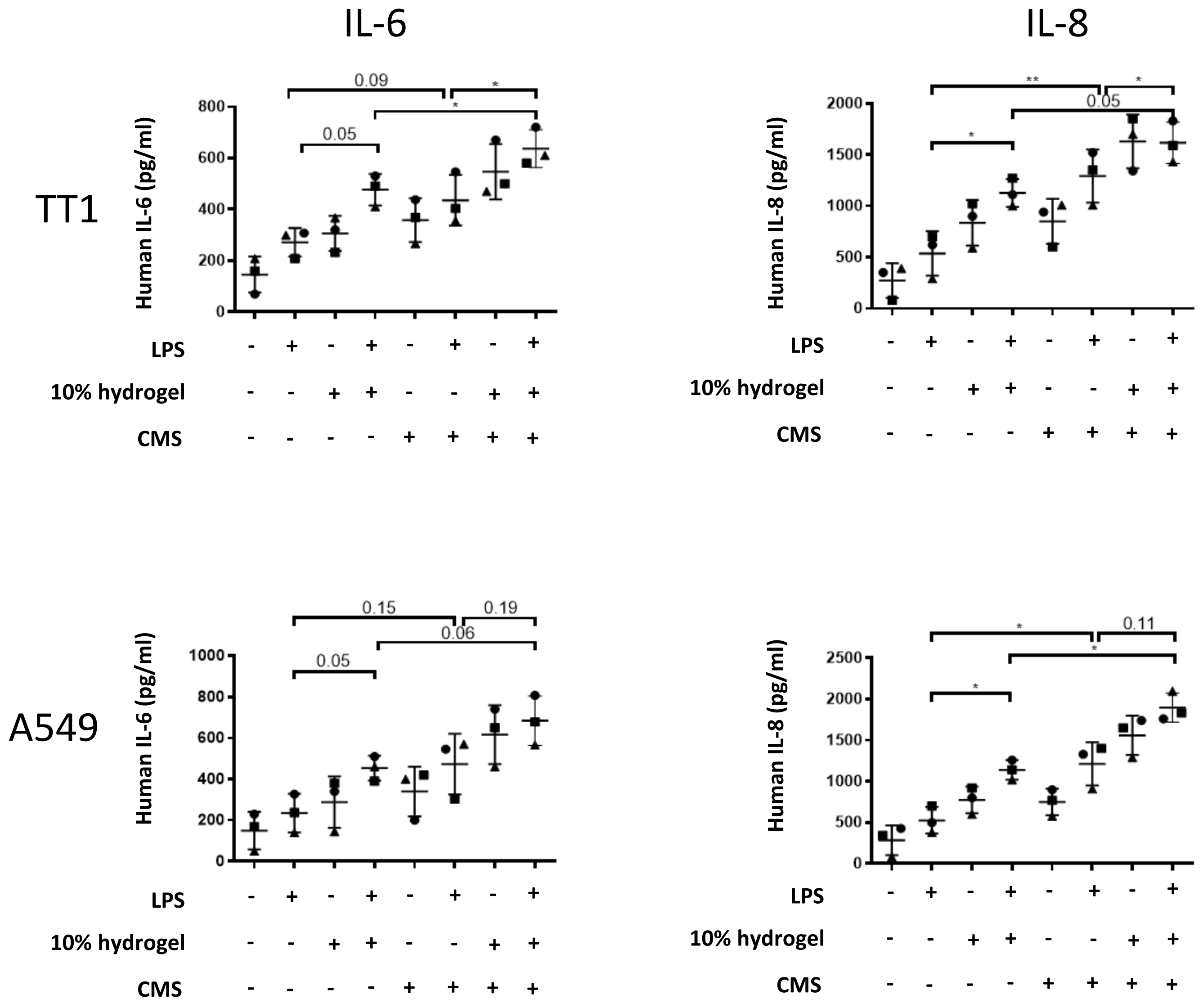
IL-6
IL-8
TT1
A549
Human IL-6 (pg/ml)
Human IL-8 (pg/ml)
0.09
0.05
0.15
0.19
0.06
0.05
0.11
LPS - + - + - + - +
10% hydrogel - - + + - - + +
CMS - - - - + + + +

### Supp. Fig. 5A)

**A)**

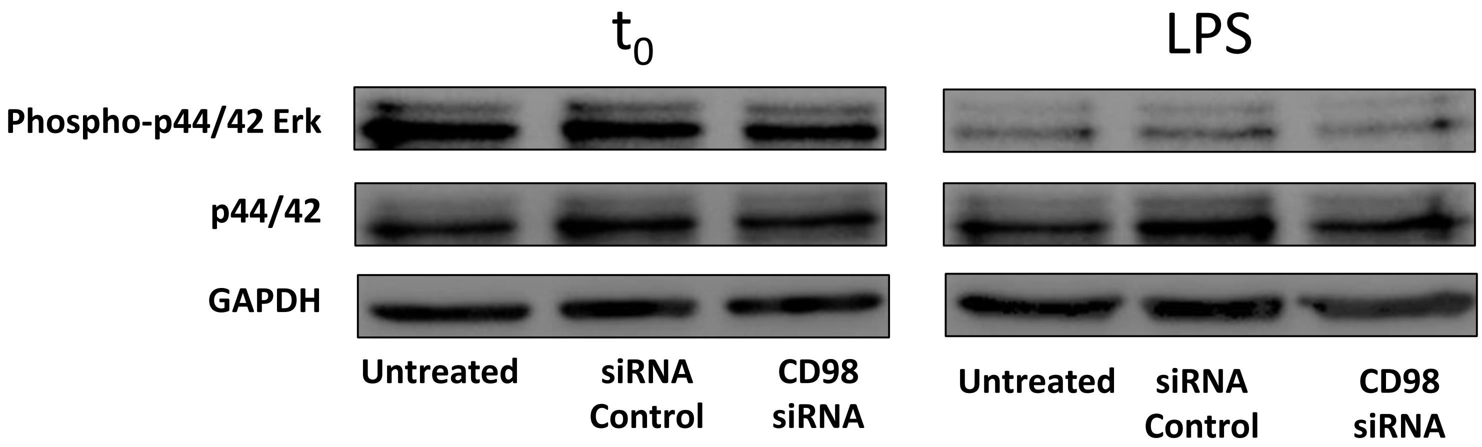

t0
LPS
Phospho-p44/42 Erk
p44/42
GAPDH
Untreated
siRNA Control
CD98 siRNA
Untreated
siRNA Control
CD98 siRNA

### Supp. Fig. 5B)

**B)**

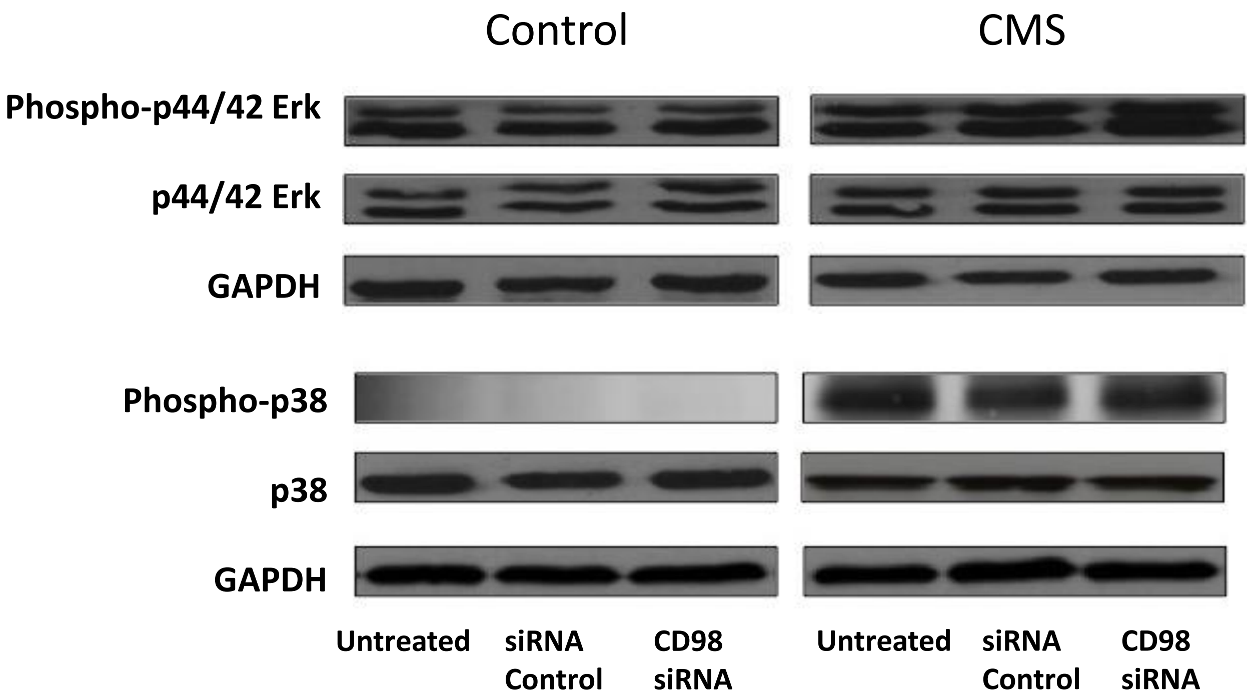
Control
CMS
Phospho-p44/42 Erk
p44/42 Erk
GAPDH
Phospho-p38
p38
GAPDH
Untreated
siRNA Control
CD98 siRNA
Untreated
siRNA Control
CD98 siRNA

### Supp. Fig. 6A)

**A)**

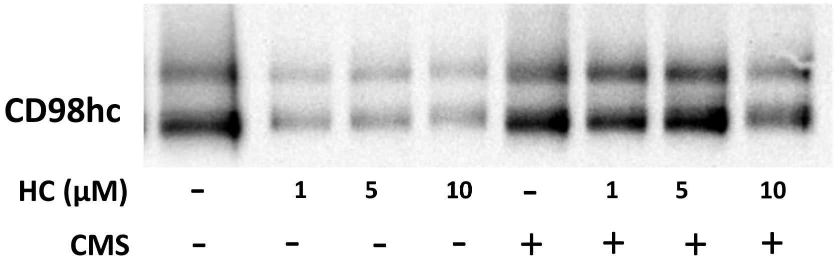


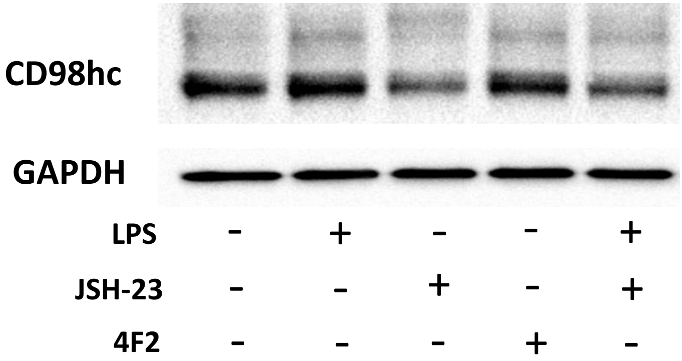

### Supp. Fig. 6B)

**B)**

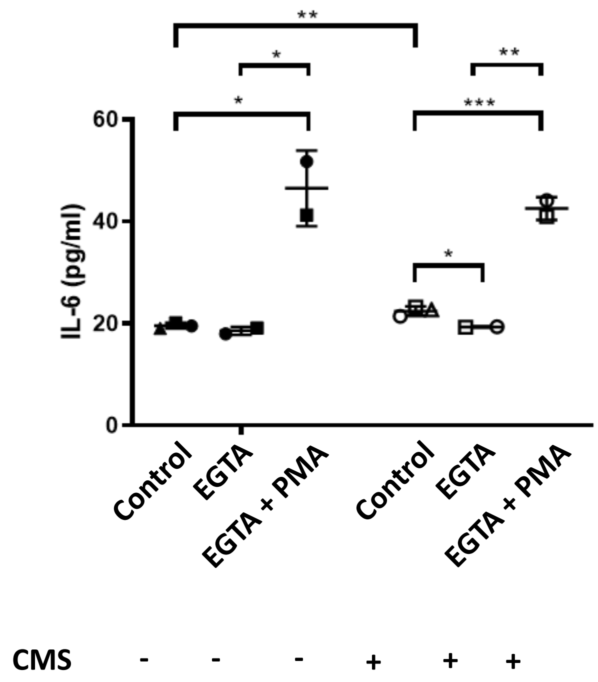

**
*
*
**
***
*
60
40
20
0
IL-6 (pg/ml)
Control
EGTA
EGTA + PMA
Control
EGTA
EGTA + PMA
CMS - - - + + +


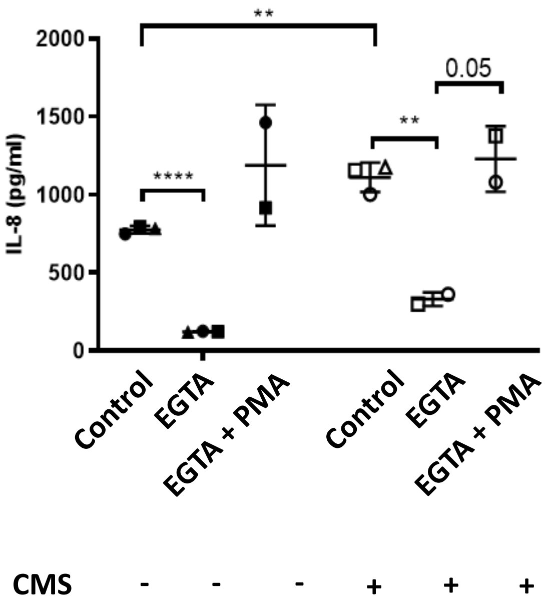

**
0.05
**
****
2000
1500
1000
500
0
IL-8 (pg/ml)
Control
EGTA
EGTA + PMA
Control
EGTA
EGTA + PMA
CMS - - - + + +

### Supp. Fig. 7

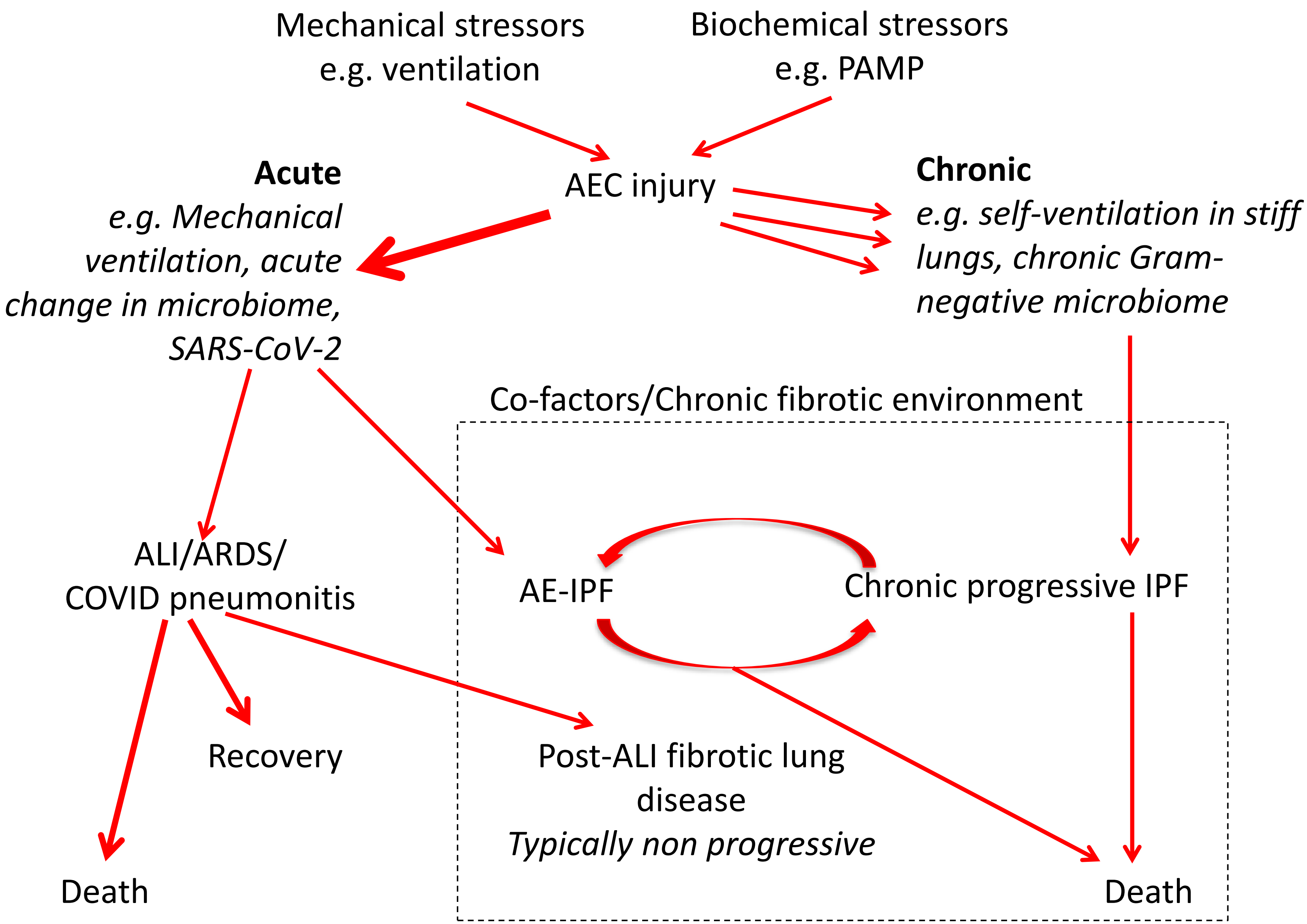
Mechanical stressors
e.g. ventilation
Biochemical stressors
e.g. PAMP
AEC injury
Acute
e.g. Mechanical ventilation, acute change in microbiome, SARS-CoV-2
Chronic
e.g. self-ventilation in stiff lungs, chronic Gram-negative microbiome
Co-factors/Chronic fibrotic environment
ALI/ARDS/
COVID pneumonitis
AE-IPF
Chronic progressive IPF
Recovery
Post-ALI fibrotic lung disease
Typically non progressive
Death
Death

### Table 1

| Method | Primary Antibody | Isotype control | Secondary Antibody |
|---|---|---|---|
| Immunofluorescence | Galectin-3 (Santa-Cruz, sc-23938) | Purified Rat IgG2a (BD Pharmingen, 559073) | Goat anti-Rat IgG (H+L) Alexa Fluor 488 (ThermoFisher Scientific, A-11006)<br>Goat anti-Rat IgG (H+L) Alexa Fluor 594 (ThermoFisher Scientific, A-11007) |
| | CD98 (Santa-Cruz, sc-376815) | Negative Control Mouse IgG1 (Dako, X0931) | Goat anti-Mouse IgG (H+L) Alexa Fluor 488 (ThermoFisher Scientific, A-11001)<br>Goat anti-Mouse IgG (H+L) Alexa Fluor 594 (ThermoFisher Scientific, A-11005) |
| | β1-integrin (Abcam, ab24693) | Negative Control Mouse IgG1 (Dako, X0931) | Goat anti-Mouse IgG (H+L) Alexa Fluor 488 (ThermoFisher Scientific, A-11001)<br>Goat anti-Mouse IgG (H+L) Alexa Fluor 594 (ThermoFisher Scientific, A-11005) |
| Western blotting/Co-IP | Galectin-3 (Santa-Cruz, sc-23938) | Purified Rat IgG2a (BD Pharmingen, 559073) | Goat anti-Rat IgG HRP (ThermoFisher Scientific, 31470) |
| | CD98 (Santa-Cruz, sc-376815) | Negative Control Mouse IgG1 (Dako, X0931) | Polyclonal goat anti-mouse HRP (Dako, P0447) |
| | β1-integrin (BD Pharmingen, 610467) | Negative Control Mouse IgG1 (Dako, X0931) | Polyclonal goat anti-mouse HRP (Dako, P0447) |
| | ATP1A1 (Novus Biologicals, NB300-146) | Negative Control Mouse IgG1 (Dako, X0931) | Polyclonal goat anti-mouse HRP (Dako, P0447) |
| | SARS-CoV-2 Spike Antibody (Sino Biological, 40150-R007) | N/A | Polyclonal goat anti-rabbit HRP (Dako, P0448) |
| PLA | Galectin-3 (ThermoFisher Scientific, PA5-34819) | Purified Rabbit IgG (BD Pharmingen, 550875) | N/A |
| | CD98 (Santa-Cruz, sc-376815) | Negative Control Mouse IgG1 (Dako, X0931) | N/A |
| | β1-integrin (Abcam, ab24693) | Negative Control Mouse IgG1 (Dako, X0931) | N/A |
| Immunohistochemistry | Galectin-3 (Santa-Cruz, sc-23938) | Purified Rat IgG2a (BD Pharmingen, 559073) | Goat anti-Rat IgG HRP (ThermoFisher Scientific, 31470) |
| | CD98 (Santa-Cruz, sc-376815) | Negative Control Mouse IgG1 (Dako, X0931) | EnVision Flex HRP (Dako) |
| | β1-integrin (Abcam, ab24693) | Negative Control Mouse IgG1 (Dako, X0931) | EnVision Flex HRP (Dako) |
| | E-cadherin (Dako, M3612) | Negative Control Mouse IgG1 (Dako, X0931) | EnVision Flex HRP (Dako) |

### Table 2

#### Table of p-values for all comparisons shown in Results/Supplemental materials (paired t-tests expect where otherwise indicated)

#### Figure 1A:

| | | **P value** |
|---|---|---|
| % Galectin-3 immunostaining | NFC vs IPF | 0.0286* |
| % CD98 immunostaining | NFC vs IPF | 0.0097* |
| % β1-integrin immunostaining | NFC vs IPF | 0.0439* |

#### Figure 1B:

| **SLC3A2** | **P value** |
|---|---|
| Non-stimulated vs TGF-β1 | 0.0195 |
| **LGALS3** | |
| Non-stimulated vs TGF-β1 | 0.0156 |

* Unpaired t-test
† Two-way ANOVA

#### Figure 2A:

| | | **P value** |
|---|---|---|
| CD98 Band density | Non stim'd vs 24h TGF-β1 | 0.0260 |
| | Non stim'd vs 24h TGF-β1 | 0.0307 |

#### Figure 2B:

| | | **P value** |
|---|---|---|
| Galectin-3/β1-integrin | Control tissue vs TGF-β1 tissue | 0.0313* |
| Galectin-3/CD98 | Control tissue vs TGF-β1 tissue | 0.0313* |

* Unpaired t-test
† Two-way ANOVA

### Figure 3A:

| | | **P value** |
|---|---|---|
| Human IL-6 (pg/ml) | Untreated vs Cynaropicrin 5μM + PBS | 0.0035 |
| | Untreated vs LPS + Veh | 0.0595 |
| | PBS + Veh vs Cynaropicrin 5μM + PBS | 0.0006 |
| | PBS + Veh vs LPS + Veh | 0.0480 |
| | LPS + Veh vs Cynaropicrin 5μM + LPS | 0.0248 |
| Human IL-8 (pg/ml) | Untreated vs Cynaropicrin 5μM + PBS | 0.0448 |
| | Untreated vs LPS + Veh | 0.0289 |
| | PBS + Veh vs Cynaropicrin 5μM + PBS | 0.0608 |
| | PBS + Veh vs LPS + Veh | 0.0195 |
| | LPS + Veh vs Cynaropicrin 5μM + LPS | 0.0022 |
| Human TNF-α (pg/ml) | Untreated vs Cynaropicrin 5μM + PBS | 0.0148 |
| | Untreated vs LPS + Veh | 0.0821 |
| | PBS + Veh vs Cynaropicrin 5μM + PBS | 0.0160 |
| | PBS + Veh vs LPS + Veh | 0.0463 |
| | LPS + Veh vs Cynaropicrin 5μM + LPS | 0.3294 |

* Unpaired t-test
† Two-way ANOVA

### Figure 3B:

| | | **P value** |
|---|---|---|
| Mouse IL-8 | +18h: PBS + Veh vs LPS + Veh | 0.0125† |
| | +18h: PBS + Cyn vs LPS + Cyn | 0.0029† |
| | +24h: PBS + Veh vs LPS + Veh | <0.0001† |
| | +24h: LPS + Cyn vs LPS + Cyn | <0.0001† |

### Figure 4A:

| **A549** | | **P value** |
|---|---|---|
| IL-6 (pg/ml) | Untreated vs LPS | 0.1504 |
| | siRNA control vs CD98hc siRNA | 0.0296 |
| | LPS + siRNA control vs LPS + CD98hc siRNA | 0.0017 |
| IL-8 (pg/ml) | Untreated vs LPS | 0.0106 |
| | siRNA control vs CD98hc siRNA | 0.2360 |
| | LPS + siRNA control vs LPS + CD98hc siRNA | 0.0440 |
| TNF-α (pg/ml) | Untreated vs LPS | 0.0010 |
| | siRNA control vs CD98hc siRNA | 0.1015 |
| | LPS + siRNA control vs LPS + CD98hc siRNA | 0.8721 |

* Unpaired t-test
† Two-way ANOVA

### Figure 4B:

| **A549** | | **P value** |
|---|---|---|
| IL-6 (pg/ml) | siRNA control vs CD98hc siRNA | 0.15 |
| | siRNA control vs siRNA control + CMS | 0.0358 |
| | siRNA control + CMS vs CD98hc siRNA + CMS | 0.0489 |
| IL-8 (pg/ml) | siRNA control vs CD98hc siRNA | 0.19 |
| | siRNA control vs siRNA control + CMS | 0.0176 |
| | siRNA control + CMS vs CD98hc siRNA + CMS | 0.0324 |
| TNF-α (pg/ml) | siRNA control vs CD98hc siRNA | 0.3257 |
| | siRNA control vs siRNA control + CMS | 0.0371 |
| | siRNA control + CMS vs CD98hc siRNA + CMS | 0.7030 |

* Unpaired t-test
† Two-way ANOVA

### Figure 4C:

| A549 | | P value |
|---|---|---|
| IL-6 (pg/ml) | Untreated vs 3% hydrogel | 0.8080 |
| | Untreated vs 10% hydrogel | 0.0099 |
| | 10% hydrogel vs 10% hydrogel + CD98hc siRNA | 0.0866 |
| | Untreated vs CMS | 0.0001 |
| | CMS vs 3% hydrogel + CMS | 0.7445 |
| | CMS vs 10% hydrogel + CMS | 0.0132 |
| | 10% hydrogel + CMS vs 10% hydrogel + CMS + CD98hc siRNA | 0.0039 |
| IL-8 (pg/ml) | Untreated vs 3% hydrogel | 0.9504 |
| | Untreated vs 10% hydrogel | 0.0267 |
| | 10% hydrogel vs 10% hydrogel + CD98hc siRNA | 0.0729 |
| | Untreated vs CMS | 0.0002 |
| | CMS vs 3% hydrogel + CMS | 0.7462 |
| | CMS vs 10% hydrogel + CMS | 0.0131 |
| | 10% hydrogel + CMS vs 10% hydrogel + CMS + CD98hc siRNA | 0.0271 |

* Unpaired t-test
† Two-way ANOVA

### Figure 5C:

| TT1 | | P value |
|---|---|---|
| IL-6 (pg/ml) | Untreated vs JSH-23 | 0.4692 |
| | Untreated vs LPS | 0.0285 |
| | LPS vs LPS + JSH-23 | 0.1569 |
| IL-8 (pg/ml) | Untreated vs JSH-23 | 0.0427 |
| | Untreated vs LPS | 0.0121 |
| | LPS vs LPS + JSH-23 | 0.1874 |
| **A549** | | **P value** |
| IL-6 (pg/ml) | Untreated vs JSH-23 | 0.2417 |
| | Untreated vs LPS | 0.0021 |
| | LPS vs LPS + JSH-23 | 0.0003 |
| IL-8 (pg/ml) | Untreated vs JSH-23 | 0.4796 |
| | Untreated vs LPS | 0.0009 |
| | LPS vs LPS + JSH-23 | 0.0015 |

* Unpaired t-test
† Two-way ANOVA

### Figure 5D:

| **TT1** | | **P value** |
|---|---|---|
| IL-6 (pg/ml) | Control vs CD98hc siRNA | 0.0053 |
| | Control vs JSH-23 | 0.1702 |
| | Control vs Control + CMS | 0.1058 |
| | CD98hc siRNA vs JSH-23 | 0.8169 |
| | Control + CMS vs CD98hc siRNA + CMS | 0.0018 |
| | Control + CMS vs JSH-23 + CMS | 0.0002 |
| | CD98hc siRNA + CMS vs JSH-23 + CMS | 0.0021 |
| IL-8 (pg/ml) | Control vs CD98hc siRNA | 0.1102 |
| | Control vs JSH-23 | 0.0040 |
| | Control vs Control + CMS | 0.0344 |
| | CD98hc siRNA vs JSH-23 | 0.7949 |
| | Control + CMS vs CD98hc siRNA + CMS | 0.0178 |
| | Control + CMS vs JSH-23 + CMS | 0.0188 |
| | CD98hc siRNA + CMS vs JSH-23 + CMS | 0.1484 |

* Unpaired t-test
† Two-way ANOVA

### Figure 5D:

| **A549** | | **P value** |
|---|---|---|
| IL-6 (pg/ml) | Control vs CD98hc siRNA | 0.1895 |
| | Control vs JSH-23 | 0.0148 |
| | Control vs Control + CMS | 0.0007 |
| | CD98hc siRNA vs JSH-23 | 0.6009 |
| | Control + CMS vs CD98hc siRNA + CMS | 0.0044 |
| | Control + CMS vs JSH-23 + CMS | 0.0243 |
| | CD98hc siRNA + CMS vs JSH-23 + CMS | 0.2118 |
| IL-8 (pg/ml) | Control vs CD98hc siRNA | 0.1575 |
| | Control vs JSH-23 | 0.0358 |
| | Control vs Control + CMS | 0.0032 |
| | CD98hc siRNA vs JSH-23 | 0.4833 |
| | Control + CMS vs CD98hc siRNA + CMS | <0.0001 |
| | Control + CMS vs JSH-23 + CMS | 0.0037 |
| | CD98hc siRNA + CMS vs JSH-23 + CMS | 0.1104 |

* Unpaired t-test
† Two-way ANOVA

### Figure 5E:

| **TT1** | | **P value** |
|---|---|---|
| IL-6 (pg/ml) | Untreated vs 10% hydrogel | 0.0014 |
| | 10% hydrogel vs 10% hydrogel + JSH-23 | 0.0006 |
| IL-8 (pg/ml) | Untreated vs 10% hydrogel | 0.0044 |
| | 10% hydrogel vs 10% hydrogel + JSH-23 | 0.0003 |

| **A549** | | **P value** |
|---|---|---|
| IL-6 (pg/ml) | Untreated vs 10% hydrogel | 0.0225 |
| | 10% hydrogel vs 10% hydrogel + JSH-23 | 0.0011 |
| IL-8 (pg/ml) | Untreated vs 10% hydrogel | 0.0005 |
| | 10% hydrogel vs 10% hydrogel + JSH-23 | 0.0003 |

* Unpaired t-test
† Two-way ANOVA

### Figure 6A:

| **TT1** | | **P value** |
| --- | --- | --- |
| IL-6 (pg/ml) | Untreated vs LPS | 0.0157 |
| | LPS vs LPS + TRPV4 agonist | 0.0406 |
| | LPS vs LPS + TRPV4 inhibitor | 0.3622 |
| | Untreated vs CMS | 0.0009 |
| | CMS vs EGTA + CMS | 0.0032 |
| | CMS vs TRPV4 agonist + CMS | 0.0003 |
| | CMS vs TRPV4 inhibitor + CMS | 0.0027 |
| IL-8 (pg/ml) | Untreated vs LPS | 0.0012 |
| | LPS vs LPS + TRPV4 agonist | 0.0026 |
| | LPS vs LPS + TRPV4 inhibitor | 0.0441 |
| | Untreated vs CMS | <0.0001 |
| | CMS vs EGTA + CMS | 0.0059 |
| | CMS vs TRPV4 agonist + CMS | 0.0119 |
| | CMS vs TRPV4 inhibitor + CMS | 0.0235 |

* Unpaired t-test
† Two-way ANOVA

### Figure 6A:

| A549 | | P value |
|---|---|---|
| IL-6 (pg/ml) | Untreated vs LPS | 0.0128 |
| | LPS vs LPS + TRPV4 agonist | 0.1042 |
| | LPS vs LPS + TRPV4 inhibitor | 0.0063 |
| | Untreated vs CMS | 0.0004 |
| | CMS vs EGTA + CMS | 0.0072 |
| | CMS vs TRPV4 agonist + CMS | 0.0006 |
| | CMS vs TRPV4 inhibitor + CMS | 0.0039 |
| IL-8 (pg/ml) | Untreated vs LPS | 0.0017 |
| | LPS vs LPS + TRPV4 agonist | 0.0015 |
| | LPS vs LPS + TRPV4 inhibitor | 0.0630 |
| | Untreated vs CMS | 0.0033 |
| | CMS vs EGTA + CMS | 0.0051 |
| | CMS vs TRPV4 agonist + CMS | 0.0319 |
| | CMS vs TRPV4 inhibitor + CMS | 0.0608 |

* Unpaired t-test
† Two-way ANOVA

### Figure 6B:

| | | P value |
|---|---|---|
| Fluorescence (AU) | Control vs TRPV4 agonist | 0.2173 |
| | Control vs CD98hc siRNA | 0.0084 |
| | Control vs EGTA | 0.0024 |
| | Control vs TRPV4 inhibitor | 0.0043 |
| | Control vs NF-KB inhibitor | 0.1640 |
| | Control vs Control + CMS | 0.0006 |
| | Control + CMS vs TRPV4 agonist + CMS | 0.0008 |
| | Control + CMS vs CD98hc siRNA + CMS | <0.0001 |
| | Control + CMS vs EGTA + CMS | 0.0100 |
| | Control + CMS vs TRPV4 inhibitor + CMS | 0.0025 |
| | Control + CMS vs NF-KB inhibitor + CMS | 0.0058 |

* Unpaired t-test
† Two-way ANOVA

### Figure 7A:

| **A549** | | **P value** |
|---|---|---|
| IL-6 (pg/ml) | DMSO + Glycerol/TBS vs DMSO + RBD | 0.0006 |
| | DMSO + RBD vs Dexamethasone + RBD | 0.0014 |
| | DMSO + RBD vs Cynaropicrin + RBD | 0.0457 |
| IL-8 (pg/ml) | DMSO + Glycerol/TBS vs DMSO + RBD | 0.0004 |
| | DMSO + RBD vs Dexamethasone + RBD | 0.0341 |
| | DMSO + RBD vs Cynaropicrin + RBD | 0.0005 |
| TNF-α (pg/ml) | DMSO + Glycerol/TBS vs DMSO + RBD | 0.0059 |
| | DMSO + RBD vs Dexamethasone + RBD | 0.0195 |
| | DMSO + RBD vs Cynaropicrin + RBD | 0.0809 |

* Unpaired t-test
† Two-way ANOVA

### Figure 7A:

| **THP-1** | | **P value** |
|---|---|---|
| IL-6 (pg/ml) | DMSO + Glycerol/TBS vs DMSO + RBD | 0.0330 |
| | DMSO + RBD vs Dexamethasone + RBD | 0.0166 |
| | DMSO + RBD vs Cynaropicrin + RBD | 0.0357 |
| IL-8 (pg/ml) | DMSO + Glycerol/TBS vs DMSO + RBD | 0.0015 |
| | DMSO + RBD vs Dexamethasone + RBD | 0.0468 |
| | DMSO + RBD vs Cynaropicrin + RBD | 0.0106 |
| TNF-α (pg/ml) | DMSO + Glycerol/TBS vs DMSO + RBD | 0.0011 |
| | DMSO + RBD vs Dexamethasone + RBD | 0.0458 |
| | DMSO + RBD vs Cynaropicrin + RBD | 0.2759 |

* Unpaired t-test
† Two-way ANOVA

### Figure 7B:

| A549 | | P value |
|---|---|---|
| IL-6 (pg/ml) | siRNA control vs siRNA control + RBD | 0.0010 |
| | siRNA control + RBD vs CD98hc siRNA + RBD | 0.0003 |
| | siRNA control + RBD vs Galectin-3 siRNA + RBD | 0.0080 |
| IL-8 (pg/ml) | siRNA control vs siRNA control + RBD | 0.0003 |
| | siRNA control + RBD vs CD98hc siRNA + RBD | 0.0155 |
| | siRNA control + RBD vs Galectin-3 siRNA + RBD | 0.0380 |
| TNF-α (pg/ml) | siRNA control vs siRNA control + RBD | 0.1584 |
| | siRNA control + RBD vs CD98hc siRNA + RBD | 0.1180 |
| | siRNA control + RBD vs Galectin-3 siRNA + RBD | 0.1179 |

* Unpaired t-test
† Two-way ANOVA

#### Figure 7B:

| Calu-3 | | P value |
|---|---|---|
| IL-6 (pg/ml) | siRNA control vs siRNA control + RBD | <0.0001 |
| | siRNA control + RBD vs CD98hc siRNA + RBD | 0.0031 |
| | siRNA control + RBD vs Galectin-3 siRNA + RBD | 0.0112 |
| IL-8 (pg/ml) | siRNA control vs siRNA control + RBD | 0.0032 |
| | siRNA control + RBD vs CD98hc siRNA + RBD | 0.0109 |
| | siRNA control + RBD vs Galectin-3 siRNA + RBD | 0.0252 |

#### Figure 7E:

| | | P value |
|---|---|---|
| Number of plaques | siRNA control vs CD98hc siRNA | 0.5909 |
| | siRNA control + Galectin-3 siRNA | 0.3555 |

* Unpaired t-test
† Two-way ANOVA

### Supp Figure 3A:

| **Human IL-6 (pg/ml)** | | **P value** |
|---|---|---|
| LPS + Veh vs Cynaropicrin 5µM + LPS | Day 1 | 0.1370 |
| | Day 2 | 0.0248 |
| | Day 7 | 0.0344 |
| **Human IL-8 (pg/ml)** | | **P value** |
| LPS + Veh vs Cynaropicrin 5µM + LPS | Day 1 | 0.0075 |
| | Day 2 | 0.0022 |
| | Day 7 | 0.0010 |
| **Human TNF-α (pg/ml)** | | **P value** |
| LPS + Veh vs Cynaropicrin 5µM + LPS | Day 1 | 0.2633 |
| | Day 2 | 0.3294 |
| | Day 7 | 0.3192 |

* Unpaired t-test

† Two-way ANOVA

### Supp Figure 3B:

| **Human IL-6 (pg/ml)** | **P value** |
|---|---|
| PBS + Veh vs PBS + Cyn | 0.2572 |
| PBS + Veh vs LPS + Veh | <0.0001 |
| LPS + Veh vs LPS + Cyn | 0.0220 |
| **Human IL-8 (pg/ml)** | **P value** |
| PBS + Veh vs PBS + Cyn | 0.1234 |
| PBS + Veh vs LPS + Veh | 0.0169 |
| LPS + Veh vs LPS + Cyn | 0.0488 |

* Unpaired t-test
† Two-way ANOVA

### Supp Figure 4A:

| **Human IL-6 (pg/ml)** | **P value** |
|---|---|
| Control vs LPS | 0.0257 |
| siRNA control vs CD98hc siRNA | 0.0222 |
| LPS + siRNA control vs LPS + CD98hc siRNA | 0.0491 |
| **Human IL-8 (pg/ml)** | **P value** |
| Control vs LPS | 0.0404 |
| siRNA control vs CD98hc siRNA | 0.0405 |
| LPS + siRNA control vs LPS + CD98hc siRNA | 0.0448 |
| **Human TNF-α (pg/ml)** | **P value** |
| Control vs LPS | 0.0422 |
| siRNA control vs CD98hc siRNA | 0.6111 |
| LPS + siRNA control vs LPS + CD98hc siRNA | 0.9781 |

* Unpaired t-test
† Two-way ANOVA

### Supp Figure 4B:

| **TT1** | |
|---|---|
| **Human IL-6 (pg/ml)** | **P value** |
| CD98hc siRNA vs Cynaropicrin | 0.8236 |
| CD98hc siRNA + LPS vs Cynaropicrin + LPS | 0.7667 |
| **Human IL-8 (pg/ml)** | **P value** |
| CD98hc siRNA vs Cynaropicrin | 0.6126 |
| CD98hc siRNA + LPS vs Cynaropicrin + LPS | 0.6974 |
| **Human TNF-α (pg/ml)** | **P value** |
| CD98hc siRNA vs Cynaropicrin | 0.6111 |
| CD98hc siRNA + LPS vs Cynaropicrin + LPS | 0.9781 |

* Unpaired t-test
† Two-way ANOVA

### Supp Figure 4B:

| **A549** | |
|---|---|
| **Human IL-6 (pg/ml)** | **P value** |
| CD98hc siRNA vs Cynaropicrin | 0.8878 |
| CD98hc siRNA + LPS vs Cynaropicrin + LPS | 0.4778 |
| **Human IL-8 (pg/ml)** | **P value** |
| CD98hc siRNA vs Cynaropicrin | 0.6879 |
| CD98hc siRNA + LPS vs Cynaropicrin + LPS | 0.0713 |
| **Human TNF-α (pg/ml)** | **P value** |
| CD98hc siRNA vs Cynaropicrin | 0.1015 |
| CD98hc siRNA + LPS vs Cynaropicrin + LPS | 0.8721 |

* Unpaired t-test
† Two-way ANOVA

### Supp Figure 4C:

| **TT1, 2 h** | |
|---|---|
| **Human IL-6 (pg/ml)** | **P value** |
| siRNA control vs CD98hc siRNA | 0.1883 |
| siRNA control vs siRNA control + CMS | 0.0790 |
| siRNA control + CMS vs CD98hc siRNA + CMS | 0.0156 |
| **Human IL-8 (pg/ml)** | **P value** |
| siRNA control vs CD98hc siRNA | 0.0065 |
| siRNA control vs siRNA control + CMS | 0.0242 |
| siRNA control + CMS vs CD98hc siRNA + CMS | 0.0172 |
| **Human TNF-α (pg/ml)** | **P value** |
| siRNA control vs CD98hc siRNA | 0.6453 |
| siRNA control vs siRNA control + CMS | 0.0662 |
| siRNA control + CMS vs CD98hc siRNA + CMS | 0.6397 |

* Unpaired t-test
† Two-way ANOVA

### Supp Figure 4C:

| **TT1, 24 h** | |
|---|---|
| **Human IL-6 (pg/ml)** | **P value** |
| siRNA control vs CD98hc siRNA | 0.0345 |
| siRNA control vs siRNA control + CMS | 0.0325 |
| siRNA control + CMS vs CD98hc siRNA + CMS | 0.0832 |
| **Human IL-8 (pg/ml)** | **P value** |
| siRNA control vs CD98hc siRNA | 0.0041 |
| siRNA control vs siRNA control + CMS | 0.0224 |
| siRNA control + CMS vs CD98hc siRNA + CMS | 0.0023 |
| **Human TNF-α (pg/ml)** | **P value** |
| siRNA control vs CD98hc siRNA | 0.1132 |
| siRNA control vs siRNA control + CMS | 0.0151 |
| siRNA control + CMS vs CD98hc siRNA + CMS | 0.2760 |

* Unpaired t-test
† Two-way ANOVA

### Supp Figure 4C:

| A549, 24 h | |
|---|---|
| **Human IL-6 (pg/ml)** | **P value** |
| siRNA control vs CD98hc siRNA | 0.0276 |
| siRNA control vs siRNA control + CMS | 0.0519 |
| siRNA control + CMS vs CD98hc siRNA + CMS | 0.0183 |
| **Human IL-8 (pg/ml)** | **P value** |
| siRNA control vs CD98hc siRNA | 0.1697 |
| siRNA control vs siRNA control + CMS | 0.0177 |
| siRNA control + CMS vs CD98hc siRNA + CMS | 0.0356 |
| **Human TNF-α (pg/ml)** | **P value** |
| siRNA control vs CD98hc siRNA | 0.1132 |
| siRNA control vs siRNA control + CMS | 0.0217 |
| siRNA control + CMS vs CD98hc siRNA + CMS | 0.2760 |

* Unpaired t-test
† Two-way ANOVA

### Supp Figure 4D:

| **TT1** | |
|---|---|
| **Human IL-6 (pg/ml)** | **P value** |
| CD98hc siRNA vs Cynaropicrin | 0.6963 |
| CD98hc siRNA + CMS vs Cynaropicrin + CMS | 0.2020 |
| **Human IL-8 (pg/ml)** | **P value** |
| CD98hc siRNA vs Cynaropicrin | 0.6229 |
| CD98hc siRNA + CMS vs Cynaropicrin + CMS | 0.1411 |

| **A549** | |
|---|---|
| **Human IL-6 (pg/ml)** | **P value** |
| CD98hc siRNA vs Cynaropicrin | 0.6887 |
| CD98hc siRNA + CMS vs Cynaropicrin + CMS | 0.4316 |
| **Human IL-8 (pg/ml)** | **P value** |
| CD98hc siRNA vs Cynaropicrin | 0.4240 |
| CD98hc siRNA + CMS vs Cynaropicrin + CMS | 0.5826 |

* Unpaired t-test
† Two-way ANOVA

### Supp Figure 4E:

| **TT1** | |
|---|---|
| **Human IL-6 (pg/ml)** | **P value** |
| 3% hydrogel vs 10% hydrogel | 0.0197 |
| **Human IL-8 (pg/ml)** | **P value** |
| 3% hydrogel vs 10% hydrogel | 0.0010 |

| **A549** | |
|---|---|
| **Human IL-6 (pg/ml)** | **P value** |
| 3% hydrogel vs 10% hydrogel | 0.0342 |
| **Human IL-8 (pg/ml)** | **P value** |
| 3% hydrogel vs 10% hydrogel | 0.0251 |

* Unpaired t-test
† Two-way ANOVA

### Supp Figure 4F:

| TT1 | |
|---|---|
| **Human IL-6 (pg/ml)** | **P value** |
| LPS vs LPS + 10% hydrogel | 0.0551 |
| LPS vs LPS + CMS | 0.0970 |
| LPS + 10% hydrogel vs LPS + 10% hydrogel + CMS | 0.0450 |
| LPS + CMS vs LPS + 10% hydrogel + CMS | 0.0167 |
| **Human IL-8 (pg/ml)** | **P value** |
| LPS vs LPS + 10% hydrogel | 0.0117 |
| LPS vs LPS + CMS | 0.0096 |
| LPS + 10% hydrogel vs LPS + 10% hydrogel + CMS | 0.0545 |
| LPS + CMS vs LPS + 10% hydrogel + CMS | 0.0255 |

* Unpaired t-test
† Two-way ANOVA

### Supp Figure 4F:

| **A549** | |
|---|---|
| **Human IL-6 (pg/ml)** | **P value** |
| LPS vs LPS + 10% hydrogel | 0.0518 |
| LPS vs LPS + CMS | 0.1529 |
| LPS + 10% hydrogel vs LPS + 10% hydrogel + CMS | 0.0650 |
| LPS + CMS vs LPS + 10% hydrogel + CMS | 0.1996 |
| **Human IL-8 (pg/ml)** | **P value** |
| LPS vs LPS + 10% hydrogel | 0.0224 |
| LPS vs LPS + CMS | 0.0156 |
| LPS + 10% hydrogel vs LPS + 10% hydrogel + CMS | 0.0469 |
| LPS + CMS vs LPS + 10% hydrogel + CMS | 0.1136 |

* Unpaired t-test
† Two-way ANOVA
